# Not out of the box: phylogeny of the broadly sampled Buxaceae

**DOI:** 10.1101/2020.08.03.235267

**Authors:** Alexey Shipunov, Hye Ji Lee, Jinhee Choi, Kyle Pay, Sarah DeSpiegelaire, Aaron Floden

## Abstract

The Buxaceae constitute a morphologically diverse phylogenetic lineage of six genera, which includes about 140 species. The most well-known genera are *Buxus*, *Sarcococca*, and *Pachysandra.* Few species of woody *Styloceras* grow on mid-elevations in the Andes mountains region. *Didymeles*, with three species endemic to Madagascar, and the monotypic *Haptanthus* from Honduras, are the most unusual members of the group. The infra-familial classification of Buxaceae is controversial, and molecular data about many species, especially Old World, is still lacking. We used broad taxonomic sampling and molecular data from four chloroplast markers, and the nuclear ribosomal ITS to estimate their phylogeny. These data provide phylogenetic placements of 50 species and enabled better estimates of boundaries in Buxaceae. We described two subfamilies, two monotypic genera, two *Buxus* subgenera, and one new species of *Didymeles* from Madagascar.

## Introduction

Within flowering plants, the boxwood family, Buxaceae Dumort. (Dumortier, 1822), together with *Haptanthus* A. Goldberg & C. Nelson (Goldberg & Nelson, 1989) and *Didymeles* Thouars (Thouars, 1804) form an old (Takahashi et al., 2017), distinct (Worberg et al., 2007; Gutiérrez, 2014), and diverse (about 140 species) taxon. They are distributed almost worldwide (Fig. 1) with high species diversity in Tropical America, Southern Africa / Madagascar, and East Asia (Jarvis, 1989; Köhler, 2004; Köhler, 2007; Köhler, 2009). They have no close relatives; the nearest branch on the phylogenetic tree of angiosperms are East Asian *Trochodendron* and *Tetracentron*, to which they were not even considered to be related before the “molecular era” (Castilho et al., 1999; Takhtajan, 2009).

**Figure 1.**
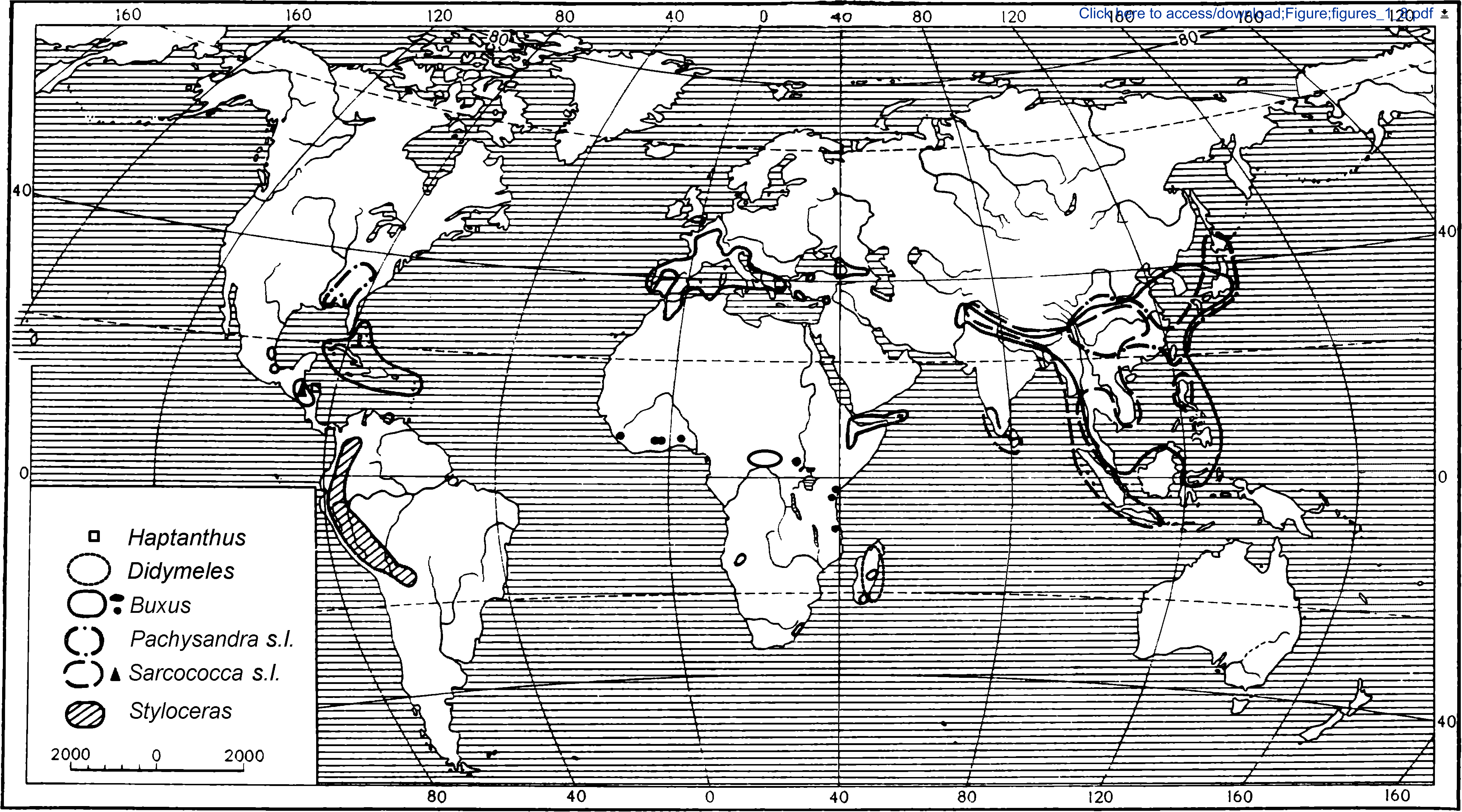
The map of the world which shows the geographic distribution of the group.

There are multiple morphological characters which unite the group (Oskolski & al., 2015), for example, cyclocytic or laterocytic stomata, frequently triplinerved leaf venation, racemose inflorescences, frequently dimerous flowers, stamen-sepalum complex in many representatives, and unusual pregnan steroidal alkaloids (Hardman, 1987; von Balthazar & Endress, 2002a, b; von Balthazar et al., 2003; Köhler, 2007). Two anomalous genera of the group have been variously treated as independent families or as part of the Buxaceae. *Haptanthus* was discovered in Atlantida province of Honduras (Goldberg & Nelson, 1989) and segregated as a monotypic family based on its perianth structure (Nelson, 2001; Shipunov, 2003), but the actual taxonomic position has been heavily deliberated (Doust & Stevens, 2005; Goldberg & Alden, 2005). As a result, it was accepted as the only *incertae sedis* genus of flowering plants by Takhtajan (2009) and even mentioned as such in the example 1 of ICN Article 3.1 (“Ex.1. The genus *Haptanthus* Goldberg & C. Nelson (in Syst. Bot. 14: 16. 1989) was originally described without being assigned to a family…”). The *locus classicus* was apparently lost through deforestation, but *Haptanthus* was recently re-discovered (Shipunov & Shipunova, 2010) and also found in more locations. Since then, it has been established *ex situ* in the Lancetilla Botanical Garden, Honduras (Bejarano, 2015). Both morphologically and molecularly, *Haptanthus* has similarities with the “core Buxaceae” as well as with *Didymeles* (Shipunov & Shipunova, 2010; Oskolski & al., 2015).

Geographically, *Didymeles* is close to the South African / Madagascan center of the group diversity, but until the “molecular era”, it was not considered as a relative to the core Buxaceae (Worberg et al., 2007; Takhtajan, 2009). There are three (Köhler, 2007) or two (Madagascar Catalogue, 2018) accepted species but also some unidentified and putatively new material (Madagascar Catalogue, 2018). One of the most striking characters of *Didymeles* is the presence of a monomerous gynoecium (Leandri, 1937).

Among the core genera of the Buxaceae, *Buxus* (Linnaeus, 1753) is the most speciose (ca. 110 species) and well-known (Larson, 1996) group. The infrageneric taxonomy of boxwoods is highly correlated with geography, and American and African species were at first considered as separate genera: the American *Tricera* (Schreber, 1797) and the African *Notobuxus* (Oliver, 1882). Van Tieghem (1897) split African *Buxus* in three genera, *Buxanthus* Tiegh., *Buxella* Tiegh., and *Notobuxus*. Later taxonomic revisions treated these as sections or subgenera, e.g., sect. *Tricera* (Schreb.) Baill. and subg. *Probuxus* Mathou (Baillon, 1859; see also Mathou, 1940). Though *Buxus* is widely distributed and highly diverse in comparison to other genera in the family, the apparently rapid speciation of *Buxus* on Cuba, which has almost 40 endemic boxwood species (Sauget & Liogier, 1974; Köhler, 1998; Köhler, 2004; Köhler, 2006; Gutiérrez, 2014; Köhler, 2014), is remarkable relative to the diversity elsewhere. Despite their overall similarities, there are some substantial differences between these subgroups. For example, the Old World *Buxus* typically have cortical vascular bundles in each angle of the branchlets, accompanied by fiber strands in the Eurasian taxa (Köhler, 2004; Köhler, 2007), whereas both these characters are absent in New World *Buxus*.

The subfamily *Pachysandroideae* Record & Garratt (or tribe *Pachysandreae* Reveal: Record & Garratt, 1925; Reveal, 2011, 2012) is comprised of three genera: *Pachysandra* Michx., *Sarcococca* Lindl., and *Styloceras* Kunth. The three species of *Pachysandra* (Michaux, 1803) are herbaceous or suffruticose rhizomatous plants (Robbins, 1968), and their disjunct distributions are classic examples of the East Asian — Eastern North American floristic disjunction (Gray, 1846; Raven & Axelrod, 1974). *Pachysandra procumbens* Michx. (“Allegheny spurge”), from North America, and *P. terminalis* Siebold & Zucc. (“kichijiso”) from Japan and China, are common garden plants used extensively as ground covers (Batchelor & Miyabe, 1893; Dirr & Alexander, 1979; Channell & Wood, 1987). *Pachysandra terminalis* differs significantly from the two other species in its terminal inflorescences, strongly spreading rhizomatous habitat, bicarpellate gynoecium, and white baccate fruit. *Pachysandra procumbens* and the Chinese polymorphic *P. axillaris* Franchet have been shown to be sister species in molecular phylogenetic analyses (Jiao & Li, 2009) despite their different appearance and the geographic proximity of the latter to *P. terminalis*.

The species of *Sarcococca* (Lindley, 1826) are small shrubs and understory plants of humid lowland and mountain forests, mostly in southeastern Asia. With their evergreen habit, winter-flowering, and fragrant flowers, the species of *Sarococca* are generally referred to as “Sweet Box” (Sealy, 1986). Sealy’s (1986) revision was a first step in resolving some of the many taxonomic problems and specific boundaries in *Sarcococca*. Despite his efforts, regional floristic treatments do not recognize the same suite of taxa or the same synonymy (Backer & Bakhuizen van den Brink, 1965; Min & Brückner, 2008) and there is extensive variation in morphology across the distributions of some species. *Sarcococca* is also remarkable with the apparent geographic disjunction of *S. conzattii* (Standl.) I.M. Johnst., which occurs in southern Mexico (Oaxaca) and Guatemala (Johnston, 1939). It was first described as a *Buxus* (Standley, 1936; Johnston, 1938), but Sealy (1986) doubted it belonged there or with *Sarcococca* based on its morphology, inflorescence structure, and fruit type.

The Andean *Styloceras* (Kunth, 1824) are small trees or shrubs, and with the exception of *Buxus citrifolia* (Willd.) Spreng. are the only representatives of Buxaceae in South America. *Styloceras* has been treated as a monogeneric family, the Stylocerataceae Takht. ex Reveal & Hoogland (Reveal & Hoogland, 1990), though it is now shown to be embedded within the Buxaceae (von Balthazar & al., 2000). Six species are currently recognized (Gentry & Aymard, 1993; Torrez & Jorgensen, 2010; Ulloa Ulloa & al., 2017), but we believe that the full diversity of this rare group is not yet understood.

The infra-familial classification of Buxaceae is still controversial. Mathou (1940) used two tribes, *Buxeae* Dumort. (with *Buxus* only) and *Pachysandreae* (with the other three genera of core Buxaceae), whereas Takhtajan (2009) essentially raised each genus into its own tribe. Reveal (2012) used two subfamilies, *Pachysandroideae* and *Buxoideae* Beilschm. (Beilschmid, 1833). The recent work on *Buxus* Caribbean taxa (Gutiérrez, 2014) provided a starting point for understanding the evolution of the significant portion of American boxwood species and group as a whole, but data about many other species, especially Old World, is still lacking. There is no recent synthetic classification of the family.

We attempt to provide a comprehensive classification scheme for Buxaceae through high taxonomic sampling that will serve as a framework for future studies in the group.

## Materials and Methods

Our sampling protocol, due to the broad geographic distribution of the Buxaceae and rarity of many taxa, used herbarium material wherever possible for DNA extractions. We extracted DNA from 286 samples: 271 samples from herbaria and 15 from our collections. Herbarium tissue samples were obtained from numerous herbaria (B, BO, BRIT, CAS, F, HUH, IBSC, JEPS, NBG, NY, PE, PRE, SAM, SP, SPF, TI, US, and USM) with the kind permission of the herbarium curators. All vouchers were photographed so that the DNA sequence data could be linked to imaged specimens. Preference for a sample was always given to vouchers annotated by regional or generic experts. Besides, we supplemented missing data with 175 sequences from GenBank and Barcode of Life Data System (with necessary precautions: Funk & al., 2018) of species or fragments which complemented our database. In total, we sampled all genera and 128 of 140 species (91%) in Buxaceae. Only some rare and local species which are underrepresented in herbaria and living collections, were not sampled in our DNA dataset.

We used standard approaches for DNA extraction and employed commercial DNA extraction kits. DNA was extracted using either a MO BIO PowerPlant DNA Isolation Kit (MO BIO Laboratories, Carlsbad, California, USA) or NUCLEOSPIN Plant II Kit (MACHEREY-NAGEL GmbH & Co. KG, Düren, Germany). Dry plant leaf material (typically, 0.05–0.09 g) was powdered using a sterile mortar and pestle and then processed in accordance with the supplied protocol. We increased the lysis time to 30 minutes and used thermomixer on the slow rotation speed (350 rpm) instead of a water bath. Nanodrop 1000 Spectrophotometer (Thermo Scientific, Wilmington, DE, USA) was used to assess the concentration and purity (the 260/280 nm ratio of absorbance) of DNA samples. In our phylogenetic trees, we decided to integrate data from “barcoding” markers: plastid *rbc*L plus *trn*L-F spacer and nuclear ITS to represent both chloroplast and nuclear genomes. Fortunately, herbarium specimens of Buxaceae typically retain DNA of relatively good quality for many years (Choi & al., 2015). If the particular sample did not yield a sequence of good quality, we tried to use another sample of the same species.

We sequenced the markers mentioned above using primers and protocols in accordance with recommendations of the Barcoding of Life Consortium (Kuzmina & Ivanova, 2011). PCR was carried out as follows: the reaction mixture in a total volume of 20 μL contained 5.2 μL of PCR Master Mix (components from QIAGEN Corporation, Germantown, Maryland, and Thermo Fisher Scientific, Waltham, Massachusetts for Platinum DNA Taq Polymerase), 1 μL of 10 μM forward and reverse primers, 2 μL of DNA solution from the extraction above and 10.8 μL of either MQ purified water (obtained from a Barnstead GenPure Pro system, Thermo Scientific, Langenselbold, Germany), or the TBT-PAR water mix (Samarakoon & al., 2013). Samples were incubated in a thermal cycler: 94° for 5 min, then 35 cycles of 94° for 1 min; 51° (or similar, annealing temperature was varied with a primer) for 1 min, 72° for 2 min, and finally 72° for 10 min. Single-band PCR products were sent for purification and sequencing to Functional Biosciences, Inc. (Madison, Wisconsin) and sequenced there in accordance with standard protocol. Sequences were obtained, assembled, and edited using Sequencher™ 4.5 (Genes Codes Corporation, Ann Arbor, Michigan, USA) and then aligned with AliView (Larsson, 2014) and MUSCLE (Edgar, 2004).

For all procedures and statistic calculations, the R environment (R Core Team, 2019) was used. We used Ripeline (Shipunov, 2020), the R-based DNA sequence analysis pipeline for databasing, sequence analysis, and phylogeny estimation. Ripeline is the combination of UNIX shell scripts and R scripts which allows for (a) species name checks using taxonomy database, (b) cross-validation of sequences, (c) updates from GenBank, (d) completeness analysis and species accumulation control, (e) deselection and replacement of outliers (both on the level of sequences and on the level of trees), (f) sequence alignments using the external tools, (g) flank cleaning, (h) gap coding based on Borchsenius (2009) algorithm which uses simple gap coding *sensu* Simmons and Ochotorena (2000), (i) smart (strict and semi-strict) concatenation (supermatrix production), and (j) a wide variety of phylogenetic outputs, from the k-mer alignment-free to Bayesian and maximal likelihood analyses. In addition, Ripeline is capable of using morphological characters, perform nearest neighbor imputation of missed sequences, and producing super-alignments (Ashkenazy & al., 2018). The Ripeline is available from the primary author’s Github: https://github.com/ashipunov/Ripeline. Within Ripeline, model testing and phylogenetic trees were made with APE (Paradis & al., 2004), MrBayes (Ronquist & Huelsenbeck, 2003), ips (Heibl, 2008), phangorn (Schliep, 2011) and RAxML (Stamatakis, 2014).

Before the alignment, sequence sets were constructed with the principle that sequences produced for this study had priority, and external data were added only to fill sampling gaps or to replace sequences of unreliable quality. This reduced any possible discrepancies based on incorrect identification and absence of proper vouchers in public databases (Funk & al., 2018). Phylogenetic tree construction within Ripeline used both individual markers and their combinations (supermatrices, concatenated sequences). The preference was given to the concatenation of two sequences that originated from one (our) sample (strict concatenation). On the next (semi-strict) stage, sequences with the same species name that were not derived from the same sample were concatenated.

Using Ripeline, we were able to obtain maximum likelihood (ML), Bayesian (MB), and maximum parsimony (MP) phylogenetic trees. Maximum likelihood analyses were run RAxML (Stamatakis, 2014) with 10,000 bootstrap replicates and R ips package (Heibl, 2008). We used a GTR+G+I model based on model testing with R phangorn package (Schliep, 2011). Bayesian analyses were run through the combination of MrBayes 3.2.6 (Ronquist & Huelsenbeck, 2003), and R shipunov package (Shipunov, 2019). MCMC chains were run for 1,000,000 generations, sampling every 10th generation resulting in 100,000 trees. The first 25% of trees were discarded as burn-in, and the remaining trees were summed to calculate the posterior probabilities. The convergence of Bayesian analyses was controlled using the standard deviation of frequencies across runs, and the potential scale reduction factor, PSRF (Ronquist & Huelsenbeck, 2003). Maximum parsimony analyses were run with the help of R phangorn package (Schliep, 2011) using parsimony ratchet (Nixon, 1999) with 2000 iterations and then 1000 bootstrap replicates. With the aid of the R ape package (Paradis & al., 2004), trees were rooted with *Trochodendron aralioides* and *Tetracentron sinense* as outgroups. To assess the congruence between chloroplast and nuclear data, we used the CADM test (Campbell & al., 2011).

We used two kinds of supermatrices. The first was based on plastome (hereafter, CP) sequences only and included four chloroplast regions: *rbc*L, *trn*L-F, *matK* and *petD*. The second supermatrix (hereafter, OI) included two chloroplast sequences (*rbc*L and *trn*L-F) and also ITS2. Our plastome (CP) dataset was longer (5631 bp including 589, 1303, 2614, and 1125 bp of *rbc*L, *trn*L-F, *matK*, and *petD* parts, respectively) but covered less taxonomic diversity: all genera and 72 species (51%). This matrix, therefore, follows the “more genes” approach (Rokas & Carroll, 2005). The second matrix (OI), was shorter (2581 bp including 689 bp ITS) and was generated mostly from our data that covered all genera and 128 species (91%) of the Buxaceae. To help with *Pachysandroideae* phylogeny estimation, we produced the third matrix (hereafter, “full ITS”), which uses full ITS sequences (generally, ITS1, 5.8S, and ITS2). This matrix covered five genera and 29 species of Buxaceae and was 1371 bp in length.

Datasets, scripts, and phylogenic trees used in the preparation of this publication are available from the first author’s Open Repository here: http://ashipunov.info/shipunov/open/buxineae.zip. We encourage readers to reproduce our results and develop our methods further. All sequences were deposited into the GenBank (Support Table 3).

In the paper, we followed the “appropriate citation of taxonomy” (ACT) principle (Seifert & al., 2008) and cited names of the most supra-species groups (Reveal, 2011, 2012) plus those species which are separately discussed.

## Results

In total, we obtained 359 sequences from 118 species (Support Table 3) and sequenced 50 species of Buxaceae for the first time. Of the resulting matrices that were analyzed, the average percentage of the data produced for this study (*versus* data which came from public databases) was near 74%.

CADM test for the congruence between *rbc*L + *trn*L and ITS parts of OI supermatrix returned the average Kendall concordance value (W=0.53576877), the null hypothesis of incongruence was rejected with p-value 0.01598402 (Chi-squared=11341.15325383). In contrast to Rossello et al. (2007), we did not find issues with multiple copies of ITS present in samples.

In essence, MB, ML, and MP analyses resulted in very similar trees, and overall phylogeny is almost identical in MB and ML (Figs. 2–3). Below, we describe our results based on MB analyses of CP and OI supermatrices (Figs. 4–5), and ML analysis of “full ITS” matrix (Fig. 6). All trees are accessible from the open repository.

**Figure 2.**
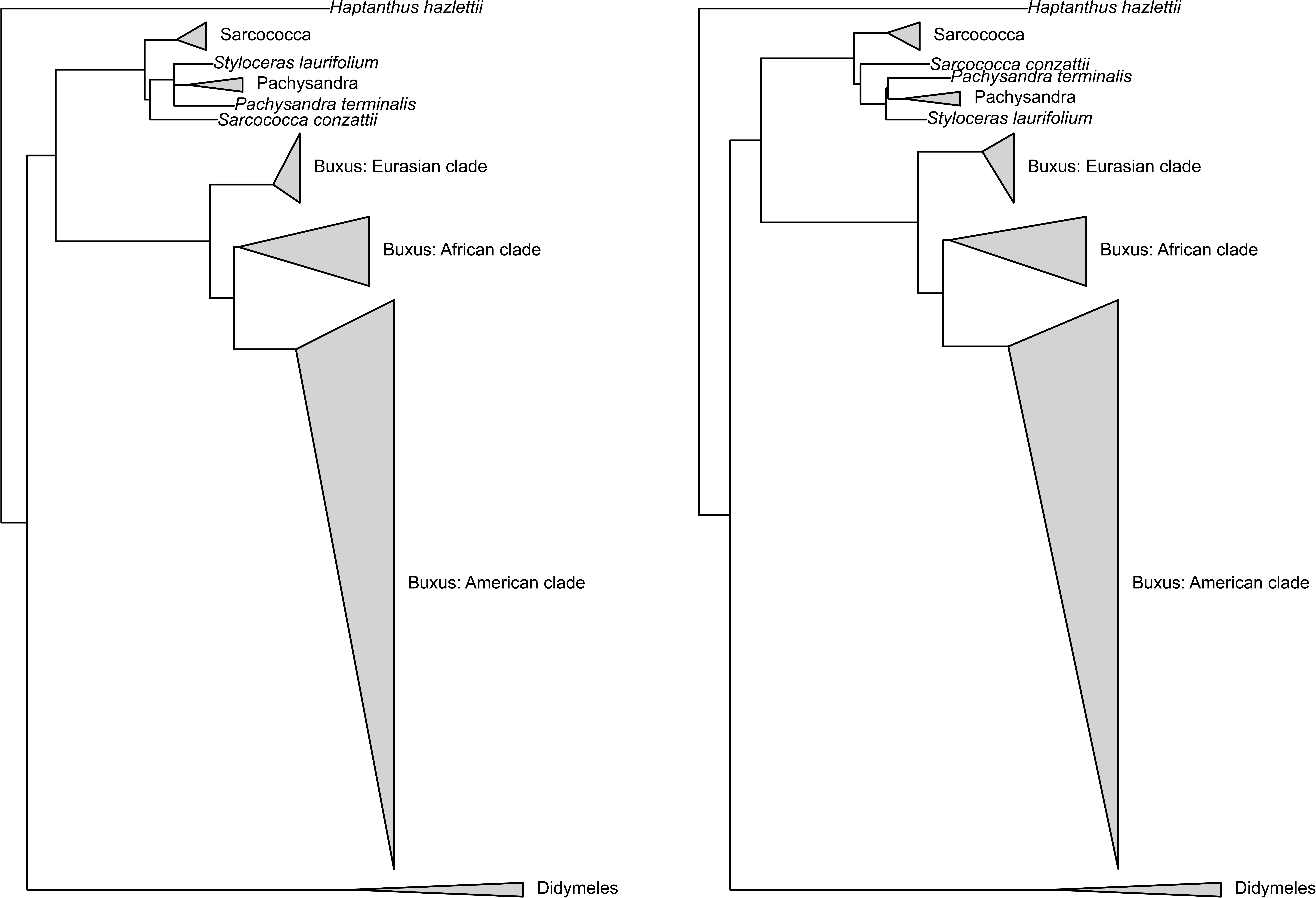
Comparison of the topologies resulted from the phylogenetic analyses of CP (plastid markers) supermatrix: Bayesian (left) and RAxML (right). Each triangle is the result of concatenation applied to the branches of the corresponding phylogenetic trees.

**Figure 3.**
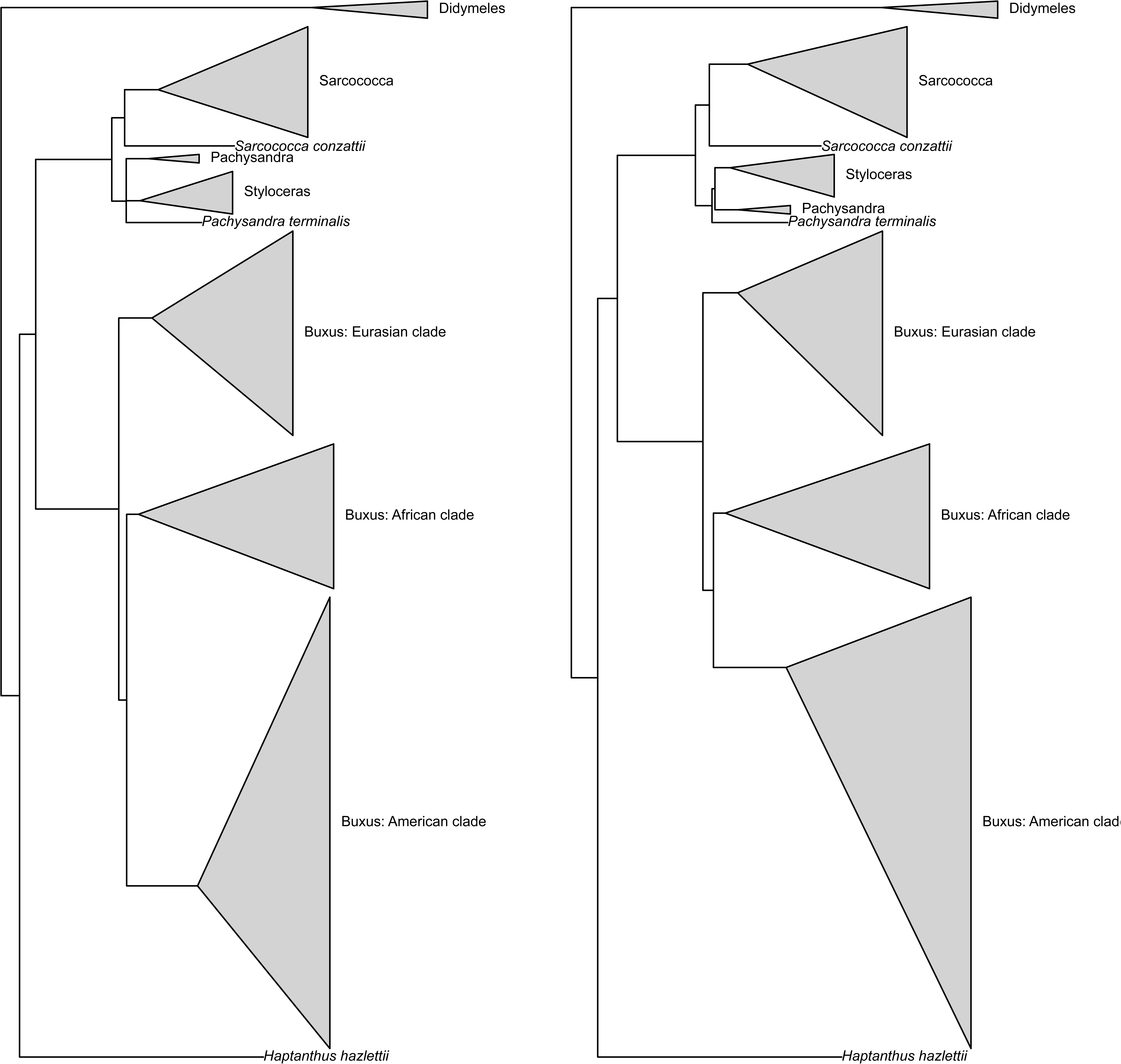
Comparison of the topologies resulted from the phylogenetic analyses of OI (*trn*L, *rbc*L and ITS2) supermatrix: Bayesian (left) and RAxML (right). Each triangle is the result of concatenation applied to the branches of the corresponding phylogenetic trees.

**Figure 4.**
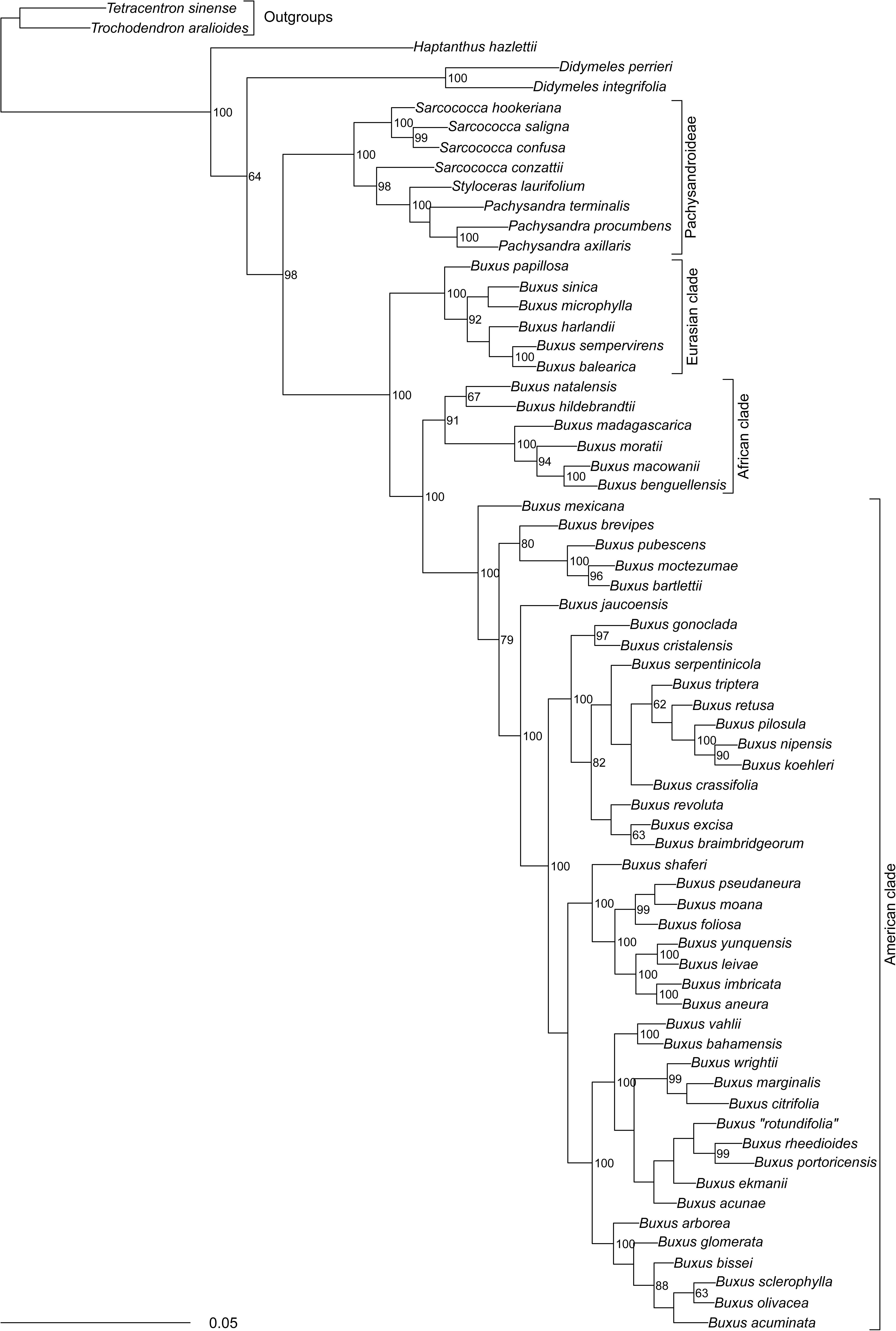
The Bayesian phylogenetic tree obtained from CP (plastid markers) supermatrix, node labels denote the BPP (in %). See the text for the explanation of clade names.

**Figure 5.**
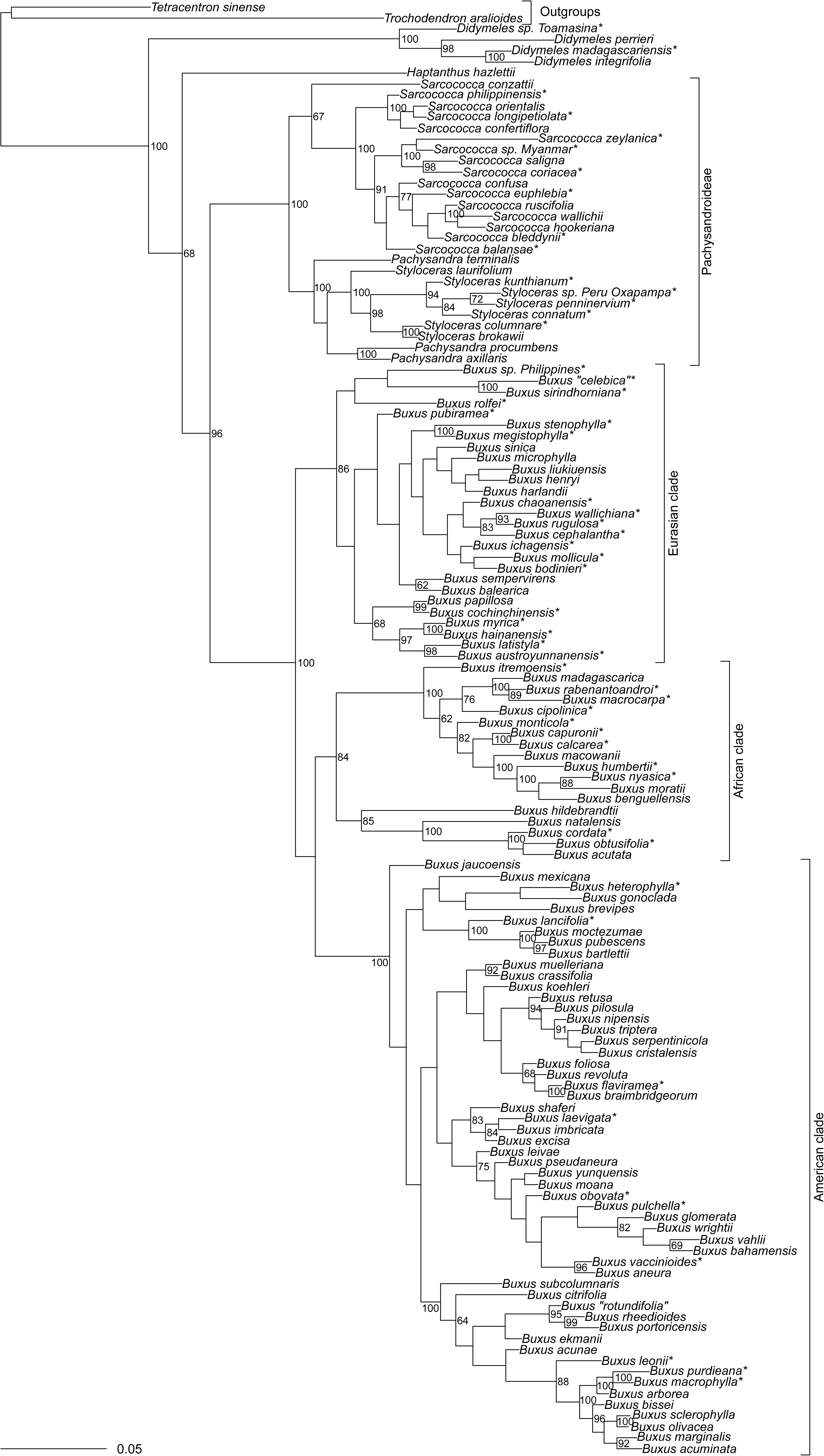
The Bayesian phylogenetic tree obtained from OI (*trn*L, *rbc*L and ITS2) supermatrix, node labels denote the BPP (in %), stars* designate species sequenced for the first time. See the text for the explanation of clade names.

**Figure 6.**
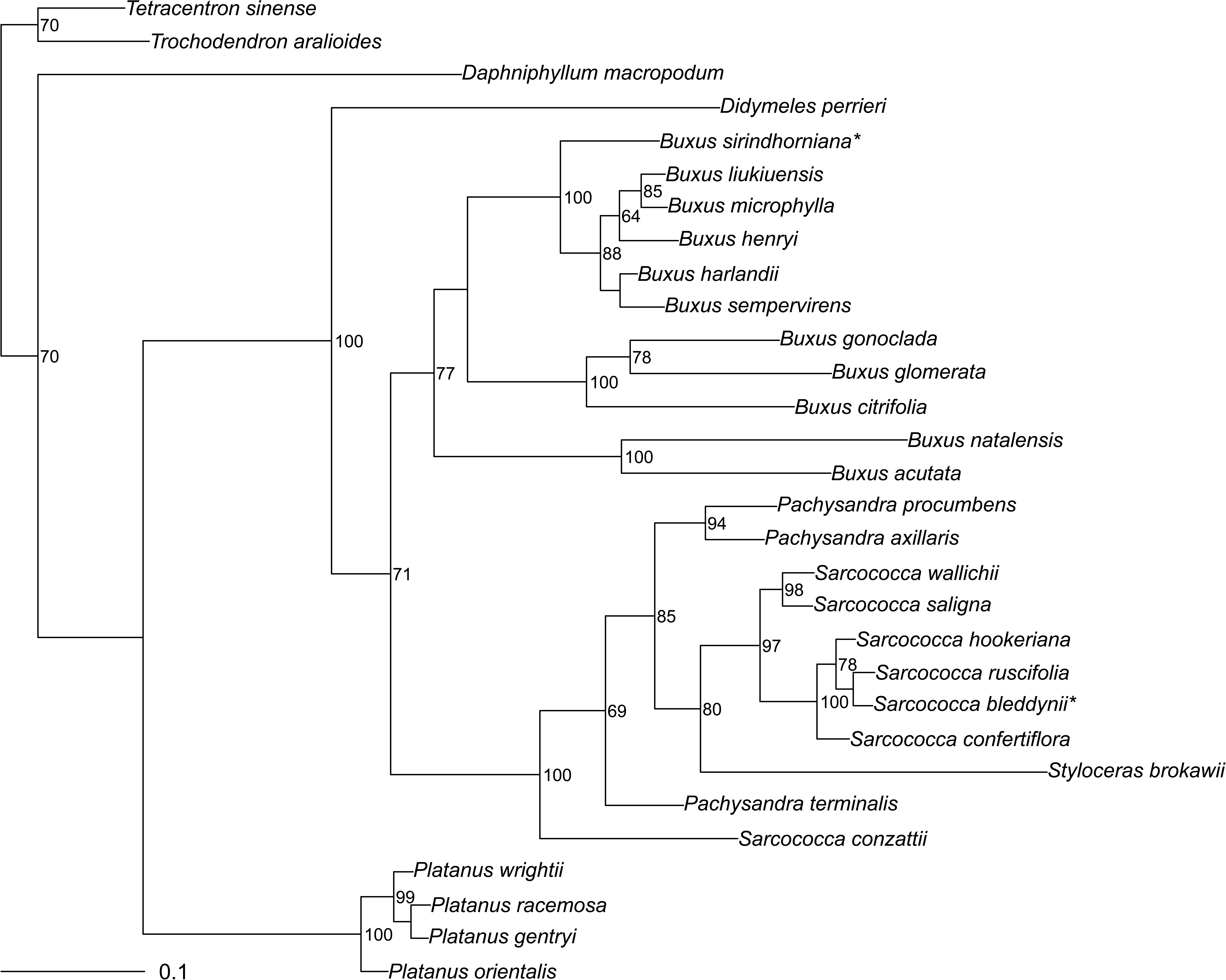
The maximum likelihood (RAxML) tree obtained from the “full ITS” matrix, node labels denote the bootstrap support, stars* designate species sequenced for the first time.

### Didymeles **and** Haptanthus

*Didymeles* and *Haptanthus* were consistently recovered as the two first branches (Figs. 2–5). The Buxaceae as a whole, as well as a node next to *Didymeles* + *Haptanthus* grade, was supported well (BPP > 96%) on CP and OI trees. A morphologically distinct (Fig. 7) sample of *Didymeles* from Toamasina (Madagascar) recovered as a first branch in the *Didymeles* clade (Fig. 4–5).

**Figure 7.**
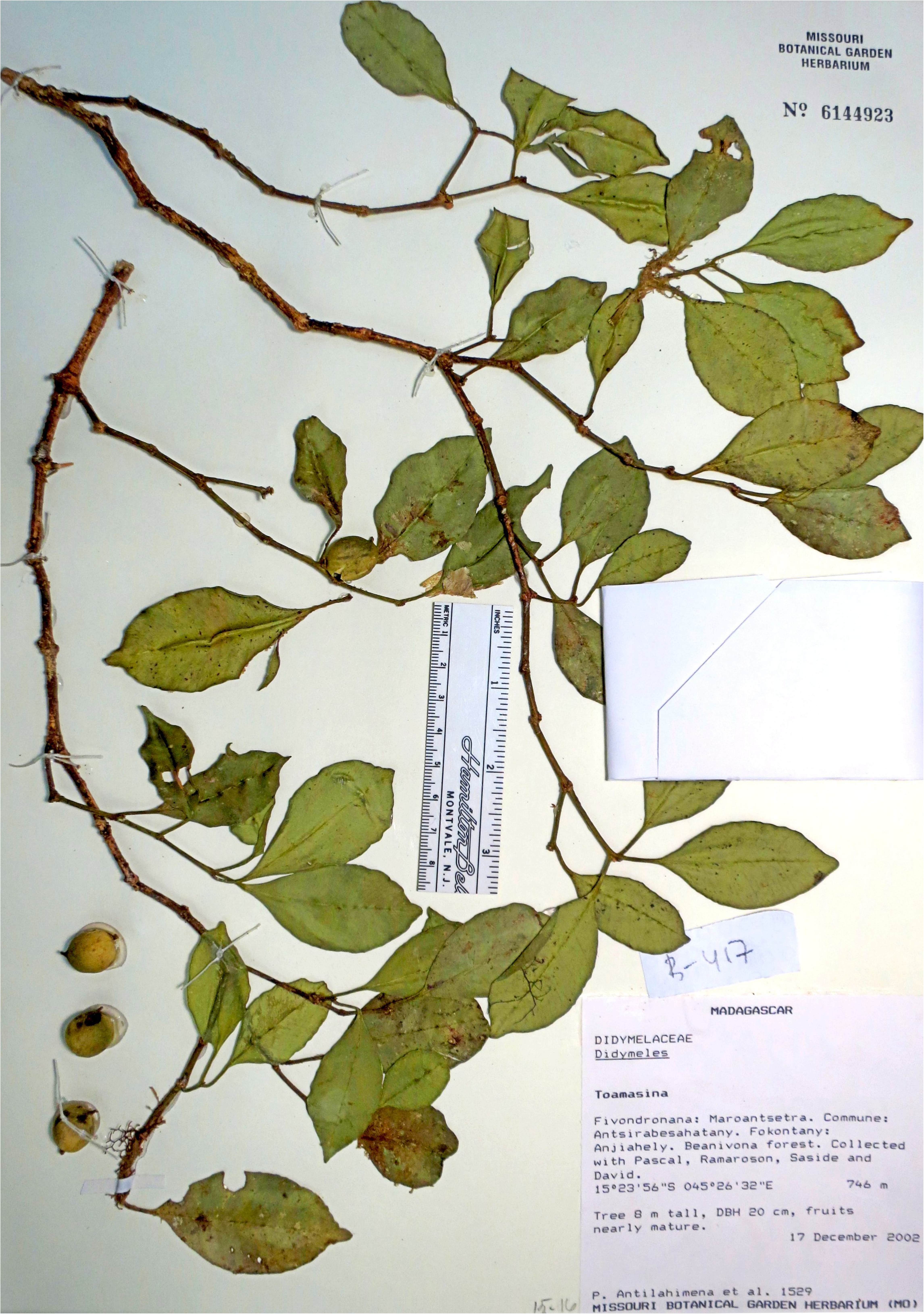
*Didymeles toamasinae*, the holotype (MO).

### Buxus

*Buxus* s.l. is recovered as monophyletic with high support (Figs. 2–5). We sequenced 38 species of *Buxus* (35%) for the first time. Within *Buxus*, three clades were recovered with as monophyletic (Figs. 4–5): an African clade with BPP 84–91% across analyses, an American clade typically highly (BPP 100%) supported, and a Eurasian clade with BPP 86–100%.

The Eurasian clade of *Buxus* is not resolved well. Several South Asian species formed a basal subclade with low support (Fig. 5). This clade includes *Buxus sirindhorniana* W.K. Soh et al. from Thailand, *B. rolfei* S.Vidal and an unidentified sample from the Philippines, and the sample of unknown origin labeled in IBSC as “*Buxus celebica* Hats.” Two other stable Asian subclades recovered (Fig. 5) in OI analyses, one consisted of *B. myrica* H.Lév., *B*. *hainanensis* Merr., *B*. *latistyla* Gagnep., and *B*. *austroyunnanensis* Hatus.; and another which was comprised of *B. stenophylla* Hance and *B. megistophylla* H.Lév.

Within the African *Buxus* clade (Figs. 4–5), species which were sometimes described under *Notobuxus* (like *Buxus cordata* (Radcl.-Sm.) Friis) were present together with species from *Buxus* s.str. (like *B. hildebrandtii* Baill. and *B. madagascarica* Baill.). Support of smaller clades varied but BPP typically was above 50%.

The American clade was the most speciose in our datasets. The first branches form a grade, which includes *B. bartlettii* Standl., *B*. *pubescens* Greenm., *B*. *moctezumae* Eg.Köhler, R.Fernald & Zamudio and *B. mexicana* Brandegee. *Buxus jaucoensis* Eg.Köhler does not hold a stable position on our trees; it is frequently sister to most or all subclades. The next branches were mostly Cuban species (for example, *Buxus koehleri* P.A.González & Borsch, *B. flaviramea* (Britton) Mathou, *B. obovata* Urb., *B. vaccinioides* (Britton) Urb.). The last big subclade includes many non-Cuban species, for example, *Buxus portoricensis* Alain and *B. citrifolia* (Figs. 4–5). Some Cuban species (for example, *Buxus* “*rotundifolia*”, see below about this name) also branch here.

### *Pachysandroideae* Record et Garratt

The results of all phylogenetic analyses of the matrices recovered a monophyletic Pachysandroideae (Figs. 2–6). *Sarcococca* was not monophyletic on CP and “full ITS” trees, and the New World *S. conzattii* was either robustly placed as sister to the other genera of the subfamily (“full ITS”), as sister to *Styloceras* + *Pachysandra* (CP) with high support, or (OI) as sister to the remaining species of *Sarcococca* (with low support).

Seven species of *Sarcococca* were sequenced for the first time (Fig. 5). There was high support (OI) for *S. confertifolia* Sealy, *S*. *longipetiolata* M. Cheng, *S. orientalis* C.Y. Wu, and *S. philippinensis* Stapf ex Sealy group. This clade was the first branch in *Sarcococca* s.str., and the remainder of the *Sarcococca* species formed another stable clade. This last clade includes the recently described Vietnamese *S. bleddynii* J.M.H.Shaw & N. van Du (Shaw, 2011), *S. euphlebia* Merrill from Hainan, unidentified *Sarcococca* sp. from Myanmar as a sister group to *S. zeylanica* Baill., and seven other species.

*Pachysandra* was not recovered as monophyletic in all analyses. Whereas a monophyletic *Pachysandra* presents in CP analyses with low support (Fig. 4), it was not monophyletic on OI and “full ITS” trees. On OI trees, *Pachysandra terminalis* was recovered as sister to *Styloceras* with high support (Fig. 5), and on “full ITS” trees, it was placed between *Sarcococca conzattii* and the remainder of the *Pachysandroideae*, albeit with low support (Fig. 6).

Four (out of six) species of *Styloceras* were sequenced for the first time. On OI trees, *Styloceras* is monophyletic and well-supported (Fig. 5). *Styloceras laurifolium* (Willd.) Kunth is sister to five other species of the genus with the reliable support; two other clades consisted of *S. kunthianum* A. Juss., *S. penninervium* A.H. Gentry & G.A. Aymard and *S. connatum* Torrez & P. Jørg. (BPP 94%) and *S. columnare* Müll.Arg. + *S. brokawii*. H.Gentry & R.B.Foster (BPP 100%). An unidentified sample of *Styloceras* from Oxapampa (Peru) is morphologically similar to *S. penninervium* and was placed sister to it on OI trees (Fig. 5).

## Discussion

Our dataset provides the most broadly sampled phylogenetic analyses of Buxaceae to date. In some groups, the amount of molecular information is doubled, and even tripled (*Styloceras*). Our molecular phylogenetic results support the elevation of *Sarcococca conzattii* into a new genus, *Sealya* (described below).

### Didymeles **and** Haptanthus

Buxaceae *sensu lato* is robustly supported in all our analyses (Figs. 2–5). *Didymeles* and *Haptanthus* form two earliest lineages sister to the remainder of Buxaceae. The analyses resulted in an equivocal placement of *Haptanthus* and *Didymeles*, which is in agreement with the earlier analyses (Shipunov & Shipunova, 2011). This instability *versus* the stability of Buxaceae *sensu lato* provides an indirect support of the integrity of the whole group. As both our results and the morphology of these two distinct genera support a close relationship to the core Buxaceae (Oskolski & al., 2015; Takahashi et al., 2017), we include them in the Buxaceae *sensu lato* as distant, early diverging lineages. This inclusion necessitates the description of the two new subfamilies in Buxaceae (see below).

### Buxus

In *Buxus* s.l. three distinctive clades were recovered that correspond perfectly to their geographic distributions. The Eurasian species were sister to the remaining *Buxus* comprised of a New World and sub-Saharan African clade sister to one another. This topology is similar to that of von Balthazar & al. (2000) and Gutiérrez (2014), and all newly sequenced data fit well with these three geographical clades.

Classification of *Buxus* as a single genus vs. multiple genera is still contentious. The recognition of *Notobuxus* separate from *Buxus* is frequently but not totally accepted (*cf*. Friis, 1989; von Balthazar et al., 2000; Köhler, 2007, 2009). At the same time, the monophyletic *Tricera* has often been included in *Buxus* despite being molecularly closer to *Notobuxus* (von Balthazar et al., 2000). A reasonable approach based on the data presented here that does not require extensive nomenclature changes, is to recognize an inclusive *Buxus* with three well-supported monophyletic clades; we treat these Eurasian, African and American clades at a subgeneric level.

The Eurasian *Buxus* are highly diverse in southeastern Asia, with two or more species extending west to Europe (Köhler & Brückner, 1989; van Laere at al., 2011). The recently described *Buxus sirindhorniana* from Thailand (Soh & al., 2014; Soh & Parnell, 2018) and *B*. *rolfei* from the Philippines are members of the earliest lineage to diverge in the Eurasian *Buxus*. Our topologies (Fig. 4–5) do not recover the proposed infrageneric divisions of Mathou (1940) and Hatusima (1942). Of Hatusima’s (1942) informal groups, none are recovered as monophyletic though some of the species that form parts of his groups (namely, “Group I”) do form well-supported clades, e.g., *B. myrica*, *B. hainanensis*, *B. latistyla*, and *B. austroyunnanensis*. This last grouping also corresponds with the more recent review of Tianlu & Brückner (2008).

*Buxus* from Africa and Madagascar have variously been split into distinct genera separate from *Buxus*, or included within a broad *Buxus* (Van Tieghem, 1897; Friis, 1989; Schatz & Lowry II, 2002). Friis (1989) recognized three sections in African *Buxus*: sect. *Tricera* (Swartz ex Schreb.) Baill. including only *B. hildebrandtii*, sect. *Notobuxus* (Oliv.) Friis and sect. *Buxella* (Tiegh.) Hutch. These divisions are also supported by palynology (Köhler & Brückner, 1982; Köhler & Brückner, 1989; Köhler, 1994). Our molecular CP and OI analyses supported the recognition of sect. *Buxella*, excluded *B. hildebrandtii* from New World *Tricera* but did not fully resolve relationships between *B. hildebrandtii* and “sect. *Notobuxus*” species. We treat the African clade as subg. *Notobuxus*, a single monophyletic group; subclades recovered here might support further sectional divisions.

American *Buxus* clade forms a well-supported monophyletic taxon endemic to the Neotropics with a single migration and rapid diversification in the Caribbean-Cuban *Buxus* species (Köhler, 2014; Gutiérrez, 2014). Our CP trees were in line with Gutiérrez (2014). OI trees were different but overall support was much lower, and we cannot fully exclude the effect of putative ITS paralogues (Rosello et al., 2007; Gutiérrez, 2014). In all, we believe that barcoding markers are not powerful enough in cases of rapid diversification, and this group still awaits detailed research. However, but we were able to place there some newly sequenced species (Fig. 5), for example, *B. lancifolia* and *B. vaccinioides*.

### Pachysandroideae

The *Pachysandroideae* is united by is crotonoid pollen (Gray & Sohma, 1964; Köhler, 2007; see also Fig. 8), but the genera are otherwise morphologically distinctive. Our results of the *Pachysandroideae* present a topology (Figs. 2–6) where the long-questionably placed *Sarcococca conzattii* (Fig. 8) was consistently recovered separate from all other species of *Sarcococca.* In our analyses, it branches distantly from the rest of *Sarcococca* (OI, and especially “full ITS”: Figs. 5–6) or as a lineage sister to the *Sarcococca* (CP: Fig. 4). We used molecular data as an additional evidence and is herein recognized a novel monotypic genus, *Sealya* (see below).

**Figure 8.**
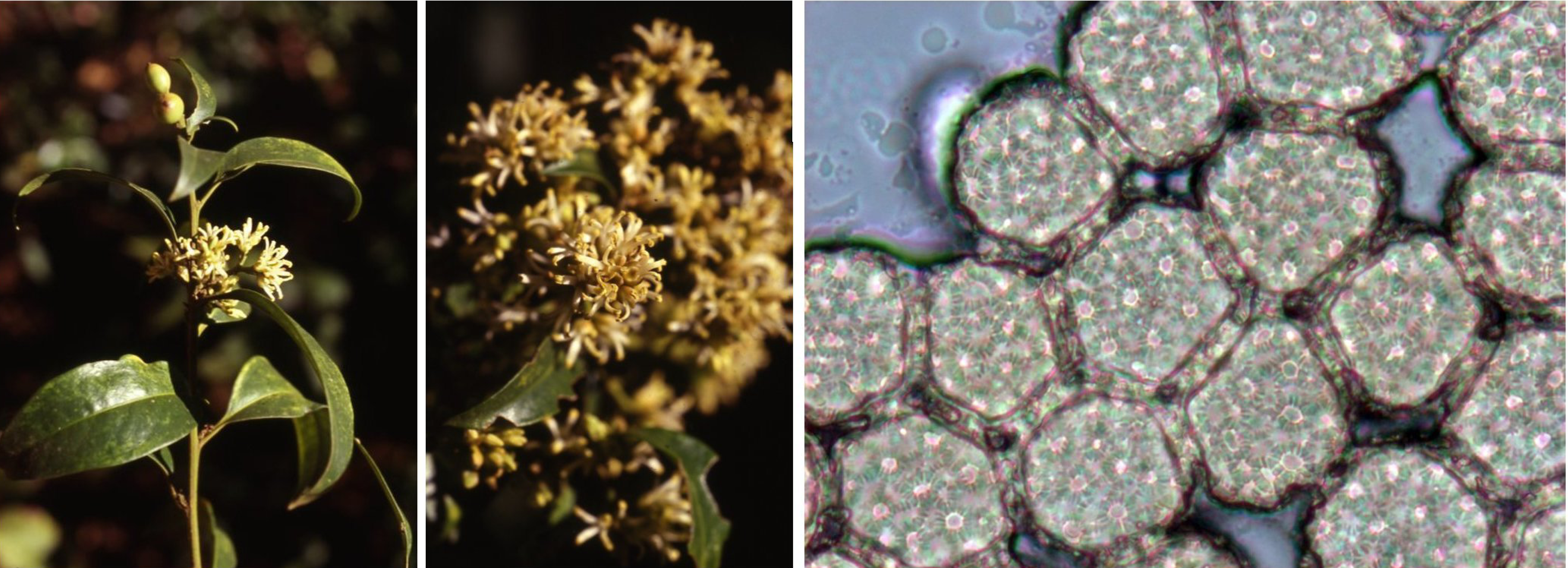
*Sealya conzattii*, left to right: branch, inflorescence, crotonoid pollen grains (photo credit to Jose Panero, Texas). This plant was also used for DNA extraction.

*Sealya* was nearly concurrently described in both *Buxus* and *Sarcococca* (Standley, 1936; Johnston, 1938), and its inclusion within *Sarcococca* was based on vegetative similarities (alternate, subcoriaceous leaves) and similar fruit types. However, it differs from *Sarcococca* in its inflorescence structure, which is similar to that of *Buxus* in having a terminal female flower; furthermore, its terminal female flowers possess the well-developed tepals and bracteoles (Sealy, 1986; Köhler, 2007). Its fruits are indehiscent like in *Sarcococca*, but have dry mesocarps and are white in contrast to the *Sarcococca* s.str. blackish-blue or reddish fruits (Sealy, 1986). *Sealya* shares the crotonoid pollen exine (Fig. 8) with all other members of *Pachysandroideae* (Köhler, 2007).

With *Sealya* excluded, *Sarcococca* is monophyletic in all our analyses. Even with our still limited sampling, it appears that the diversity in *Sarcococca* is higher than current taxonomic treatments recognize (Sealy, 1986; Min & Brückner, 2000). For example, our analyses recovered *S. euphlebia* and *S. balansae* separately and distantly placed, but Sealy (1986) placed *S. euphlebia* as a synonym of *S*. *balansae*. In contrast, Min and Brückner (2000) placed both as synonyms of *S. vagans* (which we did not sample). Inclusion of *S. vagans* might result in one of the preceding species being supported as a synonym, but not both. Additional systematic work is needed with a comprehensive sampling of widespread species to resolve taxonomy in *Sarcococca*.

Despite the low species number and relatively high level of systematic investigation (Robbins, 1968; Boufford & Xiang, 1992; von Balthazar et al. 2000; Jiao & Li, 2009), *Pachysandra* also presents a potential for the further studies. Our OI and “full ITS” analyses (Figs. 5–6) point on the possible separation of the *Pachysandra terminalis* from two other species of the genus. Morphologically, it differs from two other species in its terminal inflorescences, two-carpellate gynoecium, the smaller stigmatic region encompassing on the distal third of the style, the male flowers subtended by a coriaceous bract and two bracteoles, elongate multi-bracted pedicels of the pistillate flowers, white baccate fruits, and adaxially raised leaf veins (Robbins, 1968). Future molecular research should sample even more extensively and include *P. terminalis* from both areas of its distribution (Japan and China) as well as the broadly distributed and morphologically diverse *P. axillaris*.

Our coverage of *Styloceras* is comprehensive and represented here by seven samples and six species. *Styloceras laurifolium* holds the most basal position while the rest of the genus form two clades with the reliable support: *S. penninervium + S. kunthianum + S. connatum*, and *S. brokawii + S. columnare.* This topology does not correspond well with existing review (Torrez & Jorgensen, 2010) which places *S. kunthianum*, *S. columnare* and *S. laurifolium* together and separates *S. penninervium* from the rest of the genus. Given the scarcity of information and collections, and difficulties in DNA amplification in *Styloceras*, we believe that this is only a first step towards understanding the diversity of this rare and unusual Neotropical group.

## Taxonomic Treatment

### *Didymeles toamasinae* Floden et Shipunov, sp.nov.

Differs from *Didymeles perrieri* Leandri by smaller leaves (30–50 × 15–30 mm *vs*. 50–100 × 30–37 mm), intensively branching shoots, thinner terminal branches (1.5–2 *vs*. 2–4 mm diam.), and smaller (12–15 *vs*. 15–20 mm) mature fruits.

TYPE:—Toamasina. Fivondronana: Maroantsetra. Commume: Antisrasbesahatany. Fokontany: Anjiahely. Beanivona forest. Collected with Pascal, Ramaroson, Saside and David. 15°23’56“S 045°26’32”E. 746 m. Tree 8 m tall, DBH 20 cm, fruits nearly mature. 17 December 2002. *P. Antilahimena & al. 1529* (holotype: MO!)

Etymology:—Named after the region of collection, Toamasina, northeastern Madagascar.

Distribution:—MADAGASCAR, northeast, Atsinanana region. (MO *P. Antilahimena 2497a!*, MO *P. Antilahimena, Pascal & Ramaroson 1561*!)

Description:—Small trees 4–8 m, young stems 1–2 mm diam., internodes 2–4 cm. Leaves with petioles 1.2–1.5 cm, laminas 3–5 × 1.5–3 cm, elliptic, apex acuminate, base cuneate, with visible primary and secondary venation. Flowers not seen. Fruits 1.2–1.5 cm, ellipsoid, stylar remnants about 2 mm.

### *Buxus cyclophylla* Floden & Shipunov, nom.nov.

As the Cuban *Buxus rotundifolia* (Britton) Mathou is the latter homonym of *Buxus rotundifolia* K.Koch, we propose a new name for the former:

*Buxus cyclophylla* Floden & Shipunov, nom.nov. pro *Buxus rotundifolia* (Britton) Mathou, Rech. Fam. Buxac. 229 (1939). nom.illeg., non *Buxus rotundifolia* Hort. ex K.Koch, Dendrologie 2(2): 479 (1873).

≡ *Tricera rotundifolia* Britton, Bull. Torrey Bot. Club 42: 500 (1915).

Type: — NY *Shafer 114!* Between Camp La Barga and Camp San Benito, about 1,000 m alt., northern Oriente.

### ***Sealya*** Floden et Shipunov **gen. nov.**

Differs from *Sarcococca* by its inflorescence terminated by a female flower (*vs*. male), male and female flowers with well-developed tepals (*vs*. weakly differentiated as perianth organs), bracteoles prominent, and by its white fruit (*vs*. purple to red) with dry mesocarp (*vs*. fleshy mesocarp), while it also differs from *Buxus* by having alternate leaves, bicarpellate ovary (*vs*. tri), and fleshy, drupe-like fruit (*vs*. dry and dehiscent).

### Generic type

***Sealya conzattii*** (Standl). Floden et Shipunov **comb. nov**. TYPE:—Mexico, *C. Conzatti 2508* (holotype: F digital image!; isotypes: EAP87195 digital image!, G00359411 digital image!, US digital image!).

Basionym:—*Buxus conzattii* Standley, Publications of the Field Museum of Botany, 11: 163 (1936).

SYNONYMS**:—***Sarcococca conzattii* (Standley) I. M. Johnston, Journal of the Arnold Arboretum, 20: 240 (1939); *S. guatemalensis* I. M. Johnston, Journal of the Arnold Arboretum, 19: 121 (1938). — Type: Guatemala, *A.F. Skutch 288* (holotype: A00048976 digital image!; isotype: K000573596 digital image!, MICH digital image!, TX/LL!, US digital image!)

Etymology:—Named in honor of J. Robert Sealy who stated in his 1986 revision of *Sarcococca* that the generic placement of *S*. *conzattii* in *Sarcococca* was incorrect and its placement in *Buxus* would be “equally anomalous.”

Distribution:—MEXICO. Chiapas, Oaxaca. GUATEMALA, occurring at mid-to higher elevations.

### ***Buxus*** subgenus ***Notobuxus*** (Oliv.) Floden et Shipunov, **subg.nov.**

Basionym: —*Notobuxus* Oliv. in Hook, Ic. Pl. 14:78, t 1400. 1882. Type: —*B. natalensis* (Oliv.) Hutch., Africa, Natal, Inanda, *Wood 1357* (lectotype (designated here), K, isolectotypes, BOL136821, BOL136822, NH0001717-1, NH0001717-2).

Subgenus *Notobuxus* has been repeatedly shown in molecular studies to be a monophyletic group of African and Malagasy species (Gutiérrez, 2014; von Balthazar & al., 2000). It is characterized by frequently sessile anthers, and by the frequent presence of multiple (6–10) stamens and flat disk-shaped pistillode (Friis, 1989; von Balthazar & Endress, 2002a; Köhler, 2007). *Notobuxus* Oliv. was based on two separate collections that represent syntypes because no type was designated (Article 9.3: Turland & al., 2018). From the original material (Article 9.4) cited by Oliver (1882) there was a collection with unopened flowers (*T. Cooper* 3465, K) and a collection by *Wood 1357* with opened flowers. The accompanying illustration (Oliver, 1882: Plate 1400) was drawn from the latter, thus we select *Wood 1357* to serve as the lectotype.

### ***Buxus*** subgenus ***Tricera*** (Swartz ex Schreb.) Floden et Shipunov, **subg.nov.**

Basionym: —*Tricera* Swartz ex Schreb., Gen. 630. 1791. Type:—*B. laevigata* (Sw.) Spreng. Jamaica, *O.P. Swartz* (holotype S, isotypes G, LD).

Subgenus *Tricera* is wholly New World and differs from the subgenus *Buxus* and subgenus *Notobuxus* by the absence of cortical vascular bundles in the angle of the branchlets (Köhler, 2007). The results of the molecular phylogeny support subg. *Tricera* as a robust monophyletic group endemic to the New World.

### *Didymeloideae* Floden & Shipunov, subfam. nov.

*Didymeloideae* Floden & Shipunov, subfam. nov. – Type: *Didymeles* Thouars

Distribution. – Madagascar.

Genera (1). – *Didymeles* Thouars

### *Haptanthoideae* Floden & Shipunov, subfam. nov.

*Haptanthoideae* Floden & Shipunov, subfam. nov. – Type: *Haptanthus* A. Goldberg & C. Nelson

Distribution. – Central America, Honduras, Atlantida province.

Genera (1). – *Haptanthus* A. Goldberg & C. Nelson

### *Kichijiso* Floden et Shipunov gen. nov.

Differs from *Pachysandra* by its terminal inflorescences, gynoecium with two-carpels, a stigmatic region encompassing only the distal third of the style, male flowers subtended by a coriaceous bract and two bracteoles, the elongate multi-bracted pedicels of the pistillate flowers, its white baccate fruits, and its adaxially raised leaf veins.

Etymology: The generic name, *Kichijiso*, is one of the Japanese common names of this plant (Batchelor & Miyabe, 1893: “Kichijisô, the fruit of which are eaten raw”). This selection of the generic name follows other Latinized generic names such as *Aucuba*, *Kirengeshoma*, *Nandina*, and *Sasa* (Stearn, 2004).

Distribution: Japan and eastern China (Min Tianlu & Brückner, 2008)

### Generic type

***Kichijiso terminalis*** (Sieb. et Zucc.) Floden et Shipunov **comb. nov**. TYPE:—(lectotype selected by Robbins, 1960) Japonia, 1842, *Siebold, P.F. von, s.n.* (M-0120840).

Basionym:—*Pachysandra terminalis* Siebold & Zuccarini, Abh. Math.-Phys. Cl. Königl. Bayer. Akad. Wiss. 4(2): 142. 1845.

## Conclusion

With the broad taxonomic sampling, we provide a comprehensive approach to improve the classification scheme of the Buxaceae. We hope that our results might serve as a framework for future studies in the group, which will eventually reach the highest phylogenetic accuracy. The use of short barcoding molecular markers is likely the best choice for our mostly herbarium-oriented approach. However, at the same time, we argue that more in-depth research that incorporates data on genomics, morphology, anatomy, chemistry, distribution, and fossil history is required to provide overall integrative and synthetic revisions of the genera of the Buxaceae.

### Key to the Buxaceae *sensu lato*

1 Plants dioecious; flowers apetalous; male with one stamen pair; female flowers paired, carpels single and uni-ovulate, seeds exalbuminous … ***Didymeles*** (subfam. *Didymeloideae*)

– Plants monoecious; flowers with weakly differentiated perianth parts; male flowers with decussate tepals, 2, 4, or 6–10 stamens; female with spiraled tepals, carpels bi-to pluri-ovulate, seeds albuminous … 2

2 Flowers apparently naked; male flowers with two stamens fused into one staminate organ; female flowers 3-carpellate, carpels pluri-ovulate (8–15), placentation parietal … ***Haptanthus*** (subfam. *Hapanthoideae*)

– Flowers with perianth; male flowers with free stamens, female flowers 2–3-carpellate, carpels bi-ovulate, placentation axile … 3

3 Tepals absent in male flowers, stamens numerous; rudiment of ovary wanting … ***Styloceras*** (subfam. *Pachysandroideae*)

– Tepals present; stamens usually 4, rarely 6–10 … 4

4 Leaves decussate; female flowers terminal in racemes or clusters; fruit a 3-valved capsule … 5 (*Buxus*, subfam. *Buxoideae*)

– Leaves alternate; female flowers at base of racemes or spikes; fruit more or less drupaceous … 7 (the rest of subfam. *Pachysandroideae*)

5 Cortical vascular bundles wanting (American) … ***Buxus*** subg. ***Tricera***

– Cortical vascular bundles in each angle of the branches … 6

6 Cortical vascular bundles with fibre strands; male flowers with 4 stamens, anthers long exserted; pistillode present … ***Buxus*** subg. ***Buxus***

– Cortical vascular bundles without fibre strands; male flowers with (4) 6–10 stamens; anthers usually sessile; pistillode as a flat disk, or absent … ***Buxus*** subg. ***Notobuxus***

7 Woody shrubs or small trees; leaves entire; fruit more or less drupaceous … 8

– Perennial herbs with procumbent stems; leaves serrate to dentate or deeply toothed; flowers borne at the base of the stem or terminally; fruit an indehiscent capsule or subdrupaceous … 9

8 Female flowers lateral on inflorescences; gynoecium 2-or 3-carpellate; fruit drupaceous, reddish, purple, or blackish … ***Sarcococca***

– Female flowers terminal on inflorescences; gynoecium 2-carpellate; fruit with dry mesocarp, white … ***Sealya***

9 Inflorescences at base of stem or proximal axils; gynoecium 3-carpellate; fruits reddish-brown, indehiscent capsule … ***Pachysandra*** (*P. axillaris* and *P. procumbens*)

– Inflorescences terminal; gynoecium 2-carpellate; fruits white, subdrupaceous … ***Pachysandra terminalis***

**(Kichijiso terminalis)**

## Acknowledgements

Our research was supported by North Dakota INBRE and by the Department of Biology of Minot State University. Some of the sequences are results of collaboration with the Barcoding of Life project. We are grateful to the curators of all herbaria provided us with samples, namely US, NY, HUH, MO, CAS, UC, JEPS, K, F, BRIT, B, BO, IBSC, MICH, MO, NY, PE, TI, SPF, SP, USM, PRE, NBG, SAM. AF thanks Dr. Schilling for his encouragement and allowance of taking on extracurricular projects into my many interests while working towards my Ph.D., Dr. Jose Panero for sending the duplicate specimens that instigated part of this project, Joe May and Veronica Brown at the UT Genomics Core, Kelly and Sue at Far Reaches Farm for an additional *Sarcococca wallichii*, Dr. P.D. González Gutiérrez for discussion of his thesis work of Neotropical *Buxus*, and Dr. T. Mitchell and B. Wynn-Jones for support in Vietnam. AS thanks Dr. Hidetoshi Nagamasu (University of Kyoto) and Ms. Chikako Hasekura (Tokyo University of Agriculture) for the help with Japanese plants, and Ekaterina Shipunova, Maxim Nuraliev (Moscow State University), Polina Volkova (Papanin Institute for Biology of Inland Waters, Russia), Maria Kuzmina (University of Guelph), and Subdirección Científica, Jardín Botánico de Bogotá José Celestino Mutis (Bogota, Colombia) for help with obtaining and sequencing samples.

## Author Contributions

AS provided the design of the research; AF and AS performed data analysis, interpretation and writing of the manuscript; all co-authors participate in data collection.

## Support materials

**Support Table 1.**
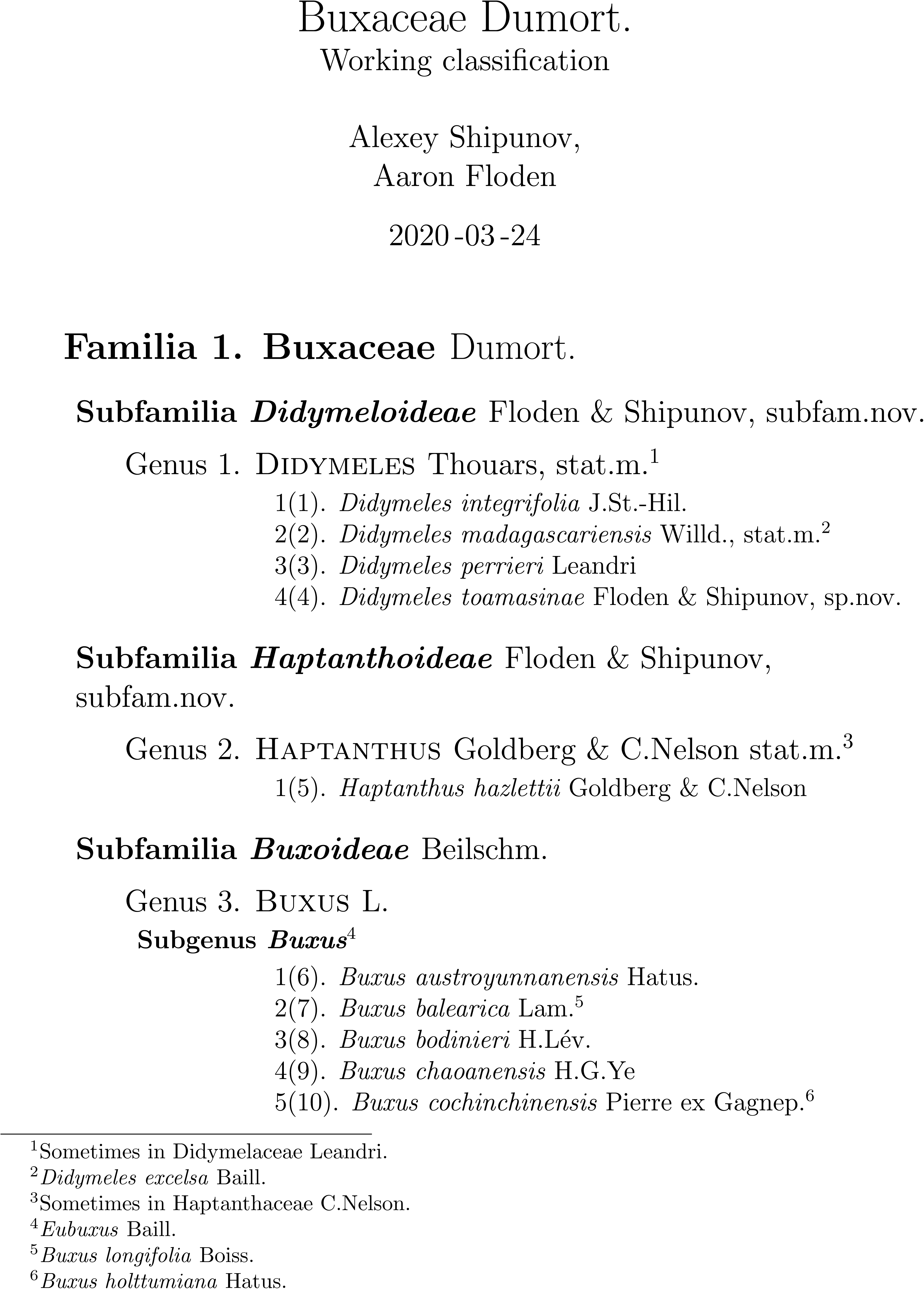
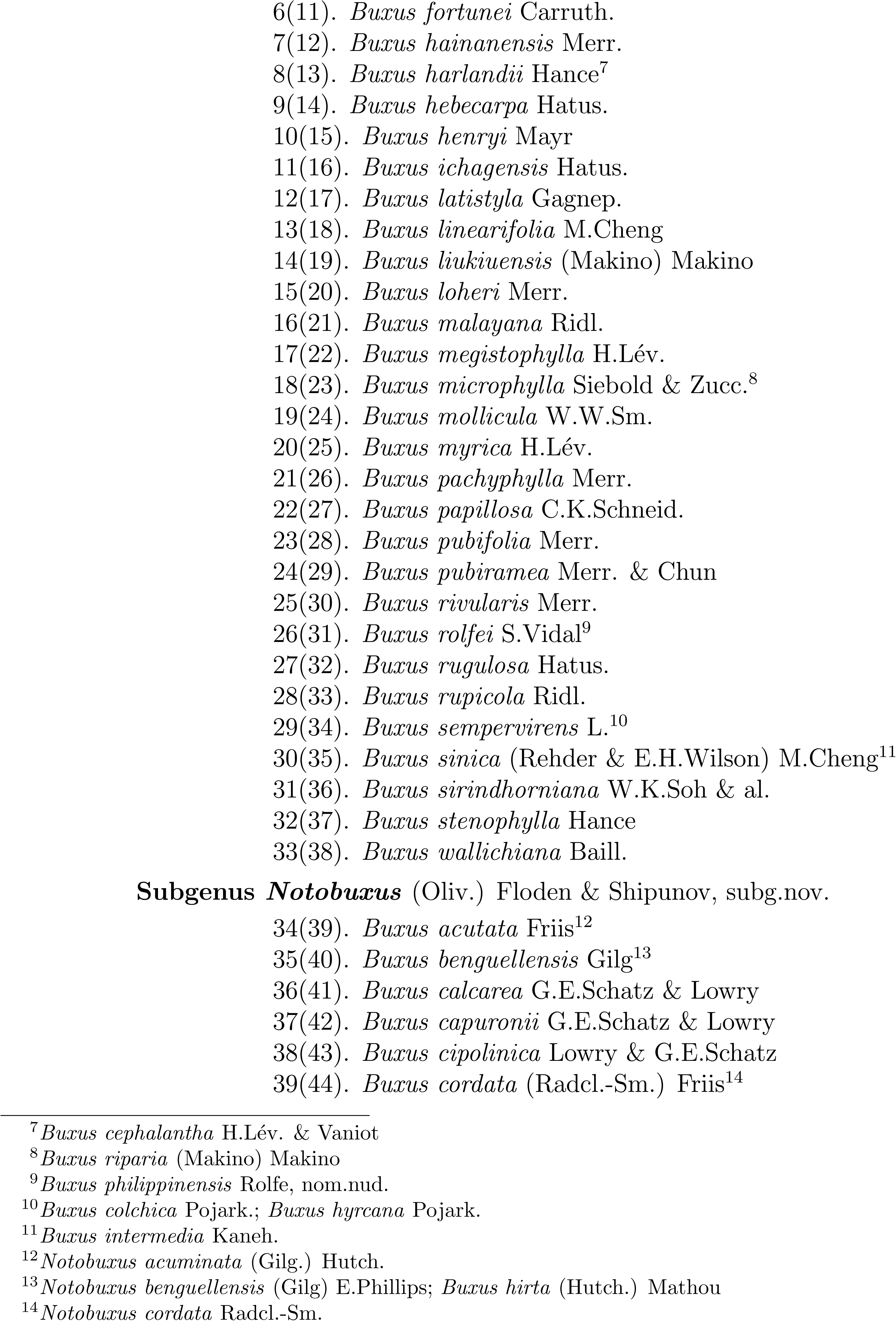
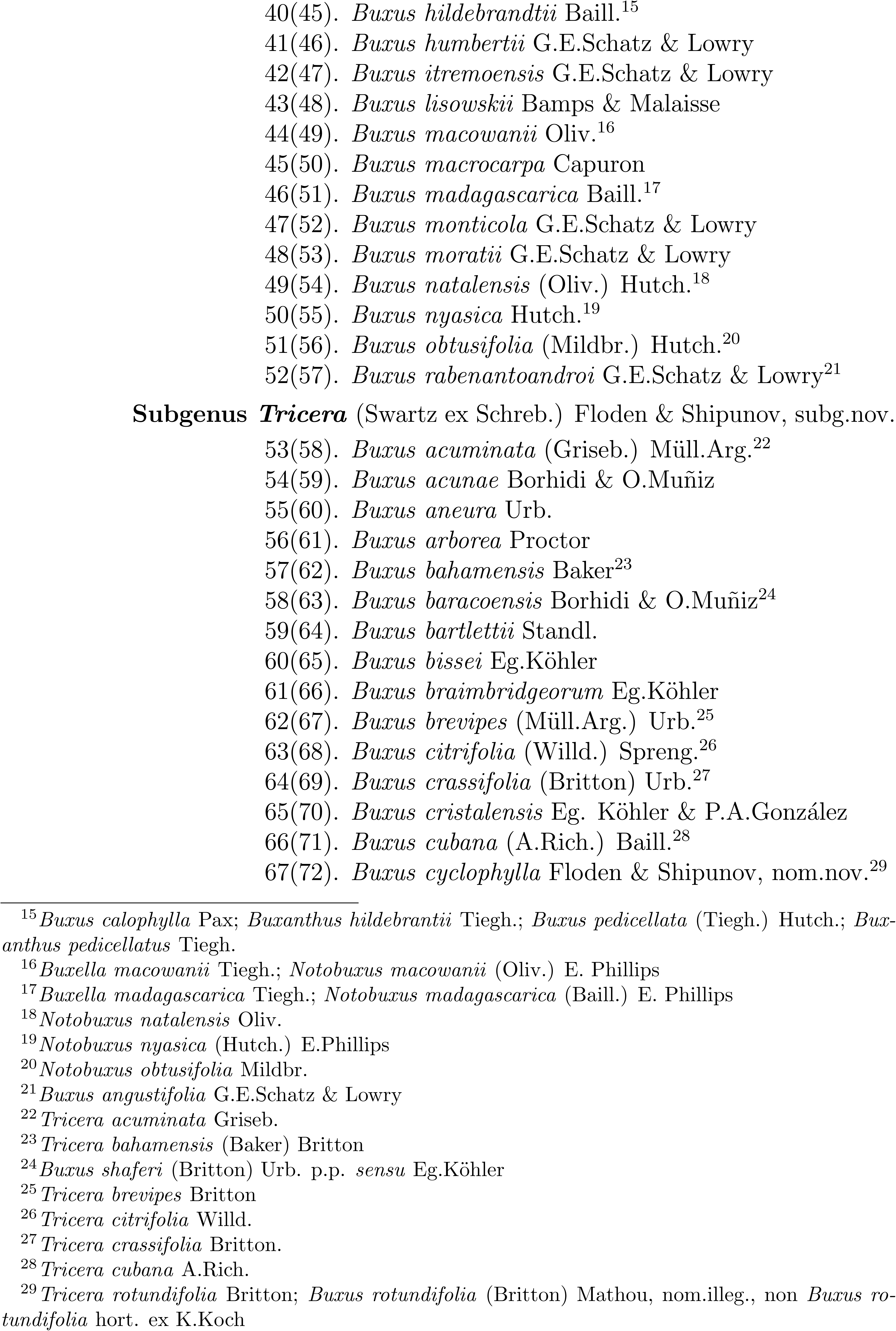
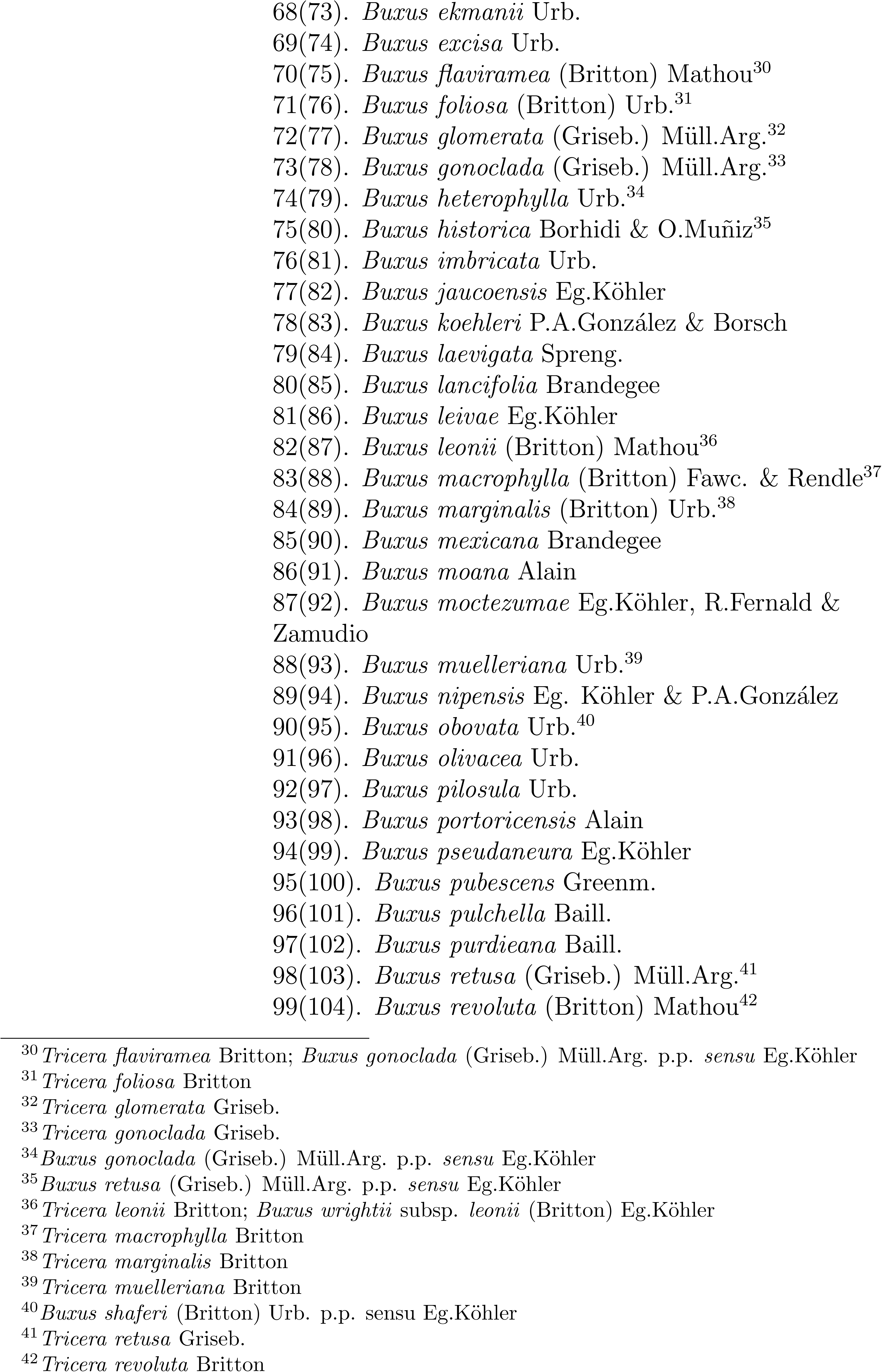
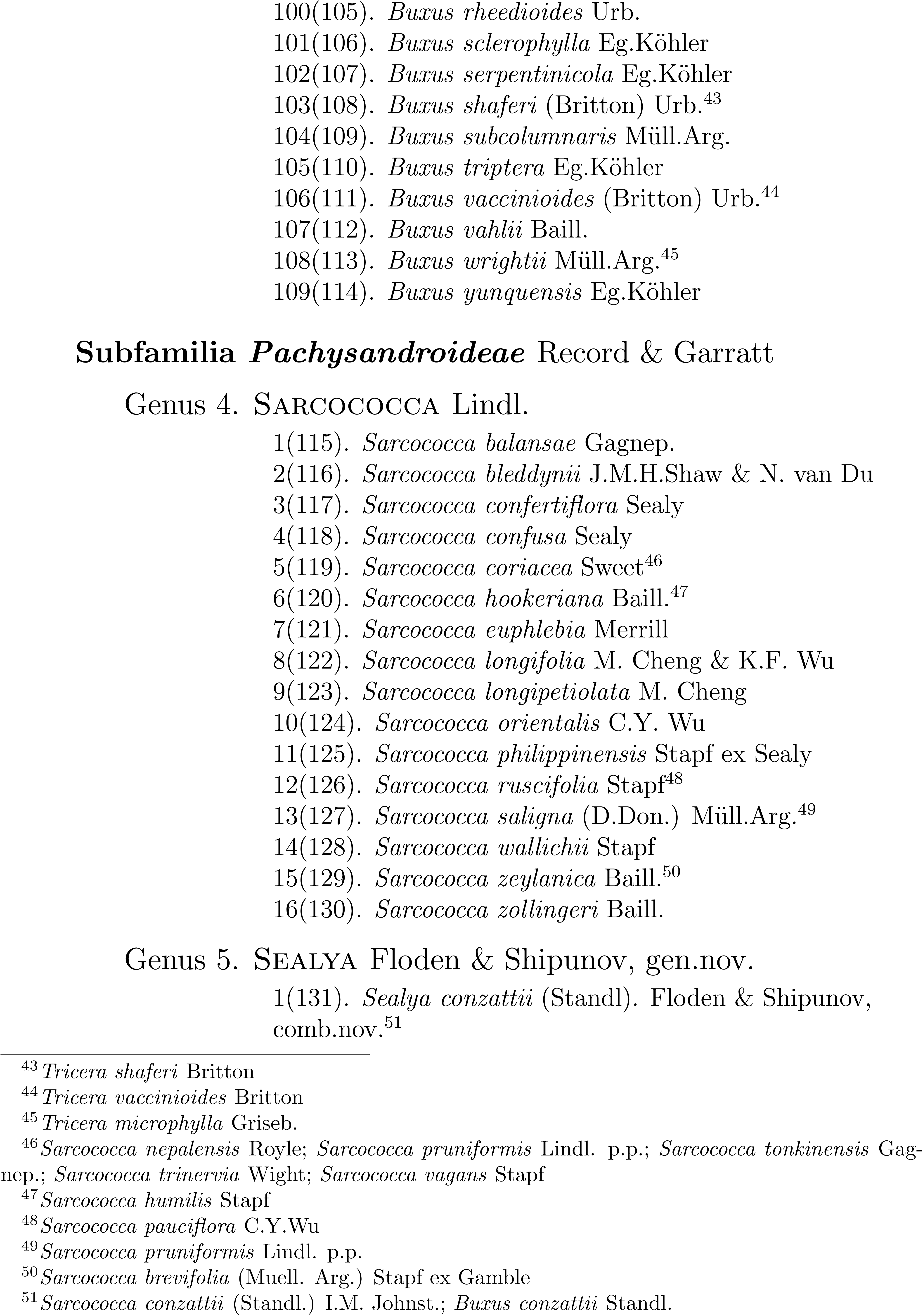
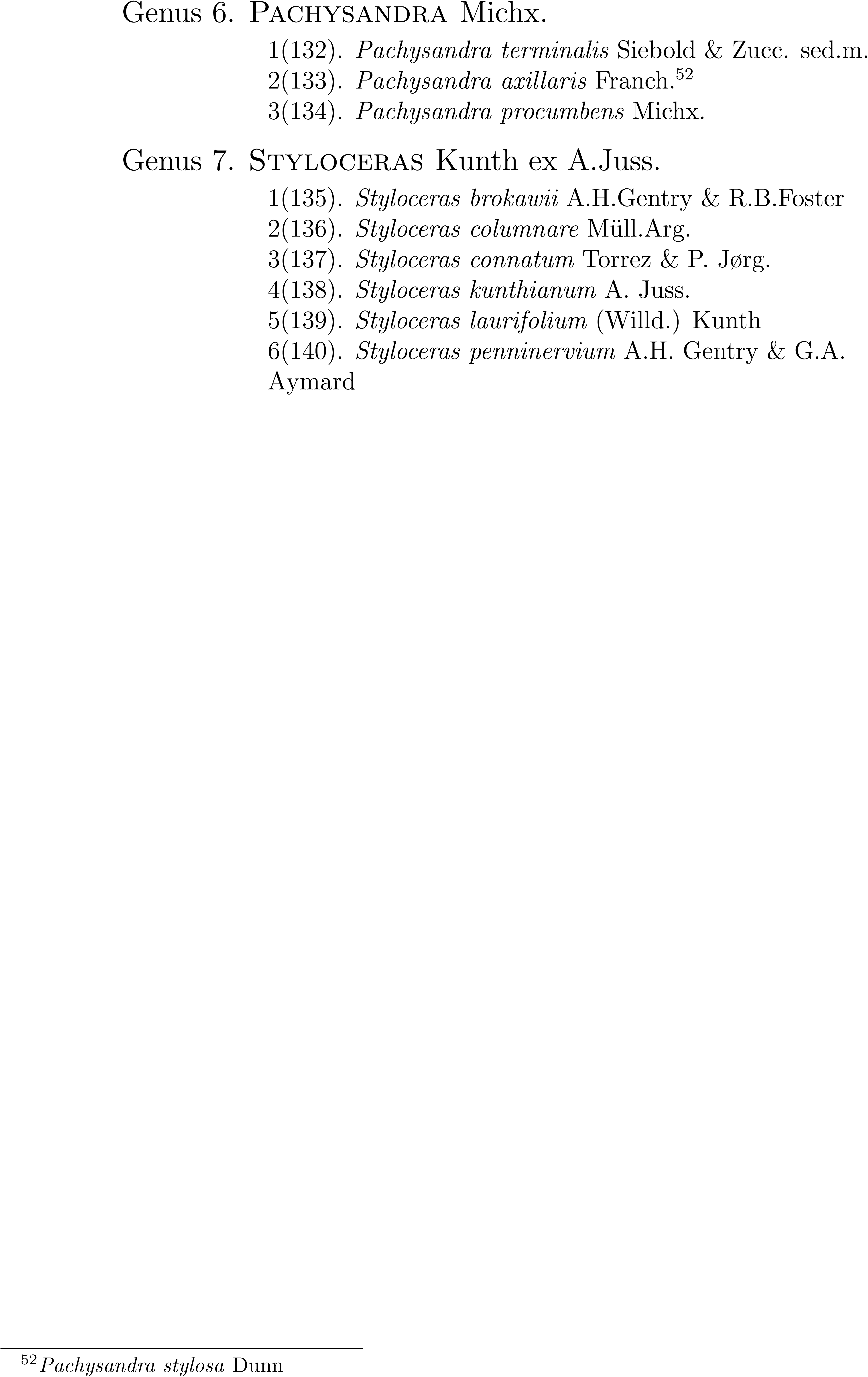
Working classification of Buxaceae.

**Support Table 2.**
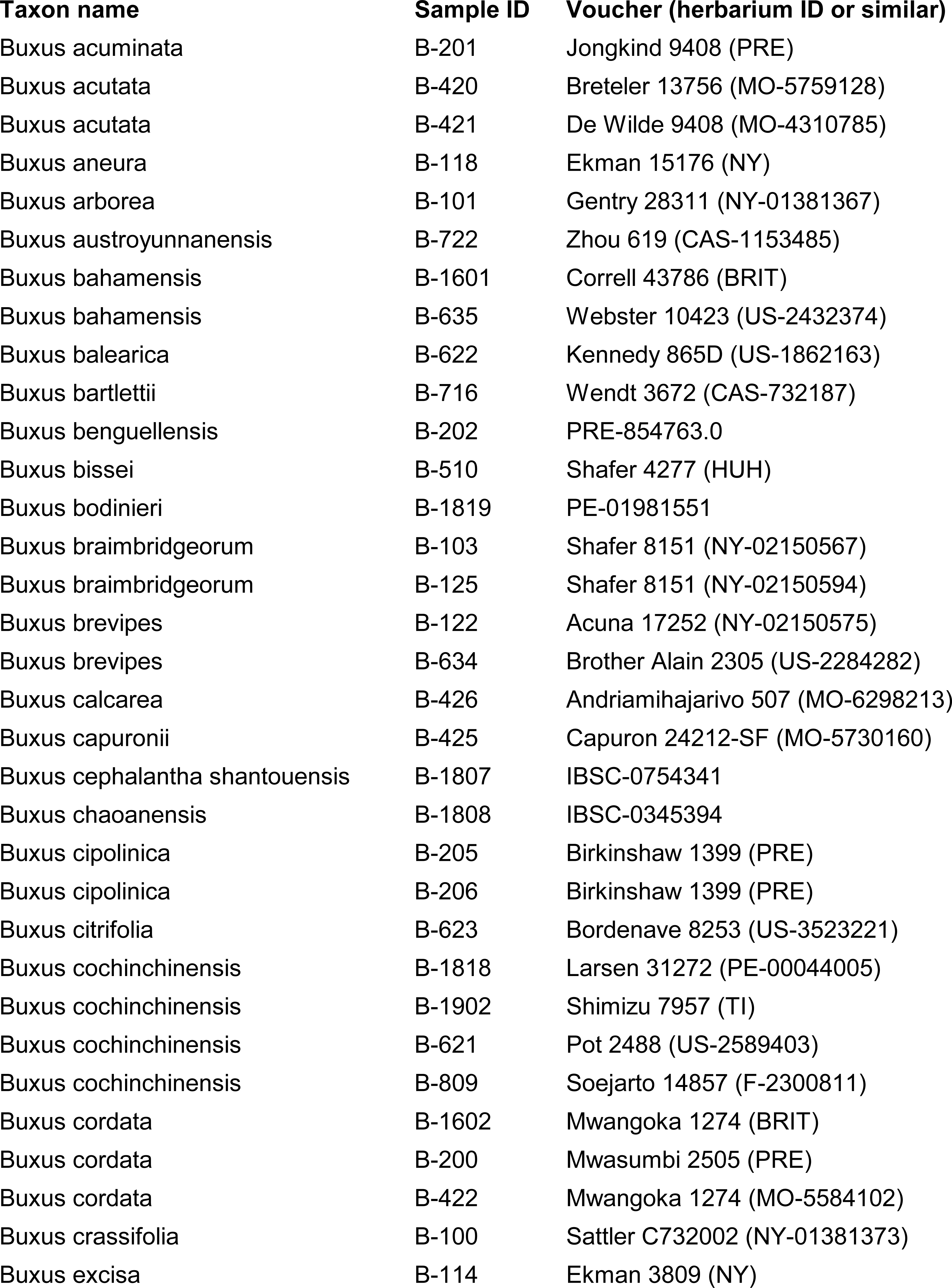
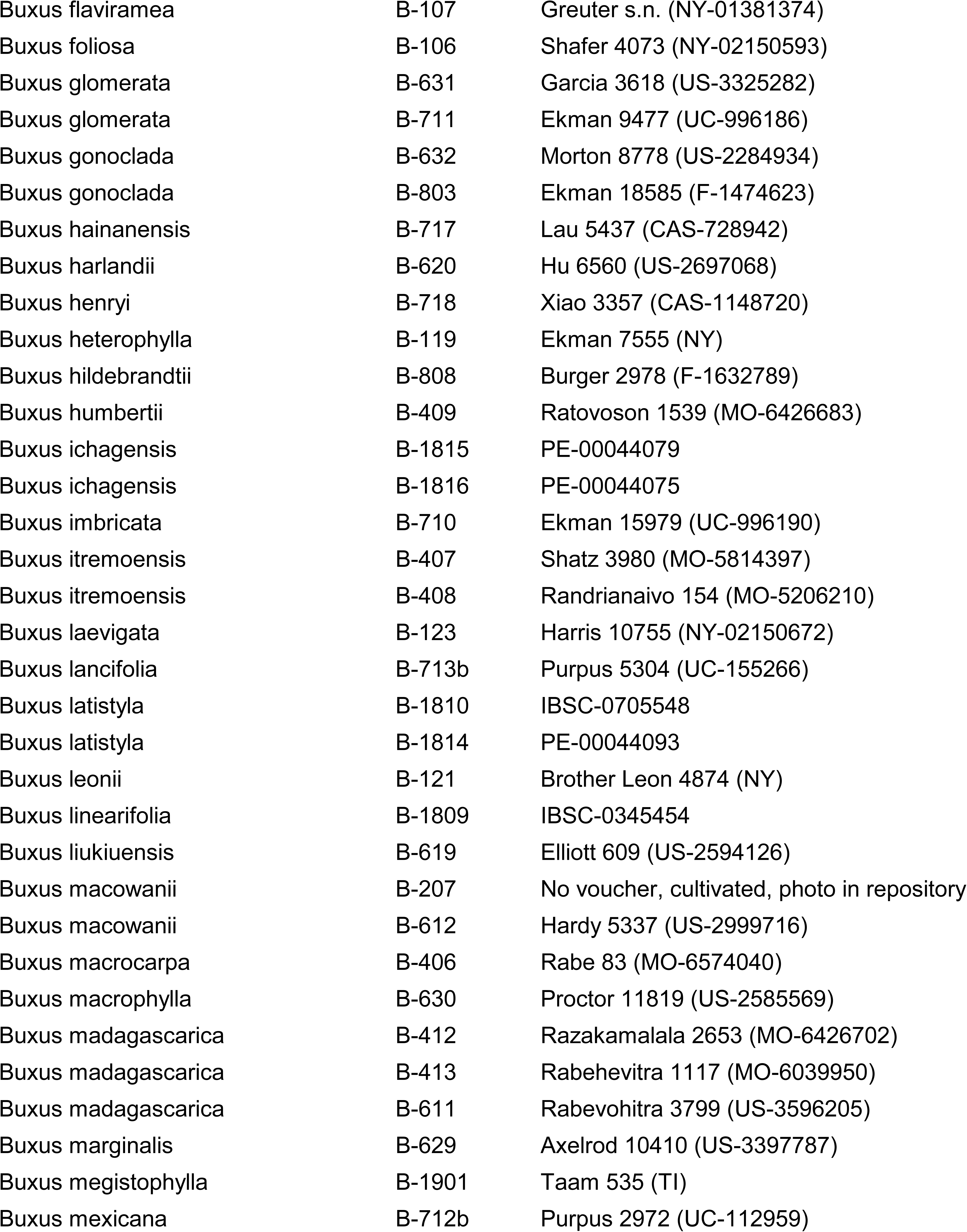
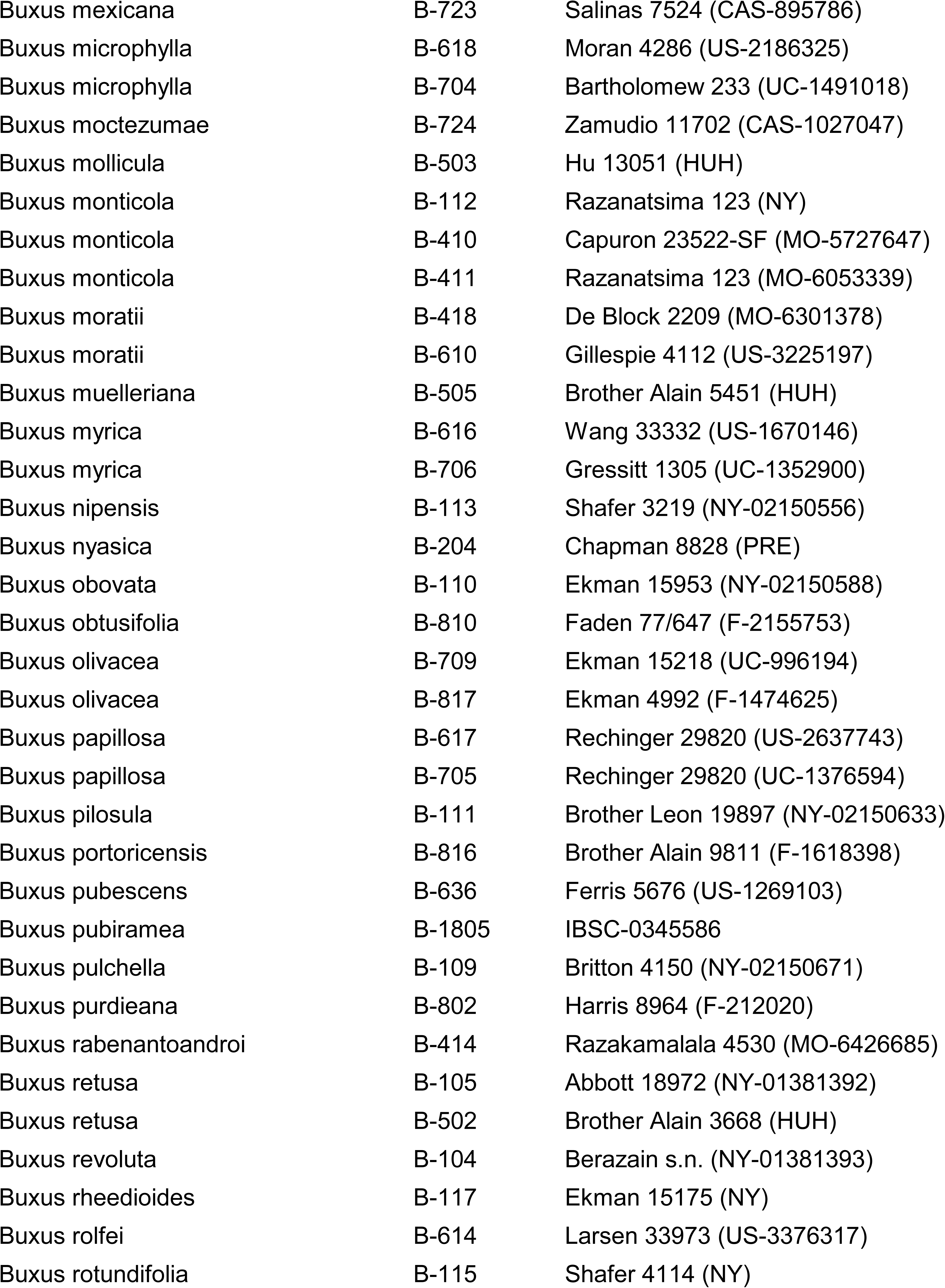
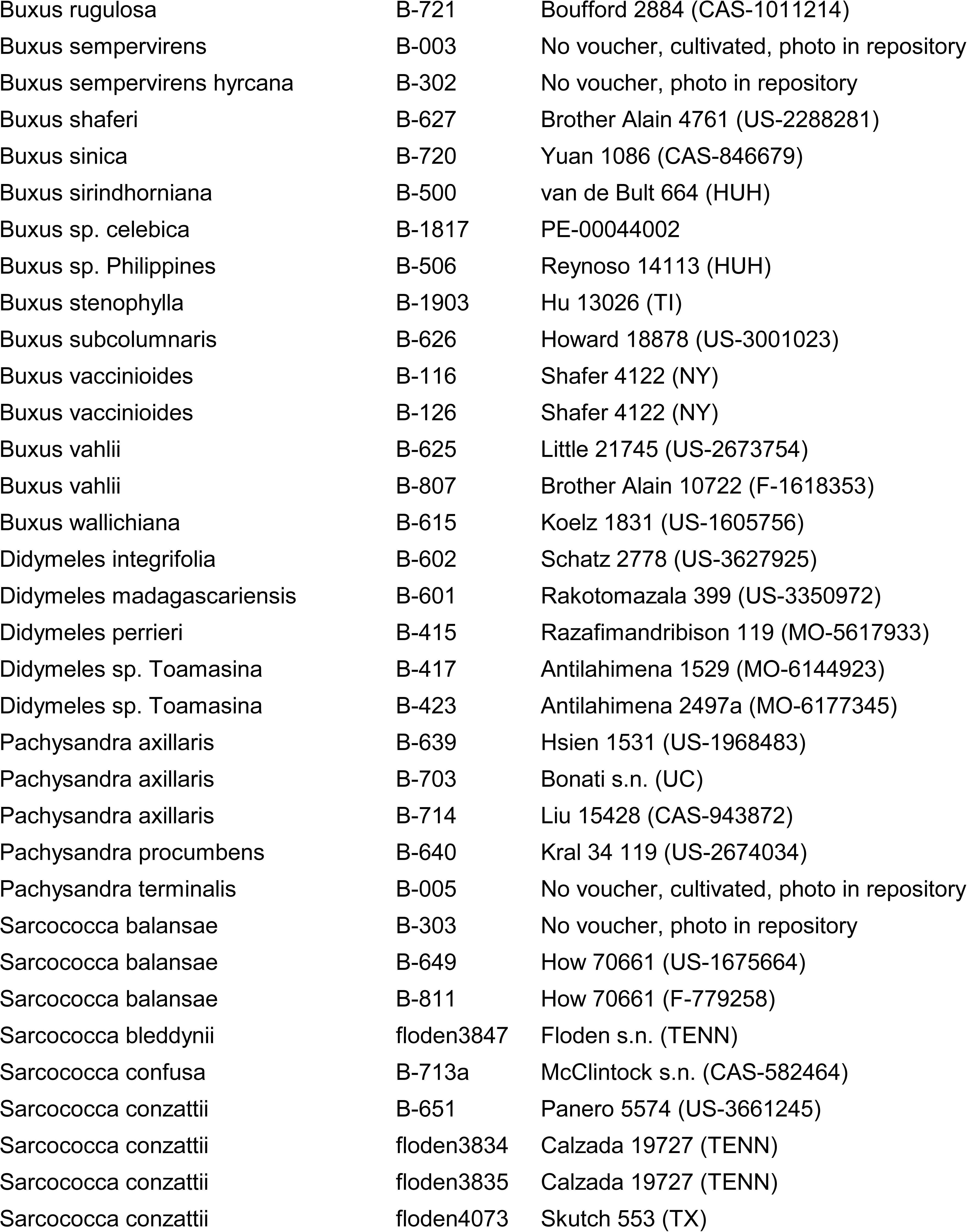
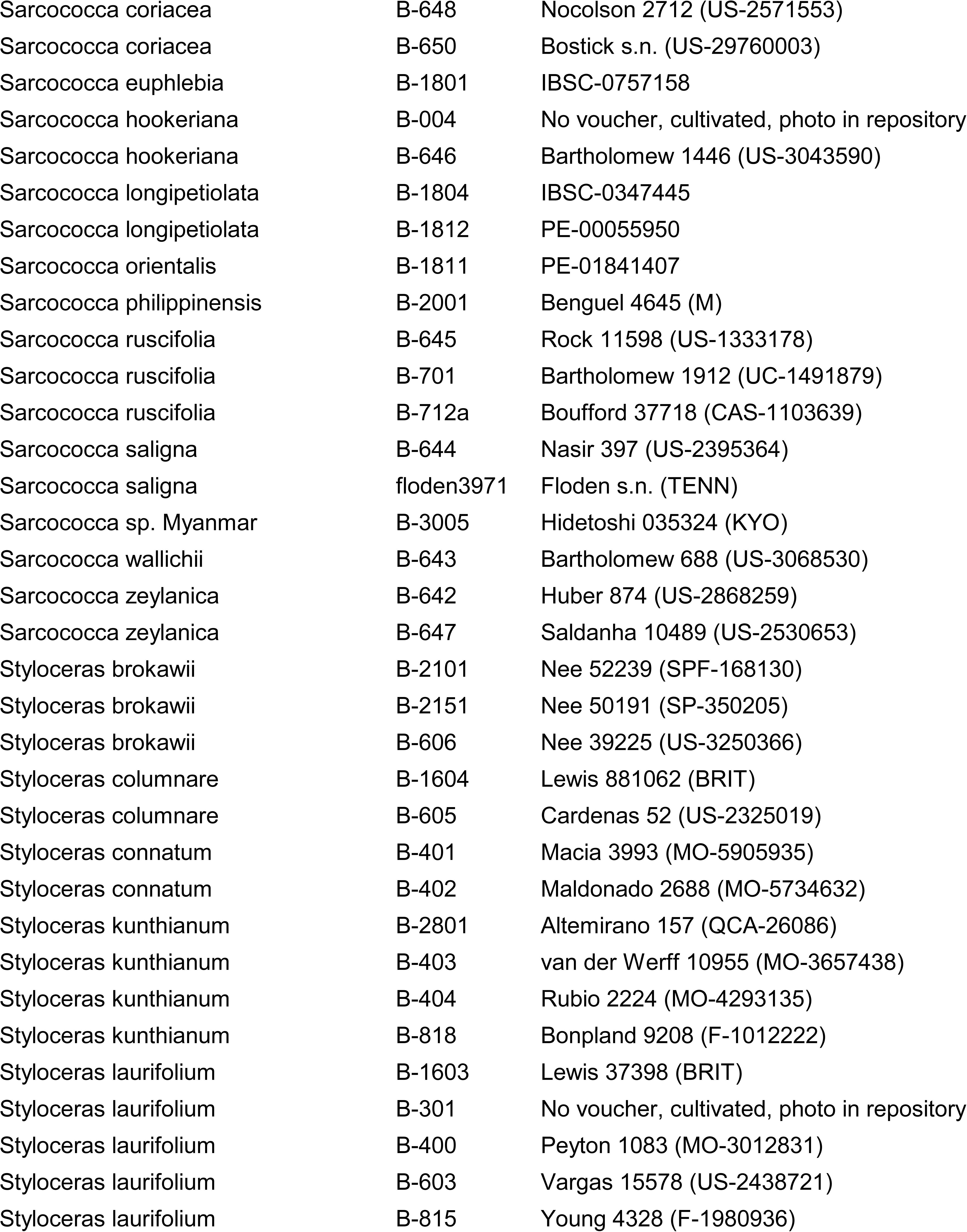
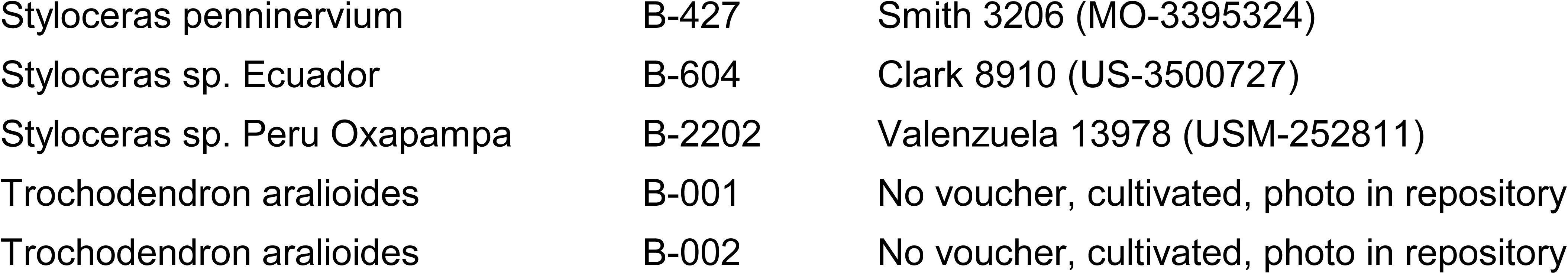
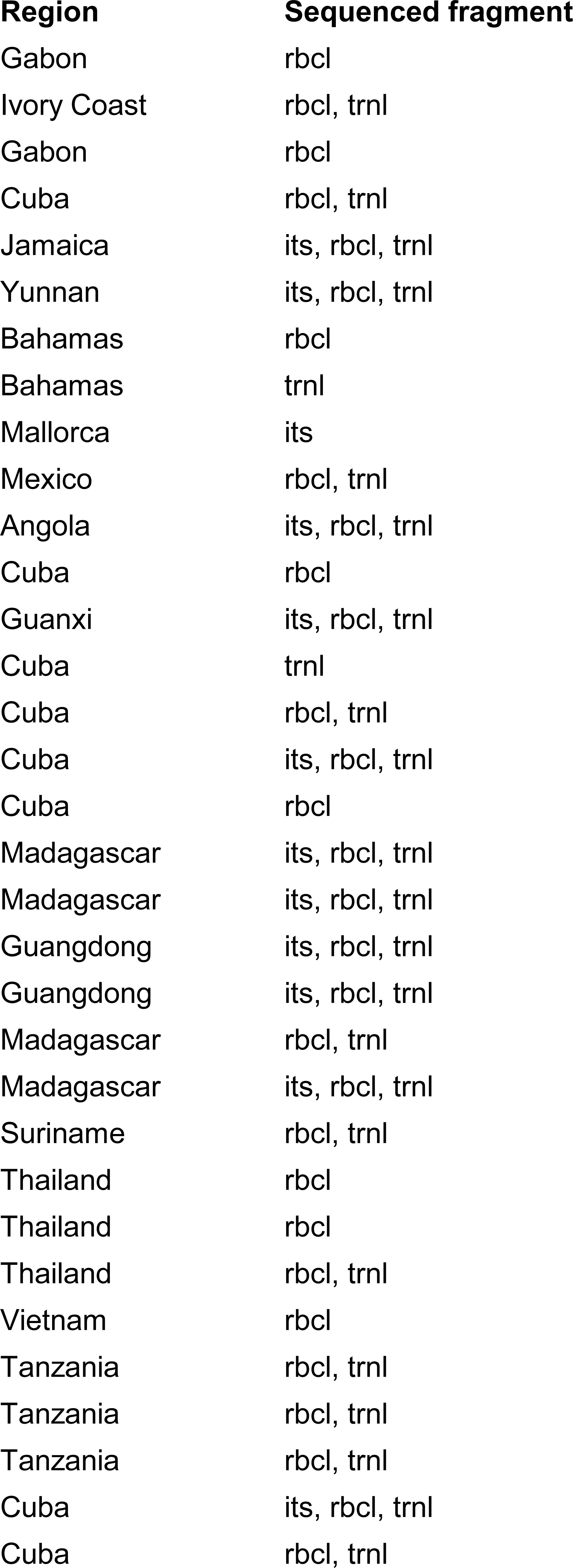
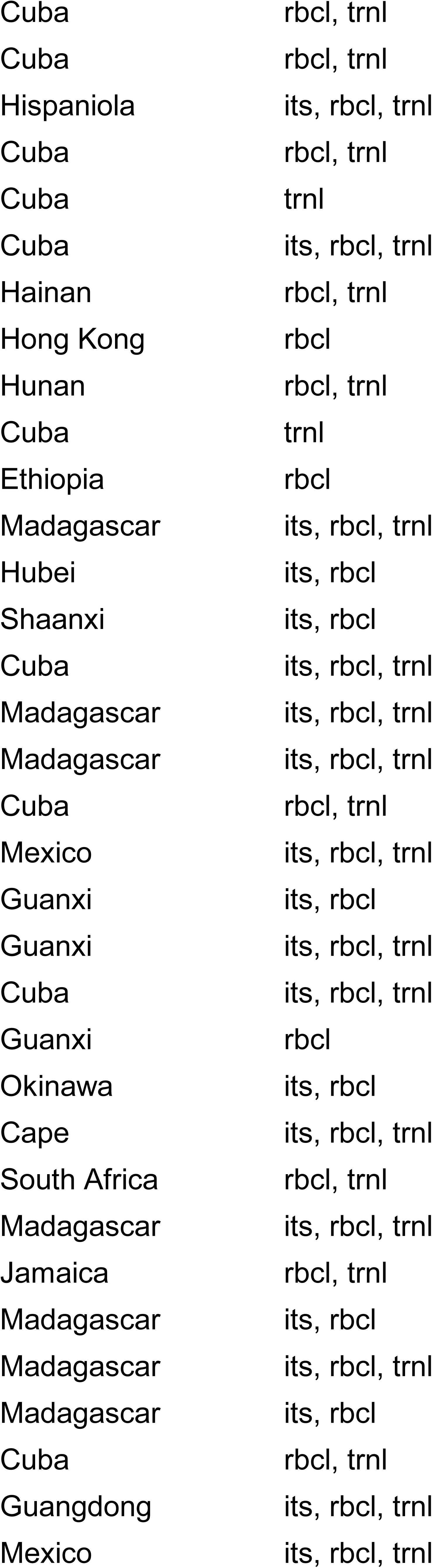
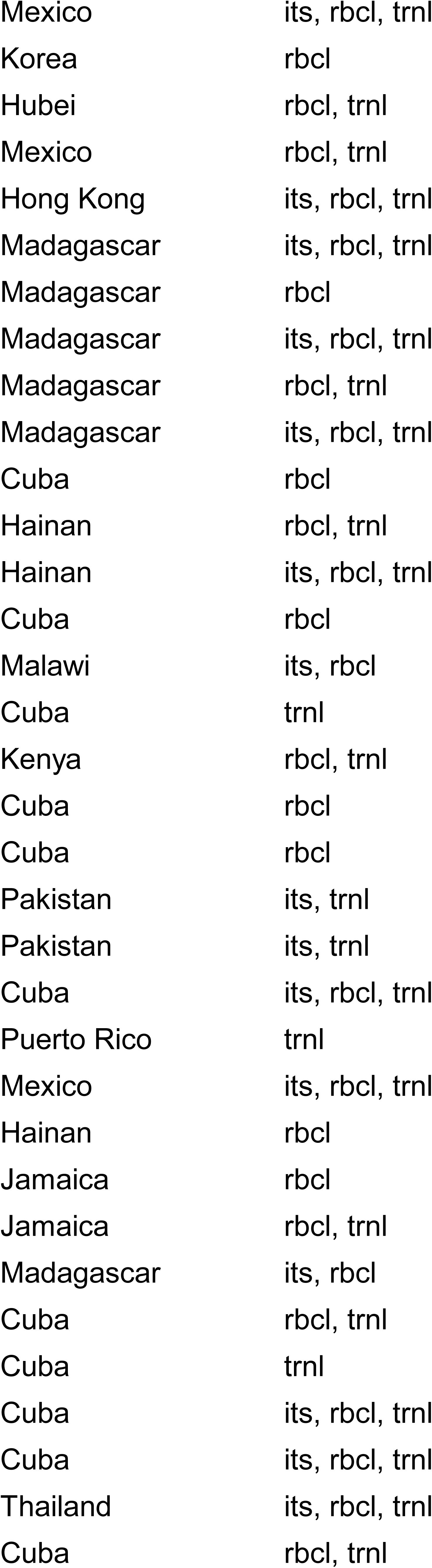
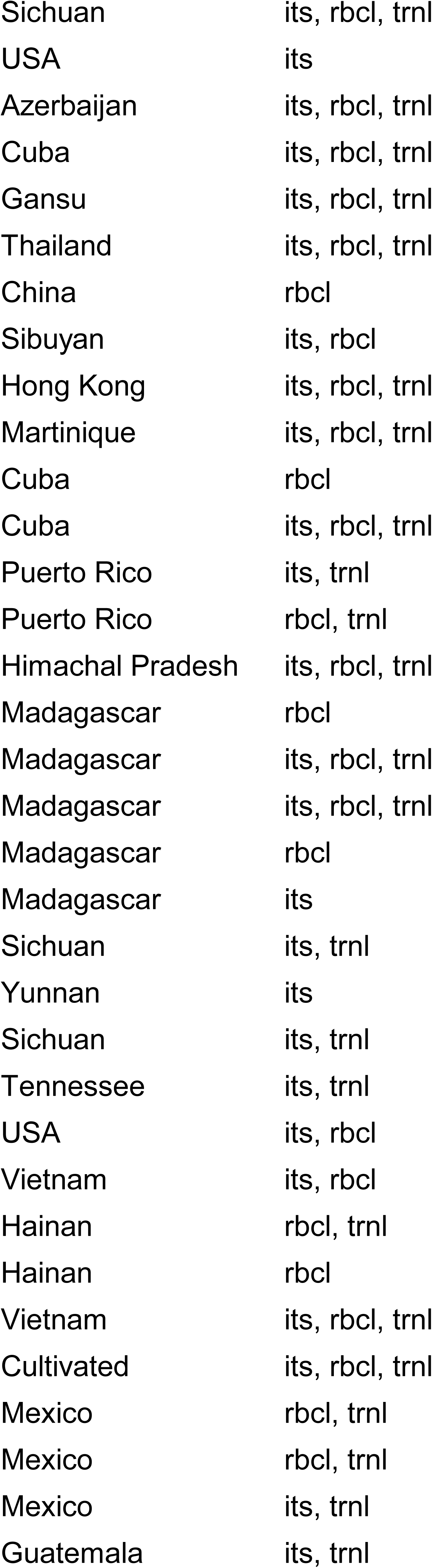
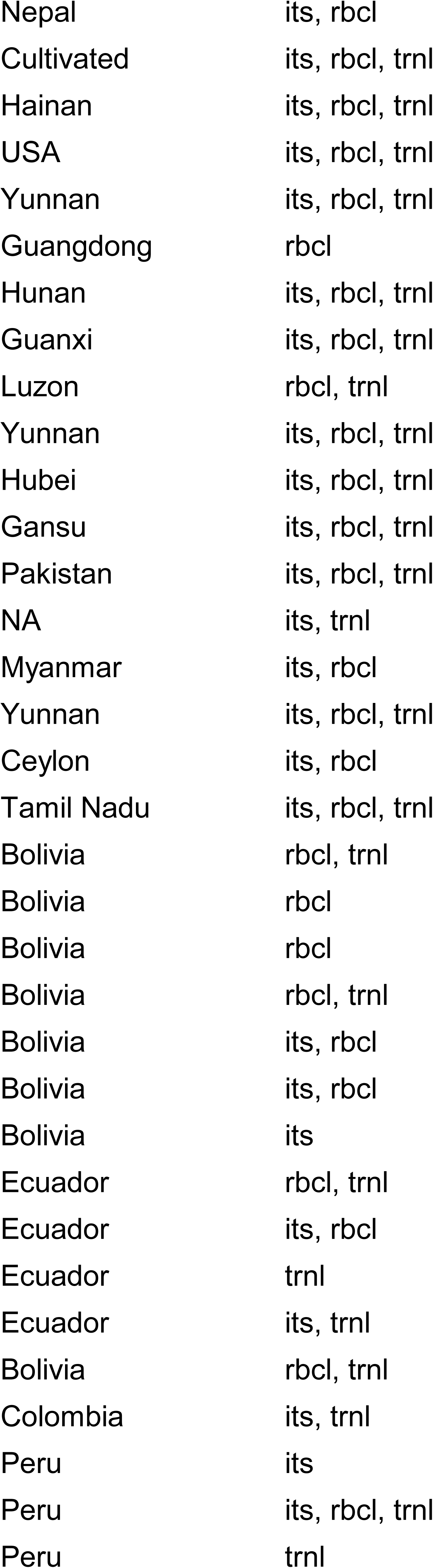
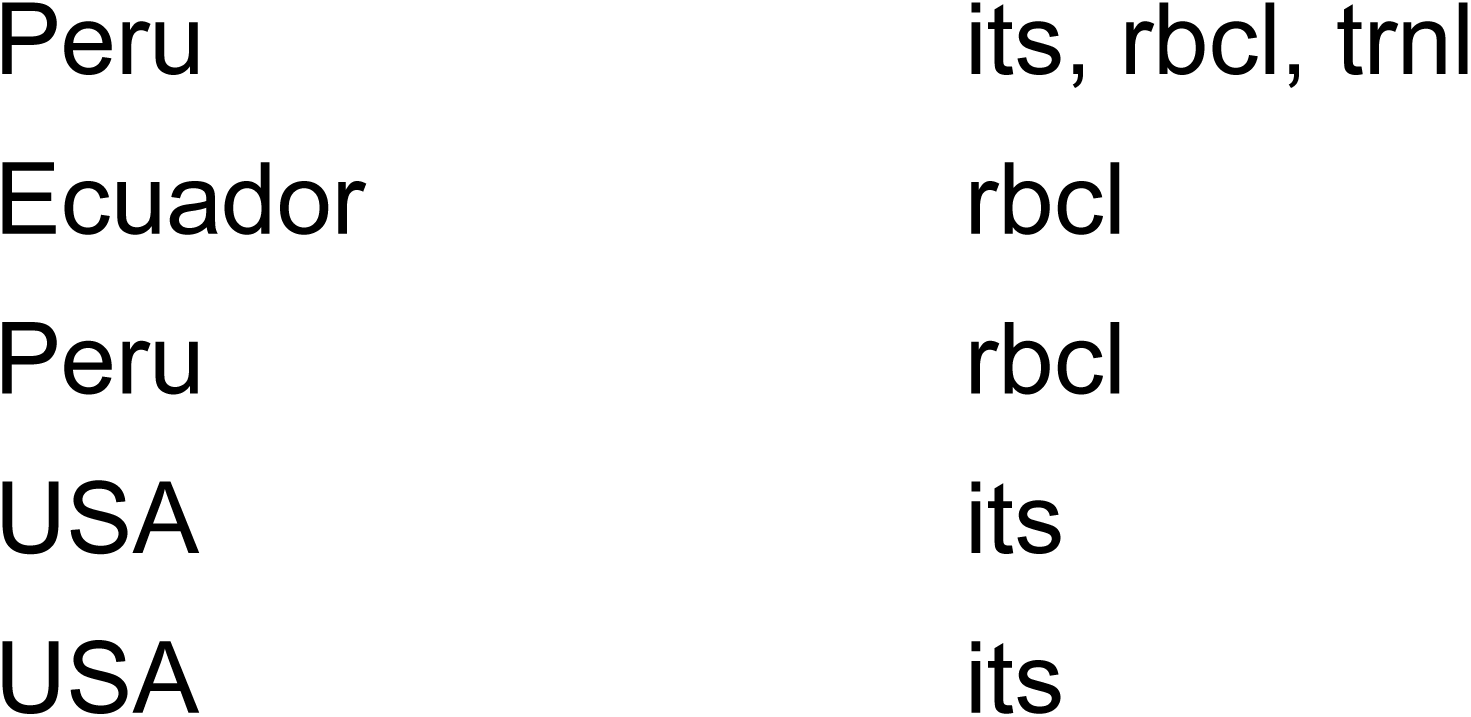
Vouchers of Buxaceae samples.

| **Taxon name** | **Sample ID** | **Voucher (herbarium ID or similar)** |
| --- | --- | --- |
| *Buxus acuminata* | B-201 | Jongkind 9408 (PRE) |
| *Buxus acutata* | B-420 | Breteler 13756 (MO-5759128) |
| *Buxus acutata* | B-421 | De Wilde 9408 (MO-4310785) |
| *Buxus aneura* | B-118 | Ekman 15176 (NY) |
| *Buxus arborea* | B-101 | Gentry 28311 (NY-01381367) |
| *Buxus austroyunnanensis* | B-722 | Zhou 619 (CAS-1153485) |
| *Buxus bahamensis* | B-1601 | Correll 43786 (BRIT) |
| *Buxus bahamensis* | B-635 | Webster 10423 (US-2432374) |
| *Buxus balearica* | B-622 | Kennedy 865D (US-1862163) |
| *Buxus bartlettii* | B-716 | Wendt 3672 (CAS-732187) |
| *Buxus benguelensis* | B-202 | PRE-854763.0 |
| *Buxus bissei* | B-510 | Shafer 4277 (HUH) |
| *Buxus bodinieri* | B-1819 | PE-01981551 |
| *Buxus braimbridgeorum* | B-103 | Shafer 8151 (NY-02150567) |
| *Buxus braimbridgeorum* | B-125 | Shafer 8151 (NY-02150594) |
| *Buxus brevipes* | B-122 | Acuna 17252 (NY-02150575) |
| *Buxus brevipes* | B-634 | Brother Alain 2305 (US-2284282) |
| *Buxus calcarea* | B-426 | Andriamihajarivo 507 (MO-6298213) |
| *Buxus capuronii* | B-425 | Capuron 24212-SF (MO-5730160) |
| *Buxus cephalantha shantouensis* | B-1807 | IBSC-0754341 |
| *Buxus chaoanensis* | B-1808 | IBSC-0345394 |
| *Buxus cipolinica* | B-205 | Birkinshaw 1399 (PRE) |
| *Buxus cipolinica* | B-206 | Birkinshaw 1399 (PRE) |
| *Buxus citrifolia* | B-623 | Bordenave 8253 (US-3523221) |
| *Buxus cochinchinensis* | B-1818 | Larsen 31272 (PE-00044005) |
| *Buxus cochinchinensis* | B-1902 | Shimizu 7957 (TI) |
| *Buxus cochinchinensis* | B-621 | Pot 2488 (US-2589403) |
| *Buxus cochinchinensis* | B-809 | Soejarto 14857 (F-2300811) |
| *Buxus cordata* | B-1602 | Mwangoka 1274 (BRIT) |
| *Buxus cordata* | B-200 | Mwasumbi 2505 (PRE) |
| *Buxus cordata* | B-422 | Mwangoka 1274 (MO-5584102) |
| *Buxus crassifolia* | B-100 | Sattler C732002 (NY-01381373) |
| *Buxus excisa* | B-114 | Ekman 3809 (NY) |
| *Buxus flaviramea* | B-107 | Greuter s.n. (NY-01381374) |
| *Buxus foliosa* | B-106 | Shafer 4073 (NY-02150593) |
| *Buxus glomerata* | B-631 | Garcia 3618 (US-3325282) |
| *Buxus glomerata* | B-711 | Ekman 9477 (UC-996186) |
| *Buxus gonoclada* | B-632 | Morton 8778 (US-2284934) |
| *Buxus gonoclada* | B-803 | Ekman 18585 (F-1474623) |
| *Buxus hainanensis* | B-717 | Lau 5437 (CAS-728942) |
| *Buxus harlandii* | B-620 | Hu 6560 (US-2697068) |
| *Buxus henryi* | B-718 | Xiao 3357 (CAS-1148720) |
| *Buxus heterophylla* | B-119 | Ekman 7555 (NY) |
| *Buxus hildebrandtii* | B-808 | Burger 2978 (F-1632789) |
| *Buxus humbertii* | B-409 | Ratovoson 1539 (MO-6426683) |
| *Buxus ichagensis* | B-1815 | PE-00044079 |
| *Buxus ichagensis* | B-1816 | PE-00044075 |
| *Buxus imbricata* | B-710 | Ekman 15979 (UC-996190) |
| *Buxus itremoensis* | B-407 | Shatz 3980 (MO-5814397) |
| *Buxus itremoensis* | B-408 | Randrianaivo 154 (MO-5206210) |
| *Buxus laevigata* | B-123 | Harris 10755 (NY-02150672) |
| *Buxus lancifolia* | B-713b | Purpus 5304 (UC-155266) |
| *Buxus latistyla* | B-1810 | IBSC-0705548 |
| *Buxus latistyla* | B-1814 | PE-00044093 |
| *Buxus leonii* | B-121 | Brother Leon 4874 (NY) |
| *Buxus linearifolia* | B-1809 | IBSC-0345454 |
| *Buxus liukuensis* | B-619 | Elliott 609 (US-2594126) |
| *Buxus macowanii* | B-207 | No voucher, cultivated, photo in repository |
| *Buxus macowanii* | B-612 | Hardy 5337 (US-2999716) |
| *Buxus macrocarpa* | B-406 | Rabe 83 (MO-6574040) |
| *Buxus macrophylla* | B-630 | Proctor 11819 (US-2585569) |
| *Buxus madagascariensis* | B-412 | Razakamalala 2653 (MO-6426702) |
| *Buxus madagascariensis* | B-413 | Rabehevitra 1117 (MO-6039950) |
| *Buxus madagascariensis* | B-611 | Rabevohitra 3799 (US-3596205) |
| *Buxus marginalis* | B-629 | Axelrod 10410 (US-3397787) |
| *Buxus megistophylla* | B-1901 | Taam 535 (TI) |
| *Buxus mexicana* | B-712b | Purpus 2972 (UC-112959) |
| *Buxus mexicana* | B-723 | Salinas 7524 (CAS-895786) |
| *Buxus microphylla* | B-618 | Moran 4286 (US-2186325) |
| *Buxus microphylla* | B-704 | Bartholomew 233 (UC-1491018) |
| *Buxus moctezumae* | B-724 | Zamudio 11702 (CAS-1027047) |
| *Buxus mollicula* | B-503 | Hu 13051 (HUH) |
| *Buxus monticola* | B-112 | Razanatsima 123 (NY) |
| *Buxus monticola* | B-410 | Capuron 23522-SF (MO-5727647) |
| *Buxus monticola* | B-411 | Razanatsima 123 (MO-6053339) |
| *Buxus moratii* | B-418 | De Block 2209 (MO-6301378) |
| *Buxus moratii* | B-610 | Gillespie 4112 (US-3225197) |
| *Buxus muelleriana* | B-505 | Brother Alain 5451 (HUH) |
| *Buxus myrica* | B-616 | Wang 33332 (US-1670146) |
| *Buxus myrica* | B-706 | Gressitt 1305 (UC-1352900) |
| *Buxus nipensis* | B-113 | Shafer 3219 (NY-02150556) |
| *Buxus niasica* | B-204 | Chapman 8828 (PRE) |
| *Buxus obovata* | B-110 | Ekman 15953 (NY-02150588) |
| *Buxus obtusifolia* | B-810 | Faden 77/647 (F-2155753) |
| *Buxus olivacea* | B-709 | Ekman 15218 (UC-996194) |
| *Buxus olivacea* | B-817 | Ekman 4992 (F-1474625) |
| *Buxus papillosa* | B-617 | Rechinger 29820 (US-2637743) |
| *Buxus papillosa* | B-705 | Rechinger 29820 (UC-1376594) |
| *Buxus pilosula* | B-111 | Brother Leon 19897 (NY-02150633) |
| *Buxus portoricensis* | B-816 | Brother Alain 9811 (F-1618398) |
| *Buxus pubescens* | B-636 | Ferris 5676 (US-1269103) |
| *Buxus pubiramea* | B-1805 | IBSC-0345586 |
| *Buxus pulchella* | B-109 | Britton 4150 (NY-02150671) |
| *Buxus purdieana* | B-802 | Harris 8964 (F-212020) |
| *Buxus rabenantoandroi* | B-414 | Razakamalala 4530 (MO-6426685) |
| *Buxus retusa* | B-105 | Abbott 18972 (NY-01381392) |
| *Buxus retusa* | B-502 | Brother Alain 3668 (HUH) |
| *Buxus revoluta* | B-104 | Berazain s.n. (NY-01381393) |
| *Buxus rheedioides* | B-117 | Ekman 15175 (NY) |
| *Buxus rolfei* | B-614 | Larsen 33973 (US-3376317) |
| *Buxus rotundifolia* | B-115 | Shafer 4114 (NY) |
| *Buxus rugulosa* | B-721 | Boufford 2884 (CAS-1011214) |
| *Buxus sempervirens* | B-003 | No voucher, cultivated, photo in repository |
| *Buxus sempervirens hyrcana* | B-302 | No voucher, photo in repository |
| *Buxus shaferi* | B-627 | Brother Alain 4761 (US-2288281) |
| *Buxus sinica* | B-720 | Yuan 1086 (CAS-846679) |
| *Buxus sirindhorniana* | B-500 | van de Bult 664 (HUH) |
| *Buxus* sp. *celebica* | B-1817 | PE-00044002 |
| *Buxus* sp. Philippines | B-506 | Reynoso 14113 (HUH) |
| *Buxus stenophylla* | B-1903 | Hu 13026 (TI) |
| *Buxus subcolumnaris* | B-626 | Howard 18878 (US-3001023) |
| *Buxus vaccinioides* | B-116 | Shafer 4122 (NY) |
| *Buxus vaccinioides* | B-126 | Shafer 4122 (NY) |
| *Buxus vahlii* | B-625 | Little 21745 (US-2673754) |
| *Buxus vahlii* | B-807 | Brother Alain 10722 (F-1618353) |
| *Buxus wallichiana* | B-615 | Koelz 1831 (US-1605756) |
| *Didymeles integrifolia* | B-602 | Schatz 2778 (US-3627925) |
| *Didymeles madagascariensis* | B-601 | Rakotomazala 399 (US-3350972) |
| *Didymeles perrieri* | B-415 | Razafimandribison 119 (MO-5617933) |
| *Didymeles* sp. Toamasina | B-417 | Antilahimena 1529 (MO-6144923) |
| *Didymeles* sp. Toamasina | B-423 | Antilahimena 2497a (MO-6177345) |
| *Pachysandra axillaris* | B-639 | Hsien 1531 (US-1968483) |
| *Pachysandra axillaris* | B-703 | Bonati s.n. (UC) |
| *Pachysandra axillaris* | B-714 | Liu 15428 (CAS-943872) |
| *Pachysandra procumbens* | B-640 | Kral 34 119 (US-2674034) |
| *Pachysandra terminalis* | B-005 | No voucher, cultivated, photo in repository |
| *Sarcococca balansae* | B-303 | No voucher, photo in repository |
| *Sarcococca balansae* | B-649 | How 70661 (US-1675664) |
| *Sarcococca balansae* | B-811 | How 70661 (F-779258) |
| *Sarcococca bleddynii* | floden3847 | Floden s.n. (TENN) |
| *Sarcococca confusa* | B-713a | McClintock s.n. (CAS-582464) |
| *Sarcococca konzattii* | B-651 | Panero 5574 (US-3661245) |
| *Sarcococca konzattii* | floden3834 | Calzada 19727 (TENN) |
| *Sarcococca konzattii* | floden3835 | Calzada 19727 (TENN) |
| *Sarcococca konzattii* | floden4073 | Skutch 553 (TX) |
| *Sarcococca coriacea* | B-648 | Nocolson 2712 (US-2571553) |
| *Sarcococca coriacea* | B-650 | Bostick s.n. (US-29760003) |
| *Sarcococca euphlebia* | B-1801 | IBSC-0757158 |
| *Sarcococca hookeriana* | B-004 | No voucher, cultivated, photo in repository |
| *Sarcococca hookeriana* | B-646 | Bartholomew 1446 (US-3043590) |
| *Sarcococca longipetiolata* | B-1804 | IBSC-0347445 |
| *Sarcococca longipetiolata* | B-1812 | PE-00055950 |
| *Sarcococca orientalis* | B-1811 | PE-01841407 |
| *Sarcococca philippinensis* | B-2001 | Benguel 4645 (M) |
| *Sarcococca ruscifolia* | B-645 | Rock 11598 (US-1333178) |
| *Sarcococca ruscifolia* | B-701 | Bartholomew 1912 (UC-1491879) |
| *Sarcococca ruscifolia* | B-712a | Boufford 37718 (CAS-1103639) |
| *Sarcococca saligna* | B-644 | Nasir 397 (US-2395364) |
| *Sarcococca saligna* | floden3971 | Floden s.n. (TENN) |
| *Sarcococca* sp. Myanmar | B-3005 | Hidetoshi 035324 (KYO) |
| *Sarcococca wallichii* | B-643 | Bartholomew 688 (US-3068530) |
| *Sarcococca zeylanica* | B-642 | Huber 874 (US-2868259) |
| *Sarcococca zeylanica* | B-647 | Saldanha 10489 (US-2530653) |
| *Styloceras brokawii* | B-2101 | Nee 52239 (SPF-168130) |
| *Styloceras brokawii* | B-2151 | Nee 50191 (SP-350205) |
| *Styloceras brokawii* | B-606 | Nee 39225 (US-3250366) |
| *Styloceras columnare* | B-1604 | Lewis 881062 (BRIT) |
| *Styloceras columnare* | B-605 | Cardenas 52 (US-2325019) |
| *Styloceras connatum* | B-401 | Macia 3993 (MO-5905935) |
| *Styloceras connatum* | B-402 | Maldonado 2688 (MO-5734632) |
| *Styloceras kunthianum* | B-2801 | Altemirano 157 (QCA-26086) |
| *Styloceras kunthianum* | B-403 | van der Werff 10955 (MO-3657438) |
| *Styloceras kunthianum* | B-404 | Rubio 2224 (MO-4293135) |
| *Styloceras kunthianum* | B-818 | Bonpland 9208 (F-1012222) |
| *Styloceras laurifolium* | B-1603 | Lewis 37398 (BRIT) |
| *Styloceras laurifolium* | B-301 | No voucher, cultivated, photo in repository |
| *Styloceras laurifolium* | B-400 | Peyton 1083 (MO-3012831) |
| *Styloceras laurifolium* | B-603 | Vargas 15578 (US-2438721) |
| *Styloceras laurifolium* | B-815 | Young 4328 (F-1980936) |
| \_shipunov\_floden\_et\_al\_buxineae |  |  |
| Styloceras penninervium | B-427 | Smith 3206 (MO-3395324) |
| Styloceras sp. Ecuador | B-604 | Clark 8910 (US-3500727) |
| Styloceras sp. Peru Oxapampa | B-2202 | Valenzuela 13978 (USM-252811) |
| Trochodendron aralioides | B-001 | No voucher, cultivated, photo in repository |
| Trochodendron aralioides | B-002 | No voucher, cultivated, photo in repository |

| **Region** | **Sequenced fragment** |
| --- | --- |
| Gabon | rbcl |
| Ivory Coast | rbcl, trnl |
| Gabon | rbcl |
| Cuba | rbcl, trnl |
| Jamaica | its, rbcl, trnl |
| Yunnan | its, rbcl, trnl |
| Bahamas | rbcl |
| Bahamas | trnl |
| Mallorca | its |
| Mexico | rbcl, trnl |
| Angola | its, rbcl, trnl |
| Cuba | rbcl |
| Guanxi | its, rbcl, trnl |
| Cuba | trnl |
| Cuba | rbcl, trnl |
| Cuba | its, rbcl, trnl |
| Cuba | rbcl |
| Madagascar | its, rbcl, trnl |
| Madagascar | its, rbcl, trnl |
| Guangdong | its, rbcl, trnl |
| Guangdong | its, rbcl, trnl |
| Madagascar | rbcl, trnl |
| Madagascar | its, rbcl, trnl |
| Suriname | rbcl, trnl |
| Thailand | rbcl |
| Thailand | rbcl |
| Thailand | rbcl, trnl |
| Vietnam | rbcl |
| Tanzania | rbcl, trnl |
| Tanzania | rbcl, trnl |
| Tanzania | rbcl, trnl |
| Cuba | its, rbcl, trnl |
| Cuba | rbcl, trnl |
| Cuba | rbcl, trnl |
| Cuba | rbcl, trnl |
| Hispaniola | its, rbcl, trnl |
| Cuba | rbcl, trnl |
| Cuba | trnl |
| Cuba | its, rbcl, trnl |
| Hainan | rbcl, trnl |
| Hong Kong | rbcl |
| Hunan | rbcl, trnl |
| Cuba | trnl |
| Ethiopia | rbcl |
| Madagascar | its, rbcl, trnl |
| Hubei | its, rbcl |
| Shaanxi | its, rbcl |
| Cuba | its, rbcl, trnl |
| Madagascar | its, rbcl, trnl |
| Madagascar | its, rbcl, trnl |
| Cuba | rbcl, trnl |
| Mexico | its, rbcl, trnl |
| Guanxi | its, rbcl |
| Guanxi | its, rbcl, trnl |
| Cuba | its, rbcl, trnl |
| Guanxi | rbcl |
| Okinawa | its, rbcl |
| Cape | its, rbcl, trnl |
| South Africa | rbcl, trnl |
| Madagascar | its, rbcl, trnl |
| Jamaica | rbcl, trnl |
| Madagascar | its, rbcl |
| Madagascar | its, rbcl, trnl |
| Madagascar | its, rbcl |
| Cuba | rbcl, trnl |
| Guangdong | its, rbcl, trnl |
| Mexico | its, rbcl, trnl |
| Mexico | its, rbcl, trnl |
| Korea | rbcl |
| Hubei | rbcl, trnl |
| Mexico | rbcl, trnl |
| Hong Kong | its, rbcl, trnl |
| Madagascar | its, rbcl, trnl |
| Madagascar | rbcl |
| Madagascar | its, rbcl, trnl |
| Madagascar | rbcl, trnl |
| Madagascar | its, rbcl, trnl |
| Cuba | rbcl |
| Hainan | rbcl, trnl |
| Hainan | its, rbcl, trnl |
| Cuba | rbcl |
| Malawi | its, rbcl |
| Cuba | trnl |
| Kenya | rbcl, trnl |
| Cuba | rbcl |
| Cuba | rbcl |
| Pakistan | its, trnl |
| Pakistan | its, trnl |
| Cuba | its, rbcl, trnl |
| Puerto Rico | trnl |
| Mexico | its, rbcl, trnl |
| Hainan | rbcl |
| Jamaica | rbcl |
| Jamaica | rbcl, trnl |
| Madagascar | its, rbcl |
| Cuba | rbcl, trnl |
| Cuba | trnl |
| Cuba | its, rbcl, trnl |
| Cuba | its, rbcl, trnl |
| Thailand | its, rbcl, trnl |
| Cuba | rbcl, trnl |
| Sichuan | its, rbcl, trnl |
| USA | its |
| Azerbaijan | its, rbcl, trnl |
| Cuba | its, rbcl, trnl |
| Gansu | its, rbcl, trnl |
| Thailand | its, rbcl, trnl |
| China | rbcl |
| Sibuyan | its, rbcl |
| Hong Kong | its, rbcl, trnl |
| Martinique | its, rbcl, trnl |
| Cuba | rbcl |
| Cuba | its, rbcl, trnl |
| Puerto Rico | its, trnl |
| Puerto Rico | rbcl, trnl |
| Himachal Pradesh | its, rbcl, trnl |
| Madagascar | rbcl |
| Madagascar | its, rbcl, trnl |
| Madagascar | its, rbcl, trnl |
| Madagascar | rbcl |
| Madagascar | its |
| Sichuan | its, trnl |
| Yunnan | its |
| Sichuan | its, trnl |
| Tennessee | its, trnl |
| USA | its, rbcl |
| Vietnam | its, rbcl |
| Hainan | rbcl, trnl |
| Hainan | rbcl |
| Vietnam | its, rbcl, trnl |
| Cultivated | its, rbcl, trnl |
| Mexico | rbcl, trnl |
| Mexico | rbcl, trnl |
| Mexico | its, trnl |
| Guatemala | its, trnl |
| Nepal | its, rbcl |
| Cultivated | its, rbcl, trnl |
| Hainan | its, rbcl, trnl |
| USA | its, rbcl, trnl |
| Yunnan | its, rbcl, trnl |
| Guangdong | rbcl |
| Hunan | its, rbcl, trnl |
| Guanxi | its, rbcl, trnl |
| Luzon | rbcl, trnl |
| Yunnan | its, rbcl, trnl |
| Hubei | its, rbcl, trnl |
| Gansu | its, rbcl, trnl |
| Pakistan | its, rbcl, trnl |
| NA | its, trnl |
| Myanmar | its, rbcl |
| Yunnan | its, rbcl, trnl |
| Ceylon | its, rbcl |
| Tamil Nadu | its, rbcl, trnl |
| Bolivia | rbcl, trnl |
| Bolivia | rbcl |
| Bolivia | rbcl |
| Bolivia | rbcl, trnl |
| Bolivia | its, rbcl |
| Bolivia | its, rbcl |
| Bolivia | its |
| Ecuador | rbcl, trnl |
| Ecuador | its, rbcl |
| Ecuador | trnl |
| Ecuador | its, trnl |
| Bolivia | rbcl, trnl |
| Colombia | its, trnl |
| Peru | its |
| Peru | its, rbcl, trnl |
| Peru | trnl |
| Peru | its, rbcl, trnl |
| Ecuador | rbcl |
| Peru | rbcl |
| USA | its |
| USA | its |

**Support Table 3.**
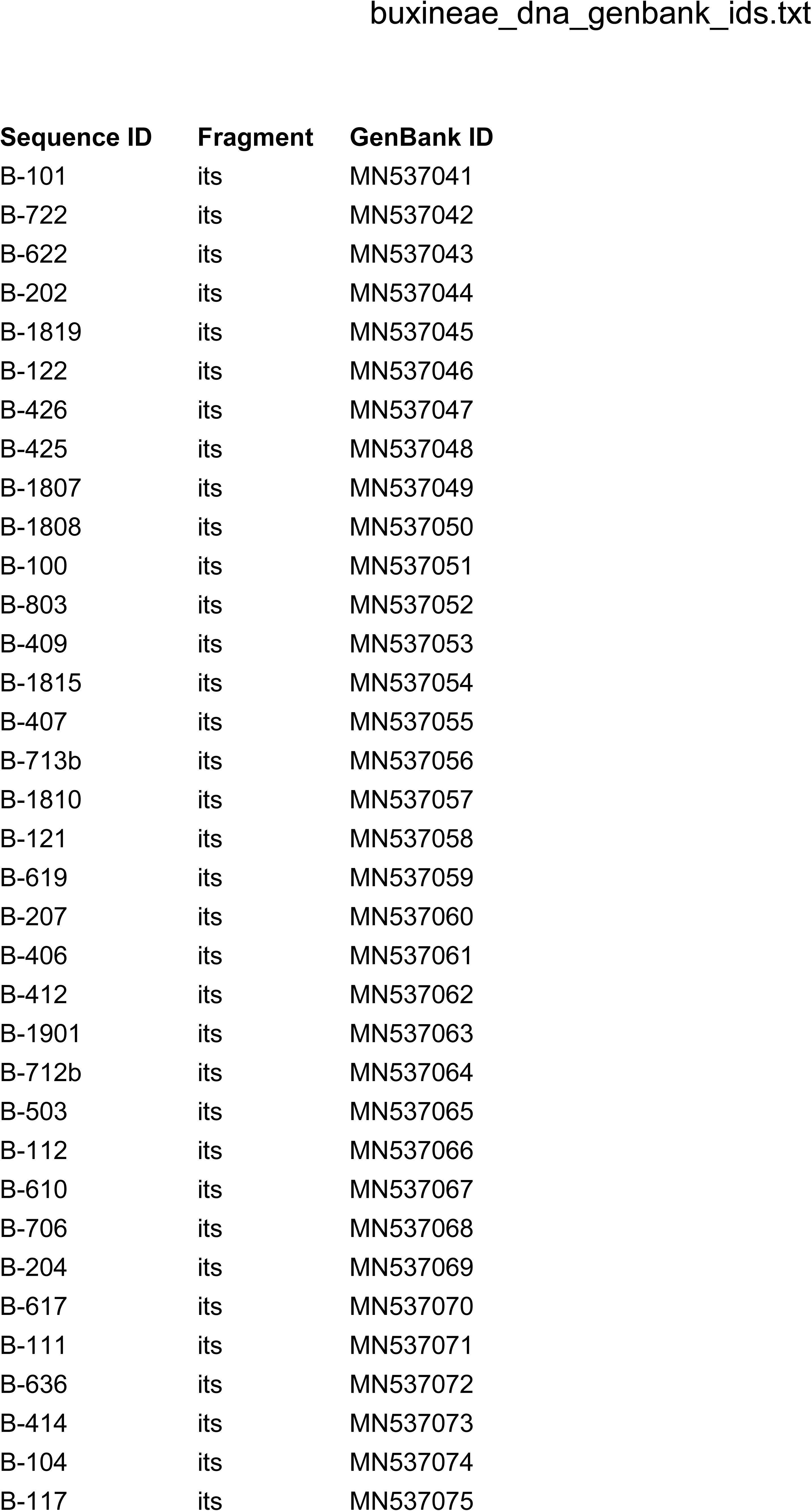
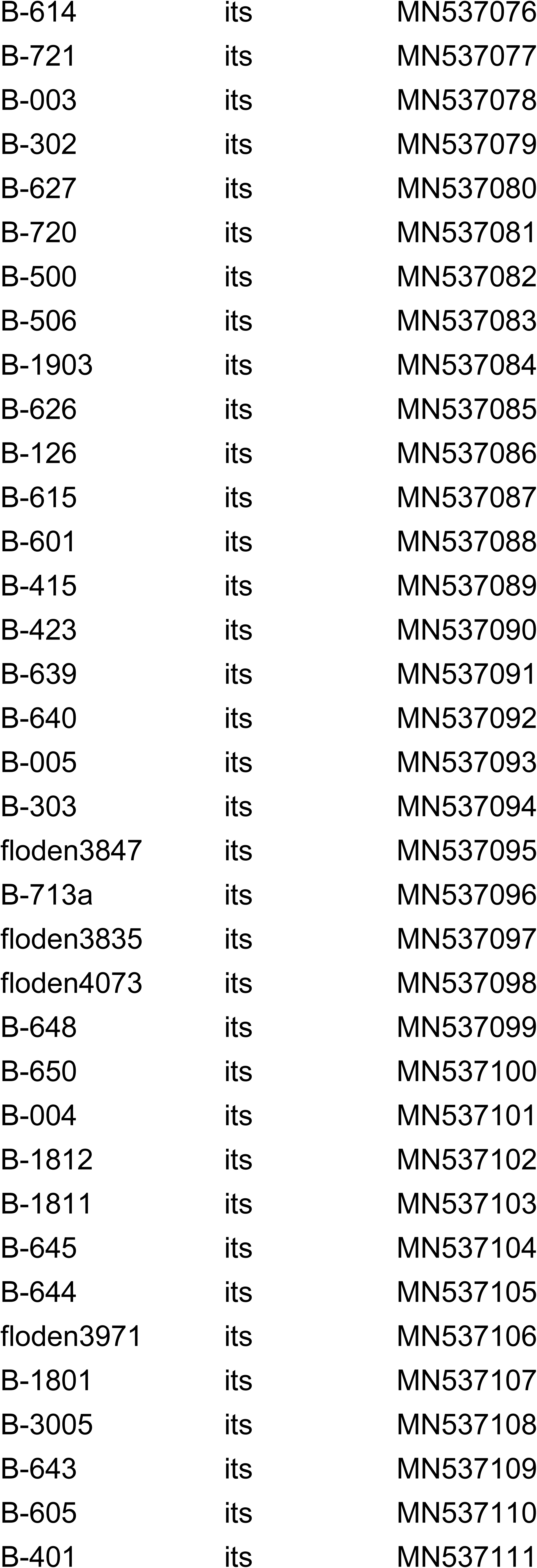
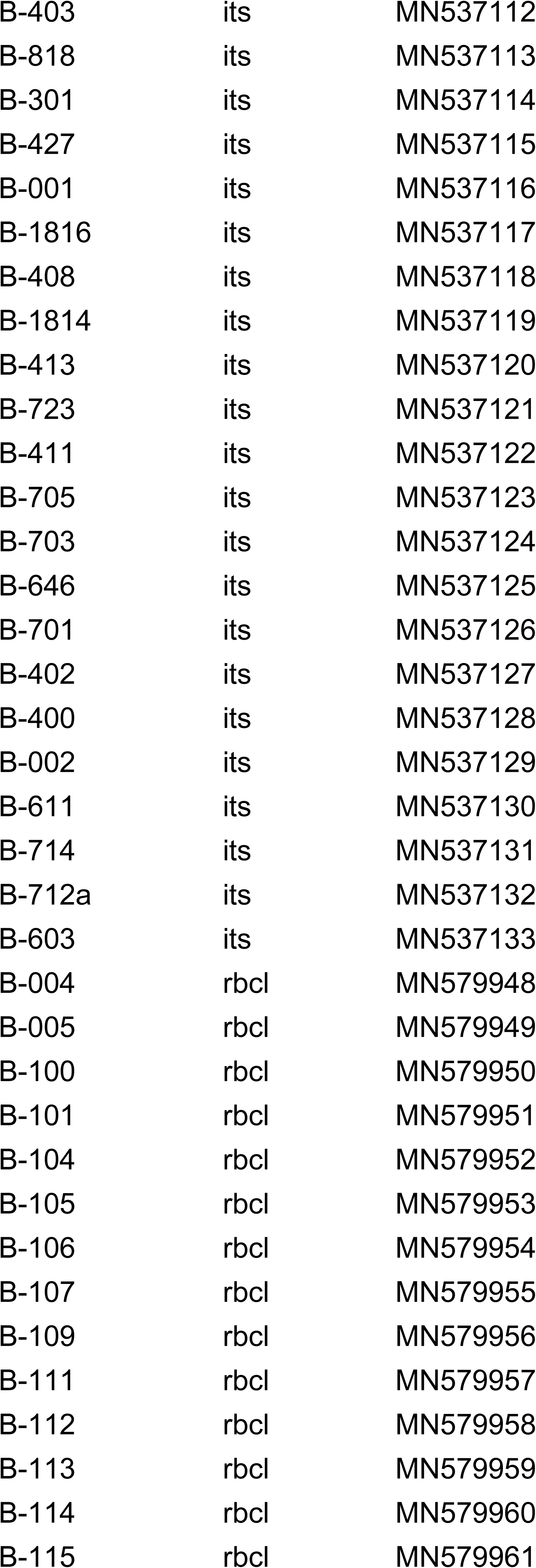
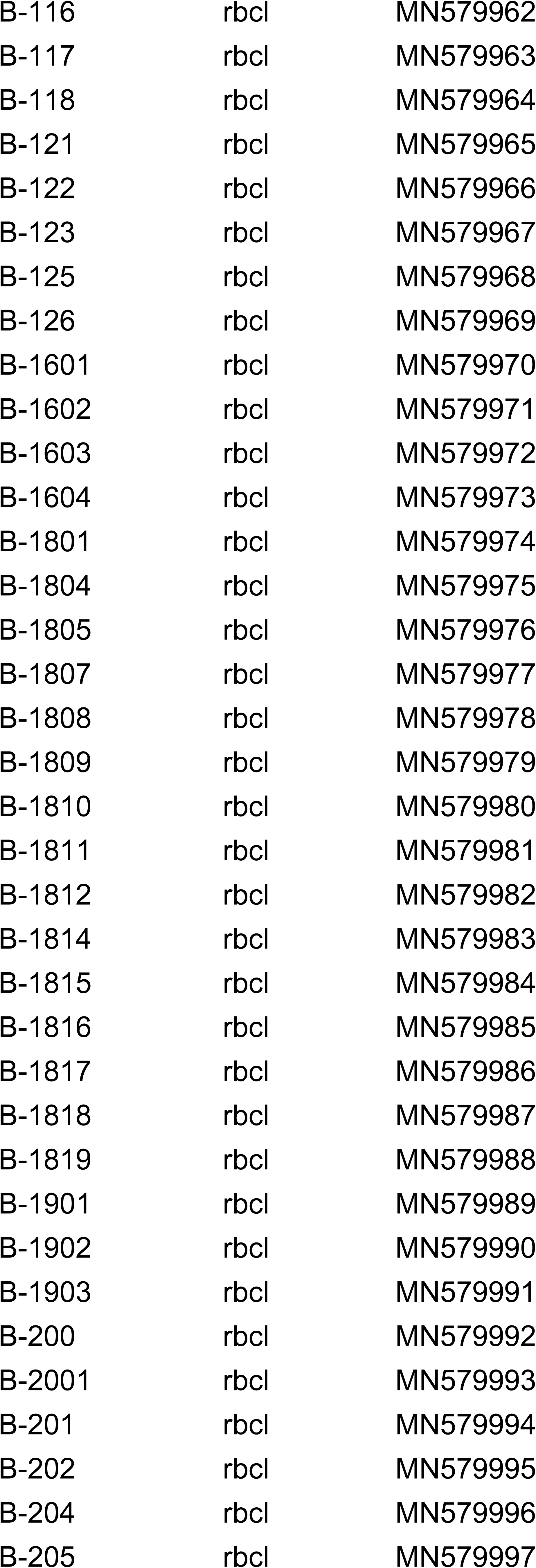
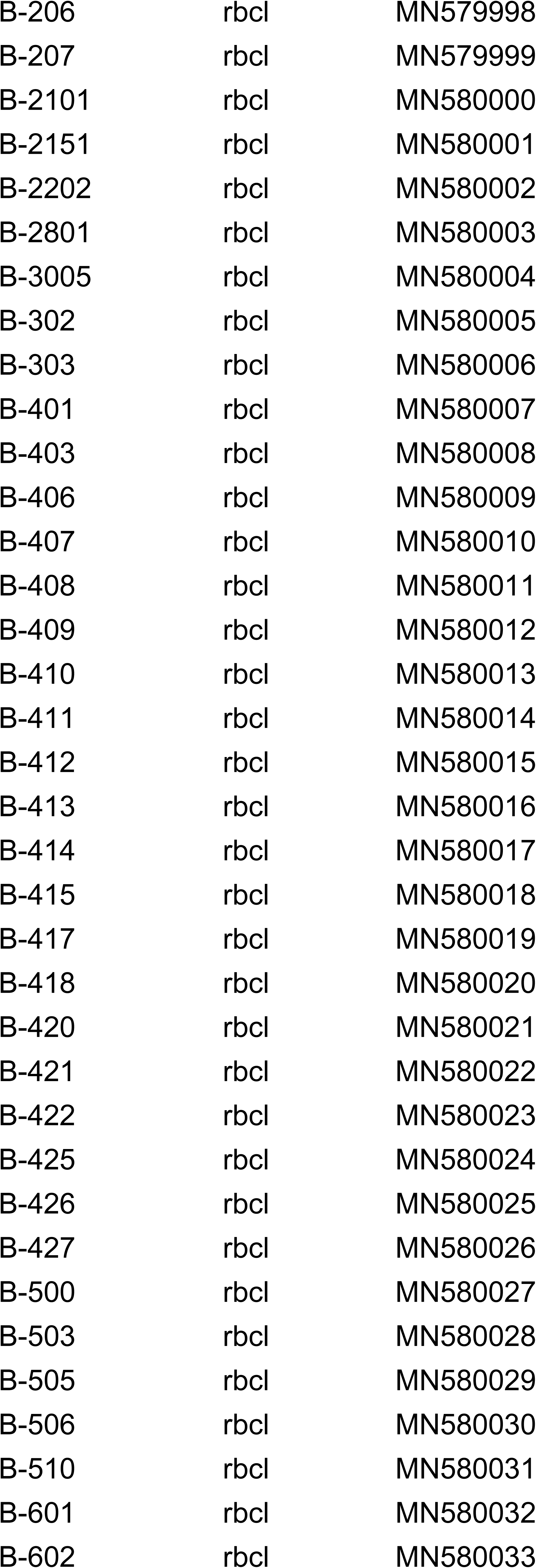
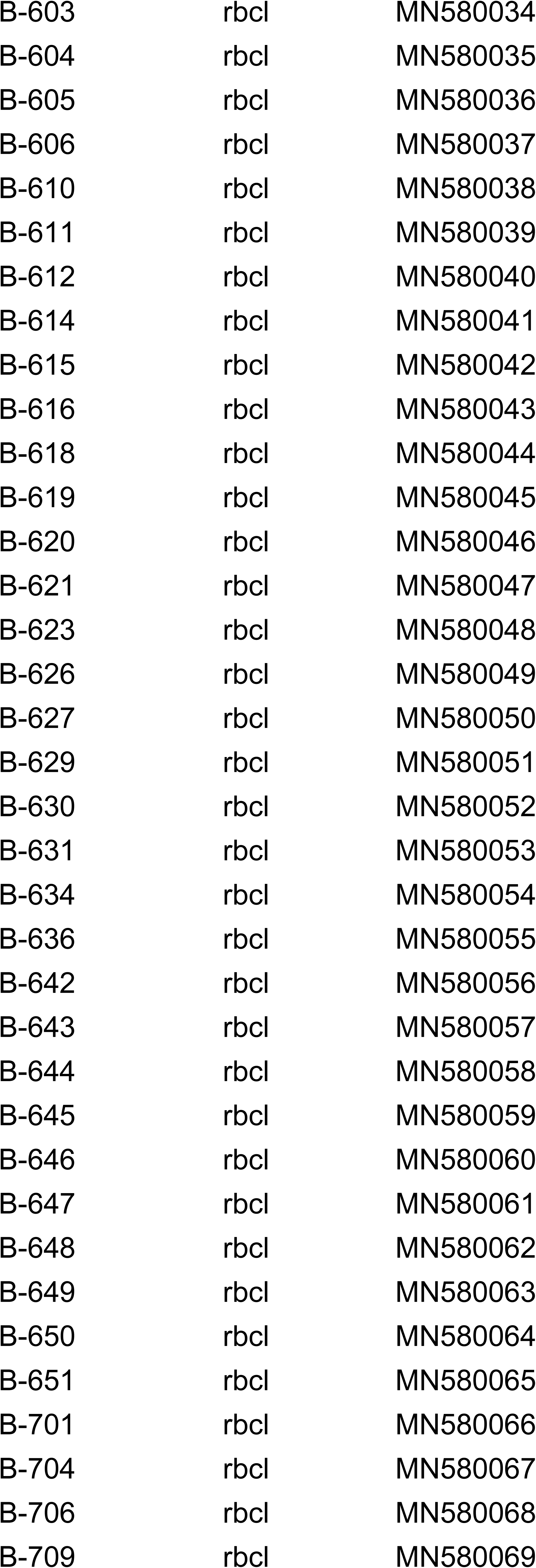
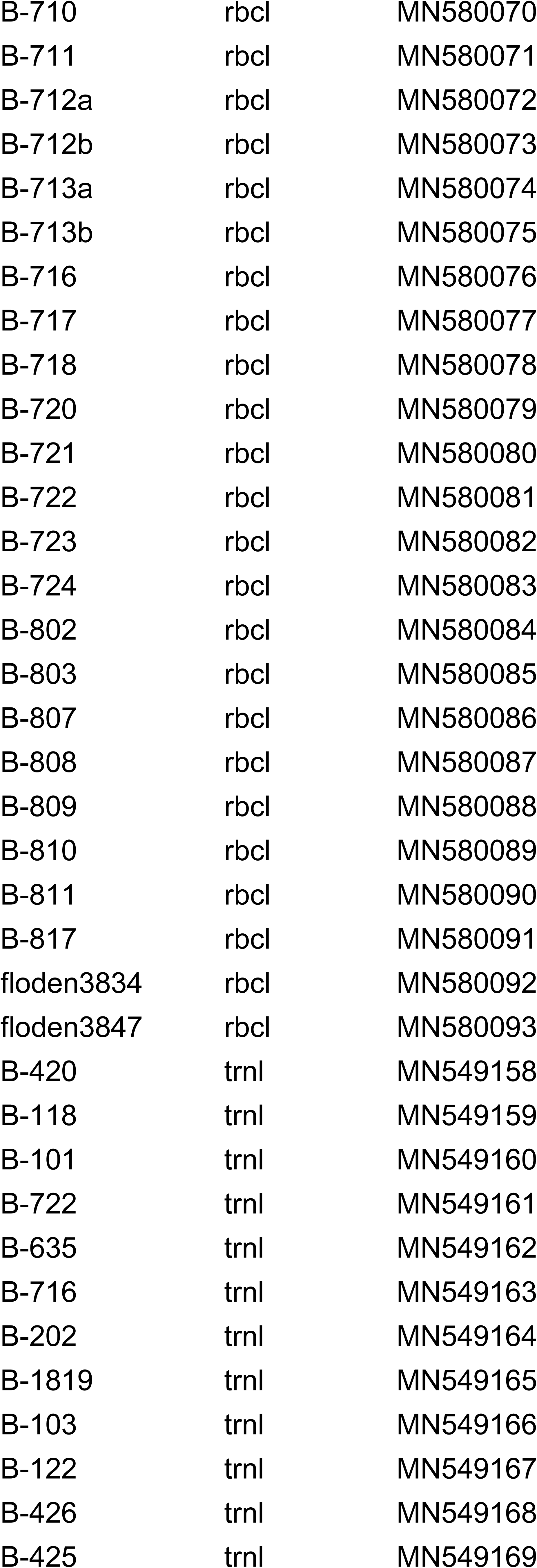
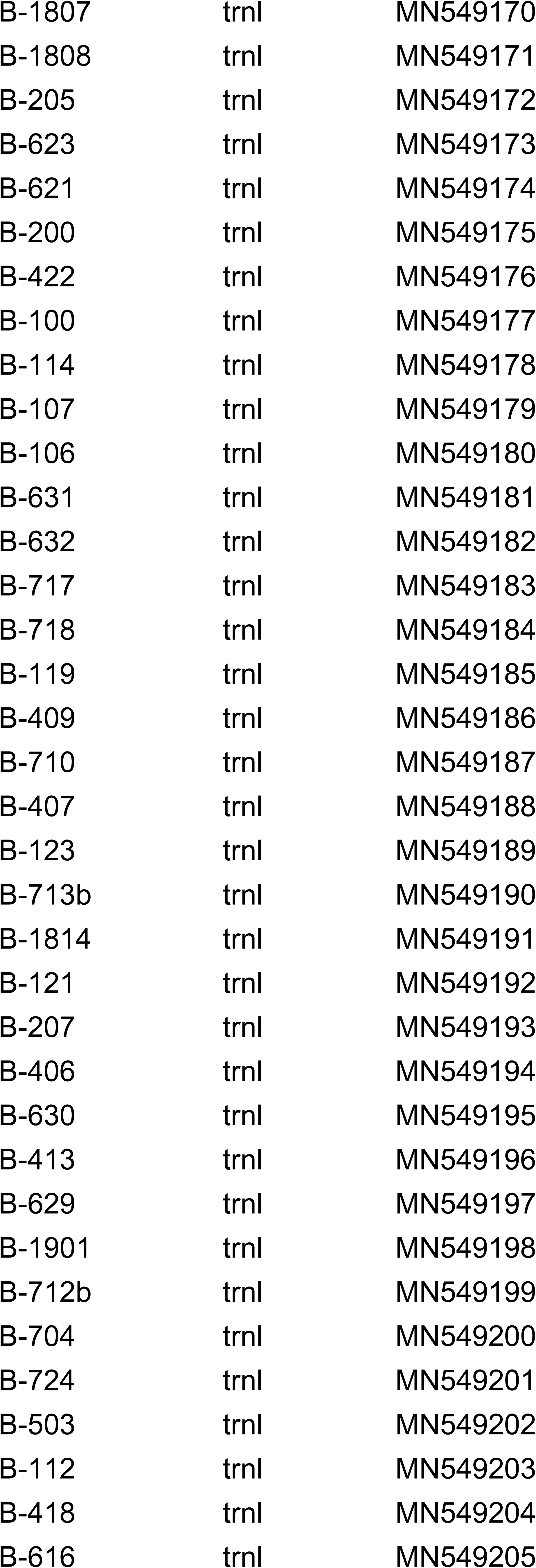
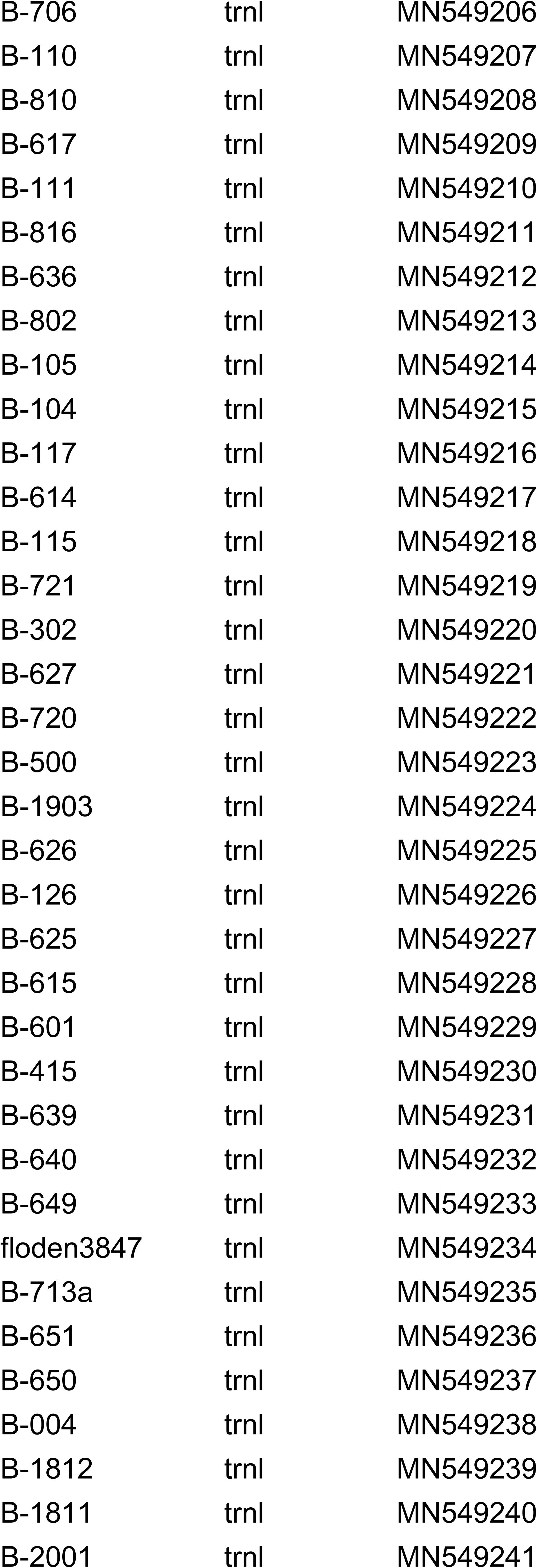
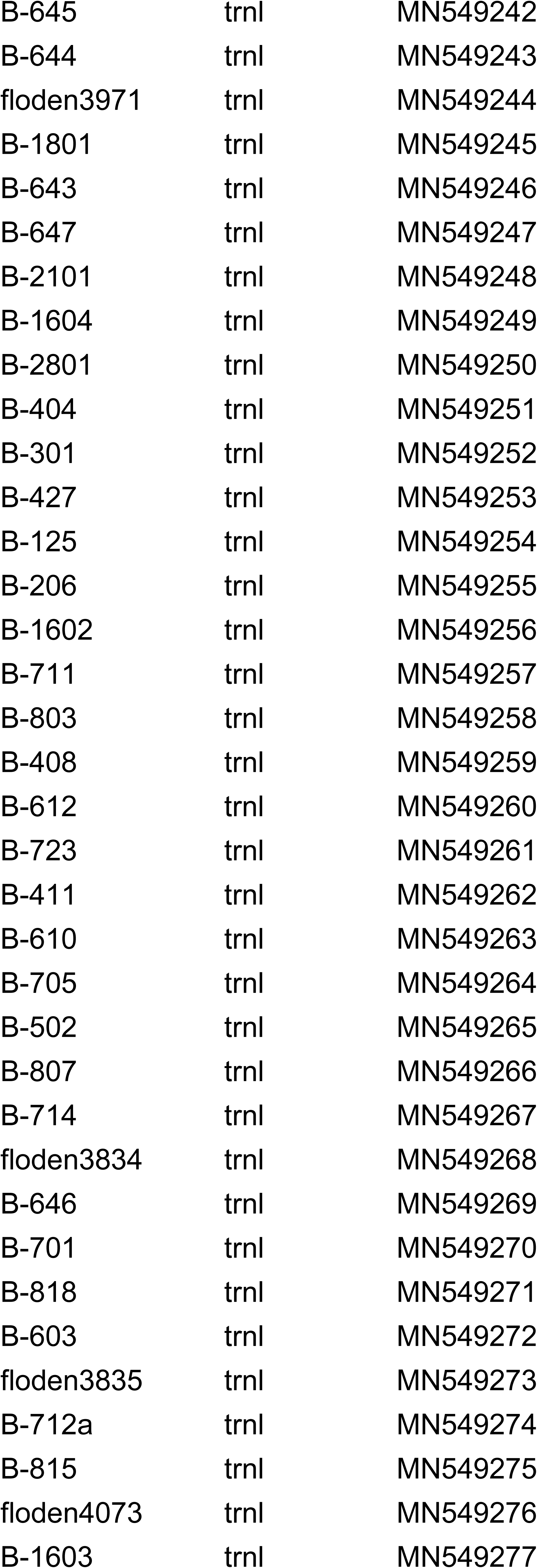
GenBank accession numbers of Buxaceae samples sequenced for this study.

buxineae\_dna\_genbank\_ids.txt
| Sequence ID | Fragment | GenBank ID |
| --- | --- | --- |
| B-101 | its | MN537041 |
| B-722 | its | MN537042 |
| B-622 | its | MN537043 |
| B-202 | its | MN537044 |
| B-1819 | its | MN537045 |
| B-122 | its | MN537046 |
| B-426 | its | MN537047 |
| B-425 | its | MN537048 |
| B-1807 | its | MN537049 |
| B-1808 | its | MN537050 |
| B-100 | its | MN537051 |
| B-803 | its | MN537052 |
| B-409 | its | MN537053 |
| B-1815 | its | MN537054 |
| B-407 | its | MN537055 |
| B-713b | its | MN537056 |
| B-1810 | its | MN537057 |
| B-121 | its | MN537058 |
| B-619 | its | MN537059 |
| B-207 | its | MN537060 |
| B-406 | its | MN537061 |
| B-412 | its | MN537062 |
| B-1901 | its | MN537063 |
| B-712b | its | MN537064 |
| B-503 | its | MN537065 |
| B-112 | its | MN537066 |
| B-610 | its | MN537067 |
| B-706 | its | MN537068 |
| B-204 | its | MN537069 |
| B-617 | its | MN537070 |
| B-111 | its | MN537071 |
| B-636 | its | MN537072 |
| B-414 | its | MN537073 |
| B-104 | its | MN537074 |
| B-117 | its | MN537075 |
| B-614 | its | MN537076 |
| B-721 | its | MN537077 |
| B-003 | its | MN537078 |
| B-302 | its | MN537079 |
| B-627 | its | MN537080 |
| B-720 | its | MN537081 |
| B-500 | its | MN537082 |
| B-506 | its | MN537083 |
| B-1903 | its | MN537084 |
| B-626 | its | MN537085 |
| B-126 | its | MN537086 |
| B-615 | its | MN537087 |
| B-601 | its | MN537088 |
| B-415 | its | MN537089 |
| B-423 | its | MN537090 |
| B-639 | its | MN537091 |
| B-640 | its | MN537092 |
| B-005 | its | MN537093 |
| B-303 | its | MN537094 |
| floden3847 | its | MN537095 |
| B-713a | its | MN537096 |
| floden3835 | its | MN537097 |
| floden4073 | its | MN537098 |
| B-648 | its | MN537099 |
| B-650 | its | MN537100 |
| B-004 | its | MN537101 |
| B-1812 | its | MN537102 |
| B-1811 | its | MN537103 |
| B-645 | its | MN537104 |
| B-644 | its | MN537105 |
| floden3971 | its | MN537106 |
| B-1801 | its | MN537107 |
| B-3005 | its | MN537108 |
| B-643 | its | MN537109 |
| B-605 | its | MN537110 |
| B-401 | its | MN537111 |
| B-403 | its | MN537112 |
| B-818 | its | MN537113 |
| B-301 | its | MN537114 |
| B-427 | its | MN537115 |
| B-001 | its | MN537116 |
| B-1816 | its | MN537117 |
| B-408 | its | MN537118 |
| B-1814 | its | MN537119 |
| B-413 | its | MN537120 |
| B-723 | its | MN537121 |
| B-411 | its | MN537122 |
| B-705 | its | MN537123 |
| B-703 | its | MN537124 |
| B-646 | its | MN537125 |
| B-701 | its | MN537126 |
| B-402 | its | MN537127 |
| B-400 | its | MN537128 |
| B-002 | its | MN537129 |
| B-611 | its | MN537130 |
| B-714 | its | MN537131 |
| B-712a | its | MN537132 |
| B-603 | its | MN537133 |
| B-004 | rbcl | MN579948 |
| B-005 | rbcl | MN579949 |
| B-100 | rbcl | MN579950 |
| B-101 | rbcl | MN579951 |
| B-104 | rbcl | MN579952 |
| B-105 | rbcl | MN579953 |
| B-106 | rbcl | MN579954 |
| B-107 | rbcl | MN579955 |
| B-109 | rbcl | MN579956 |
| B-111 | rbcl | MN579957 |
| B-112 | rbcl | MN579958 |
| B-113 | rbcl | MN579959 |
| B-114 | rbcl | MN579960 |
| B-115 | rbcl | MN579961 |
| B-116 | rbcl | MN579962 |
| B-117 | rbcl | MN579963 |
| B-118 | rbcl | MN579964 |
| B-121 | rbcl | MN579965 |
| B-122 | rbcl | MN579966 |
| B-123 | rbcl | MN579967 |
| B-125 | rbcl | MN579968 |
| B-126 | rbcl | MN579969 |
| B-1601 | rbcl | MN579970 |
| B-1602 | rbcl | MN579971 |
| B-1603 | rbcl | MN579972 |
| B-1604 | rbcl | MN579973 |
| B-1801 | rbcl | MN579974 |
| B-1804 | rbcl | MN579975 |
| B-1805 | rbcl | MN579976 |
| B-1807 | rbcl | MN579977 |
| B-1808 | rbcl | MN579978 |
| B-1809 | rbcl | MN579979 |
| B-1810 | rbcl | MN579980 |
| B-1811 | rbcl | MN579981 |
| B-1812 | rbcl | MN579982 |
| B-1814 | rbcl | MN579983 |
| B-1815 | rbcl | MN579984 |
| B-1816 | rbcl | MN579985 |
| B-1817 | rbcl | MN579986 |
| B-1818 | rbcl | MN579987 |
| B-1819 | rbcl | MN579988 |
| B-1901 | rbcl | MN579989 |
| B-1902 | rbcl | MN579990 |
| B-1903 | rbcl | MN579991 |
| B-200 | rbcl | MN579992 |
| B-2001 | rbcl | MN579993 |
| B-201 | rbcl | MN579994 |
| B-202 | rbcl | MN579995 |
| B-204 | rbcl | MN579996 |
| B-205 | rbcl | MN579997 |
| B-206 | rbcl | MN579998 |
| B-207 | rbcl | MN579999 |
| B-2101 | rbcl | MN580000 |
| B-2151 | rbcl | MN580001 |
| B-2202 | rbcl | MN580002 |
| B-2801 | rbcl | MN580003 |
| B-3005 | rbcl | MN580004 |
| B-302 | rbcl | MN580005 |
| B-303 | rbcl | MN580006 |
| B-401 | rbcl | MN580007 |
| B-403 | rbcl | MN580008 |
| B-406 | rbcl | MN580009 |
| B-407 | rbcl | MN580010 |
| B-408 | rbcl | MN580011 |
| B-409 | rbcl | MN580012 |
| B-410 | rbcl | MN580013 |
| B-411 | rbcl | MN580014 |
| B-412 | rbcl | MN580015 |
| B-413 | rbcl | MN580016 |
| B-414 | rbcl | MN580017 |
| B-415 | rbcl | MN580018 |
| B-417 | rbcl | MN580019 |
| B-418 | rbcl | MN580020 |
| B-420 | rbcl | MN580021 |
| B-421 | rbcl | MN580022 |
| B-422 | rbcl | MN580023 |
| B-425 | rbcl | MN580024 |
| B-426 | rbcl | MN580025 |
| B-427 | rbcl | MN580026 |
| B-500 | rbcl | MN580027 |
| B-503 | rbcl | MN580028 |
| B-505 | rbcl | MN580029 |
| B-506 | rbcl | MN580030 |
| B-510 | rbcl | MN580031 |
| B-601 | rbcl | MN580032 |
| B-602 | rbcl | MN580033 |
| B-603 | rbcl | MN580034 |
| B-604 | rbcl | MN580035 |
| B-605 | rbcl | MN580036 |
| B-606 | rbcl | MN580037 |
| B-610 | rbcl | MN580038 |
| B-611 | rbcl | MN580039 |
| B-612 | rbcl | MN580040 |
| B-614 | rbcl | MN580041 |
| B-615 | rbcl | MN580042 |
| B-616 | rbcl | MN580043 |
| B-618 | rbcl | MN580044 |
| B-619 | rbcl | MN580045 |
| B-620 | rbcl | MN580046 |
| B-621 | rbcl | MN580047 |
| B-623 | rbcl | MN580048 |
| B-626 | rbcl | MN580049 |
| B-627 | rbcl | MN580050 |
| B-629 | rbcl | MN580051 |
| B-630 | rbcl | MN580052 |
| B-631 | rbcl | MN580053 |
| B-634 | rbcl | MN580054 |
| B-636 | rbcl | MN580055 |
| B-642 | rbcl | MN580056 |
| B-643 | rbcl | MN580057 |
| B-644 | rbcl | MN580058 |
| B-645 | rbcl | MN580059 |
| B-646 | rbcl | MN580060 |
| B-647 | rbcl | MN580061 |
| B-648 | rbcl | MN580062 |
| B-649 | rbcl | MN580063 |
| B-650 | rbcl | MN580064 |
| B-651 | rbcl | MN580065 |
| B-701 | rbcl | MN580066 |
| B-704 | rbcl | MN580067 |
| B-706 | rbcl | MN580068 |
| B-709 | rbcl | MN580069 |
| B-710 | rbcl | MN580070 |
| B-711 | rbcl | MN580071 |
| B-712a | rbcl | MN580072 |
| B-712b | rbcl | MN580073 |
| B-713a | rbcl | MN580074 |
| B-713b | rbcl | MN580075 |
| B-716 | rbcl | MN580076 |
| B-717 | rbcl | MN580077 |
| B-718 | rbcl | MN580078 |
| B-720 | rbcl | MN580079 |
| B-721 | rbcl | MN580080 |
| B-722 | rbcl | MN580081 |
| B-723 | rbcl | MN580082 |
| B-724 | rbcl | MN580083 |
| B-802 | rbcl | MN580084 |
| B-803 | rbcl | MN580085 |
| B-807 | rbcl | MN580086 |
| B-808 | rbcl | MN580087 |
| B-809 | rbcl | MN580088 |
| B-810 | rbcl | MN580089 |
| B-811 | rbcl | MN580090 |
| B-817 | rbcl | MN580091 |
| floden3834 | rbcl | MN580092 |
| floden3847 | rbcl | MN580093 |
| B-420 | trnl | MN549158 |
| B-118 | trnl | MN549159 |
| B-101 | trnl | MN549160 |
| B-722 | trnl | MN549161 |
| B-635 | trnl | MN549162 |
| B-716 | trnl | MN549163 |
| B-202 | trnl | MN549164 |
| B-1819 | trnl | MN549165 |
| B-103 | trnl | MN549166 |
| B-122 | trnl | MN549167 |
| B-426 | trnl | MN549168 |
| B-425 | trnl | MN549169 |
| B-1807 | trnI | MN549170 |
| B-1808 | trnI | MN549171 |
| B-205 | trnI | MN549172 |
| B-623 | trnI | MN549173 |
| B-621 | trnI | MN549174 |
| B-200 | trnI | MN549175 |
| B-422 | trnI | MN549176 |
| B-100 | trnI | MN549177 |
| B-114 | trnI | MN549178 |
| B-107 | trnI | MN549179 |
| B-106 | trnI | MN549180 |
| B-631 | trnI | MN549181 |
| B-632 | trnI | MN549182 |
| B-717 | trnI | MN549183 |
| B-718 | trnI | MN549184 |
| B-119 | trnI | MN549185 |
| B-409 | trnI | MN549186 |
| B-710 | trnI | MN549187 |
| B-407 | trnI | MN549188 |
| B-123 | trnI | MN549189 |
| B-713b | trnI | MN549190 |
| B-1814 | trnI | MN549191 |
| B-121 | trnI | MN549192 |
| B-207 | trnI | MN549193 |
| B-406 | trnI | MN549194 |
| B-630 | trnI | MN549195 |
| B-413 | trnI | MN549196 |
| B-629 | trnI | MN549197 |
| B-1901 | trnI | MN549198 |
| B-712b | trnI | MN549199 |
| B-704 | trnI | MN549200 |
| B-724 | trnI | MN549201 |
| B-503 | trnI | MN549202 |
| B-112 | trnI | MN549203 |
| B-418 | trnI | MN549204 |
| B-616 | trnI | MN549205 |
| B-706 | trnI | MN549206 |
| B-110 | trnI | MN549207 |
| B-810 | trnI | MN549208 |
| B-617 | trnI | MN549209 |
| B-111 | trnI | MN549210 |
| B-816 | trnI | MN549211 |
| B-636 | trnI | MN549212 |
| B-802 | trnI | MN549213 |
| B-105 | trnI | MN549214 |
| B-104 | trnI | MN549215 |
| B-117 | trnI | MN549216 |
| B-614 | trnI | MN549217 |
| B-115 | trnI | MN549218 |
| B-721 | trnI | MN549219 |
| B-302 | trnI | MN549220 |
| B-627 | trnI | MN549221 |
| B-720 | trnI | MN549222 |
| B-500 | trnI | MN549223 |
| B-1903 | trnI | MN549224 |
| B-626 | trnI | MN549225 |
| B-126 | trnI | MN549226 |
| B-625 | trnI | MN549227 |
| B-615 | trnI | MN549228 |
| B-601 | trnI | MN549229 |
| B-415 | trnI | MN549230 |
| B-639 | trnI | MN549231 |
| B-640 | trnI | MN549232 |
| B-649 | trnI | MN549233 |
| floden3847 | trnI | MN549234 |
| B-713a | trnI | MN549235 |
| B-651 | trnI | MN549236 |
| B-650 | trnI | MN549237 |
| B-004 | trnI | MN549238 |
| B-1812 | trnI | MN549239 |
| B-1811 | trnI | MN549240 |
| B-2001 | trnI | MN549241 |
| B-645 | trnI | MN549242 |
| B-644 | trnI | MN549243 |
| floden3971 | trnI | MN549244 |
| B-1801 | trnI | MN549245 |
| B-643 | trnI | MN549246 |
| B-647 | trnI | MN549247 |
| B-2101 | trnI | MN549248 |
| B-1604 | trnI | MN549249 |
| B-2801 | trnI | MN549250 |
| B-404 | trnI | MN549251 |
| B-301 | trnI | MN549252 |
| B-427 | trnI | MN549253 |
| B-125 | trnI | MN549254 |
| B-206 | trnI | MN549255 |
| B-1602 | trnI | MN549256 |
| B-711 | trnI | MN549257 |
| B-803 | trnI | MN549258 |
| B-408 | trnI | MN549259 |
| B-612 | trnI | MN549260 |
| B-723 | trnI | MN549261 |
| B-411 | trnI | MN549262 |
| B-610 | trnI | MN549263 |
| B-705 | trnI | MN549264 |
| B-502 | trnI | MN549265 |
| B-807 | trnI | MN549266 |
| B-714 | trnI | MN549267 |
| floden3834 | trnI | MN549268 |
| B-646 | trnI | MN549269 |
| B-701 | trnI | MN549270 |
| B-818 | trnI | MN549271 |
| B-603 | trnI | MN549272 |
| floden3835 | trnI | MN549273 |
| B-712a | trnI | MN549274 |
| B-815 | trnI | MN549275 |
| floden4073 | trnI | MN549276 |
| B-1603 | trnI | MN549277 |

**Support Table 4.**
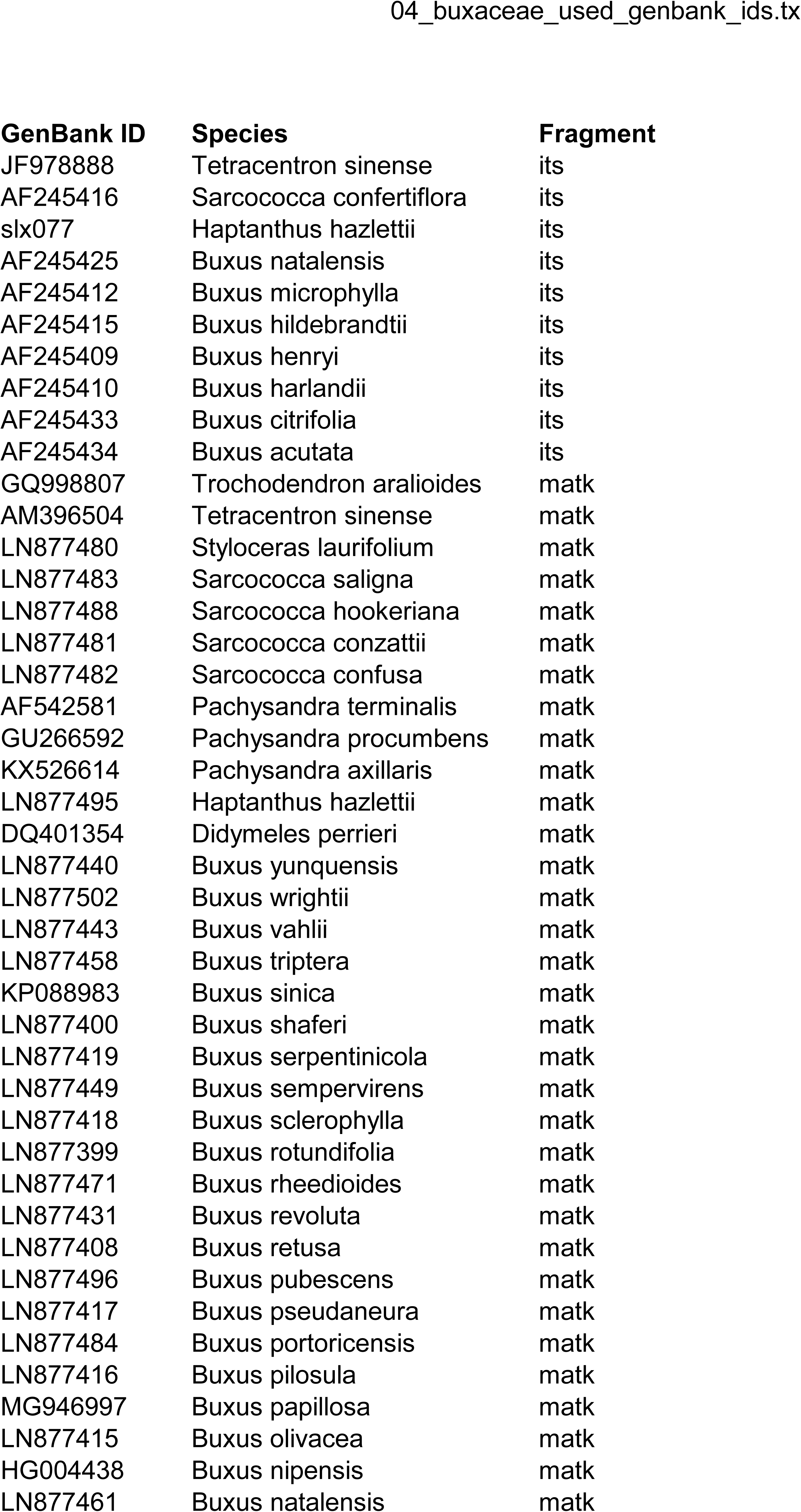
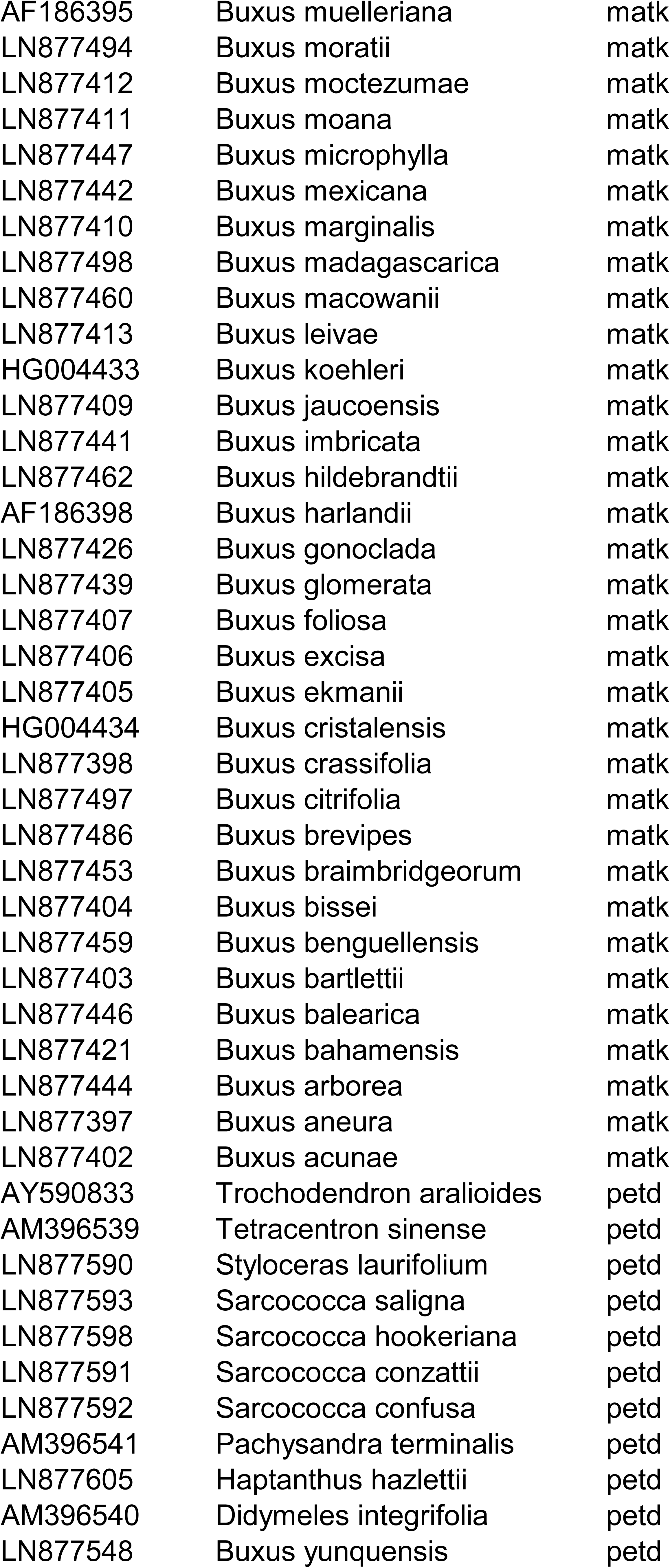
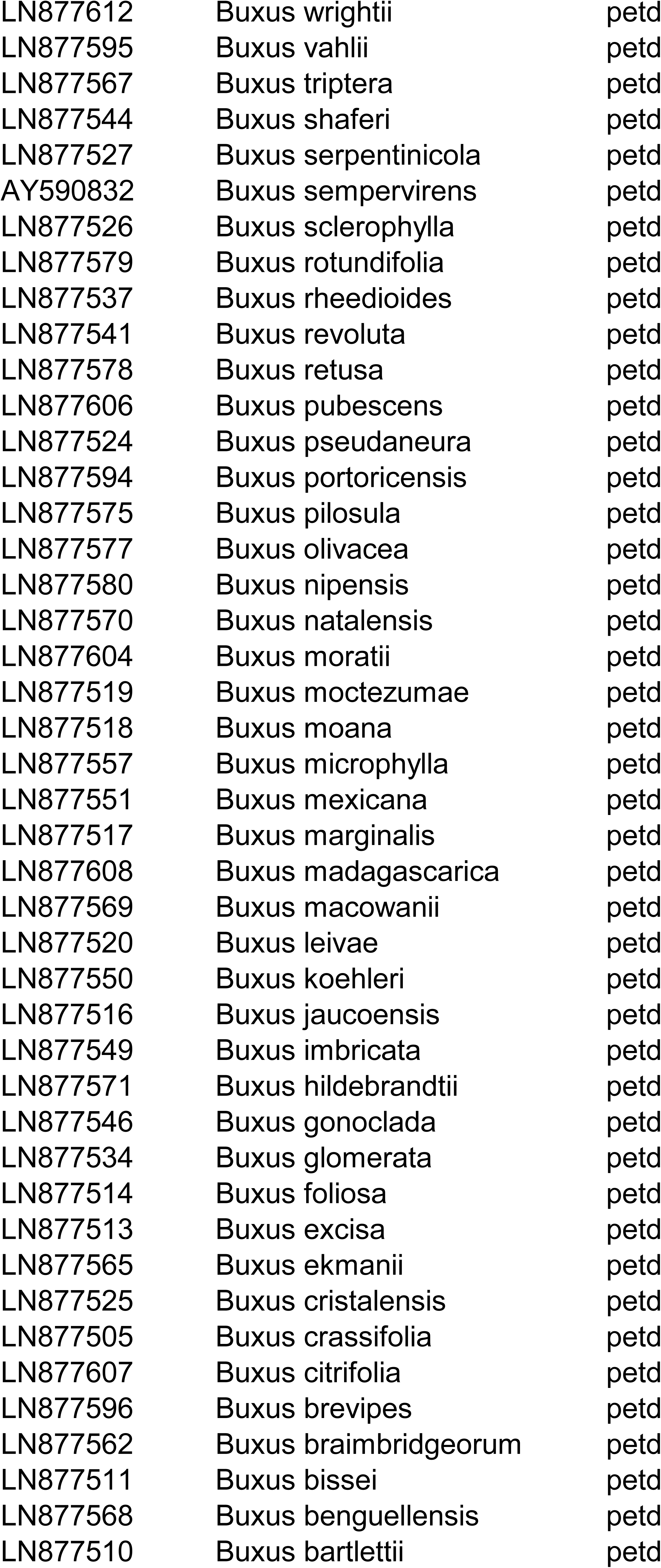
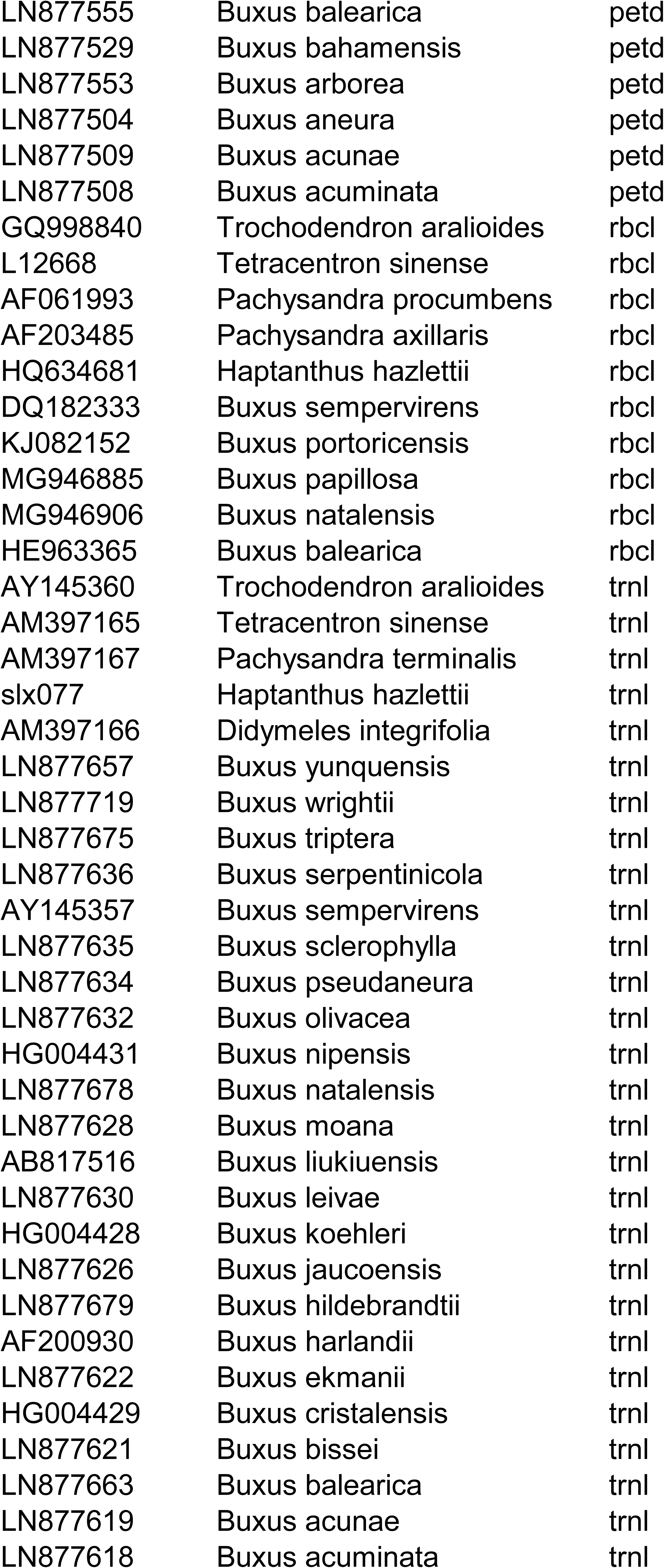
GenBank accession numbers of Buxaceae sequences of external origin used in this study.

