## Supplemental Figures and Tables for "Not out of the box: phylogeny of the broadly sampled Buxaceae": photographs_of_plants_sampled_in_the_field.pdf

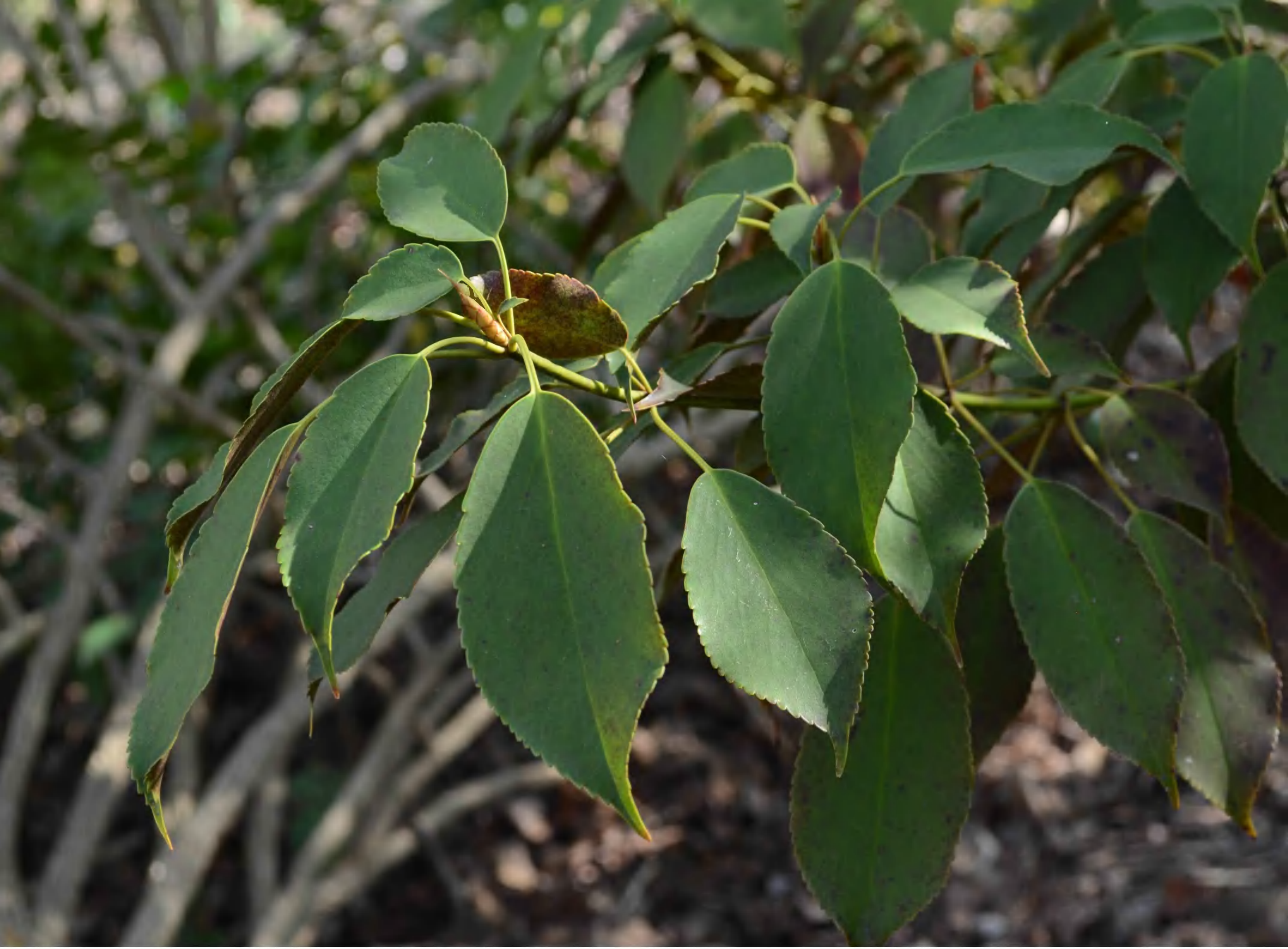

b_001_b_002_trochodendron_aralioides_dsc
_5730.jpg

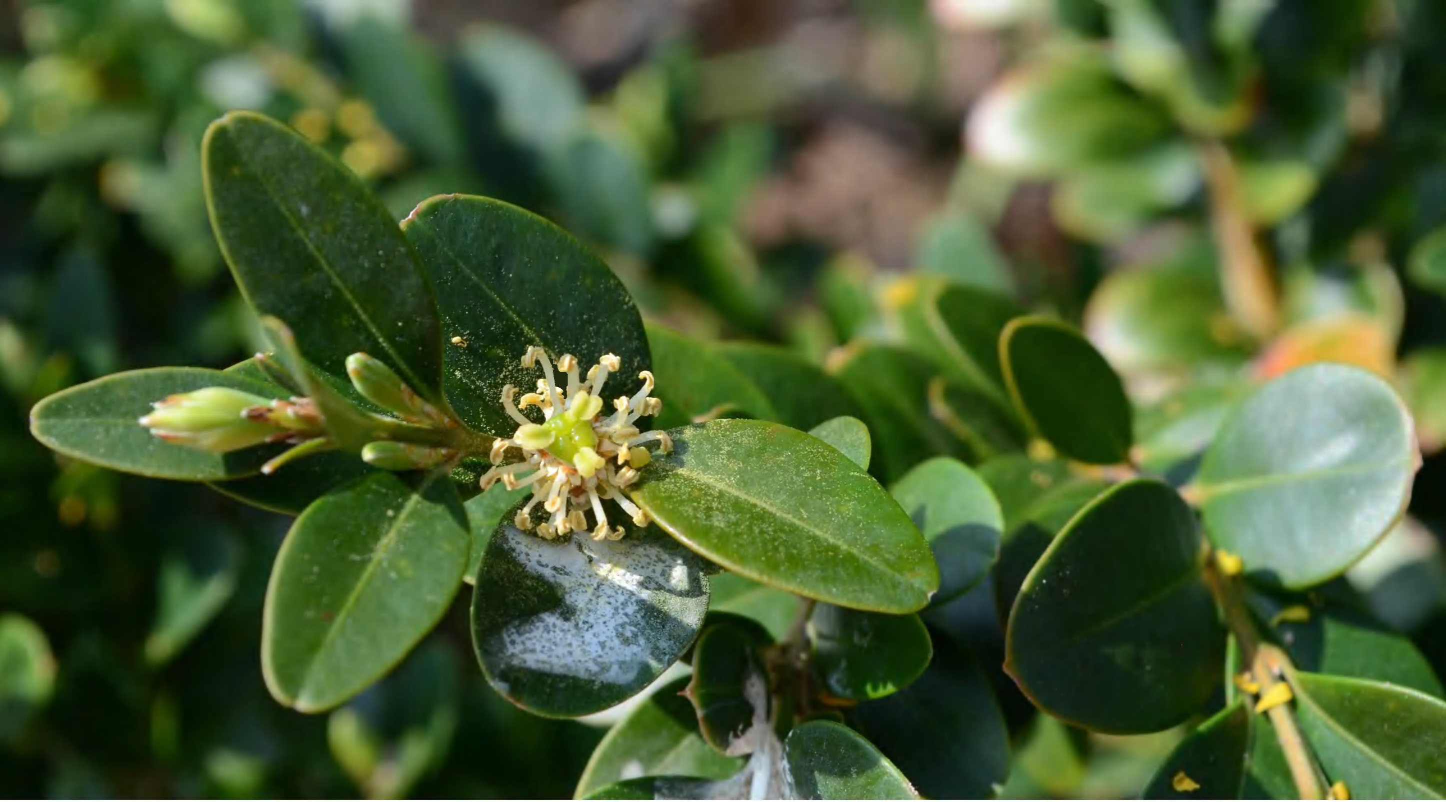

b_003_buxus_sempervirens_dsc_5708.jpg

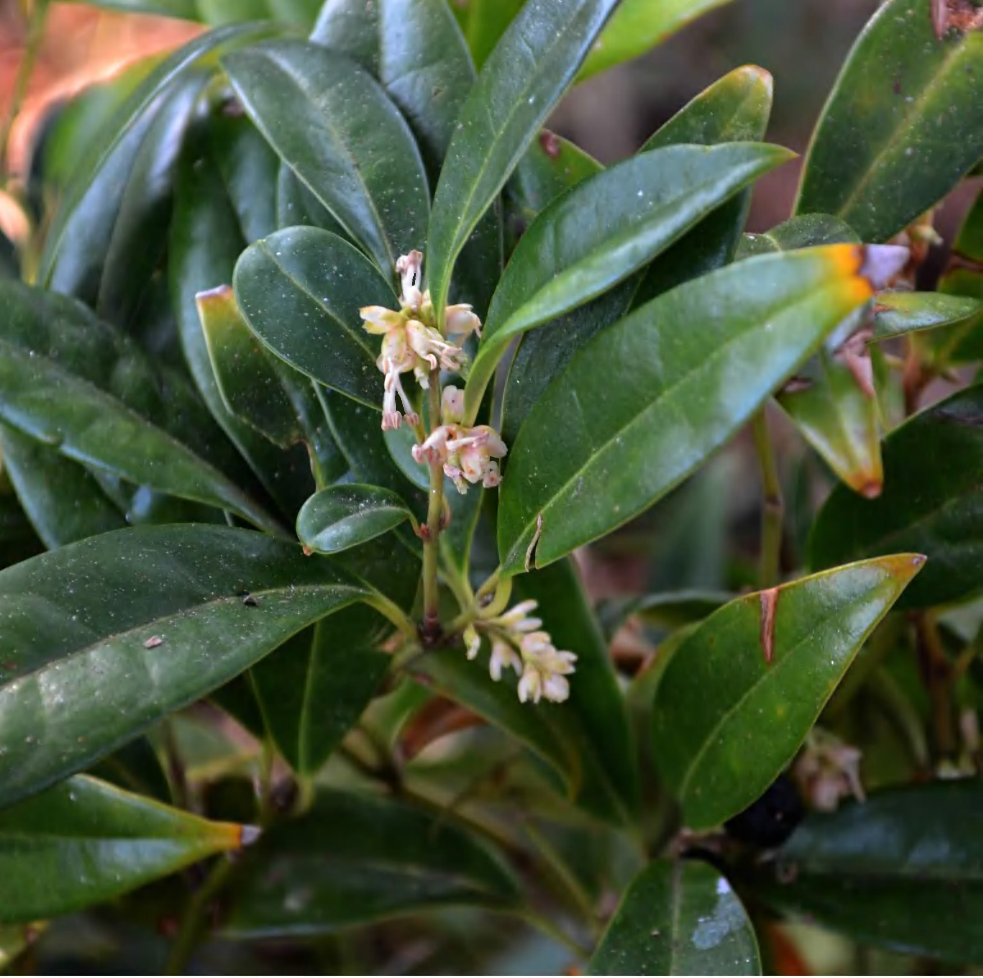

b_004_sarcococca_hookeriana_dsc_5703.jpg

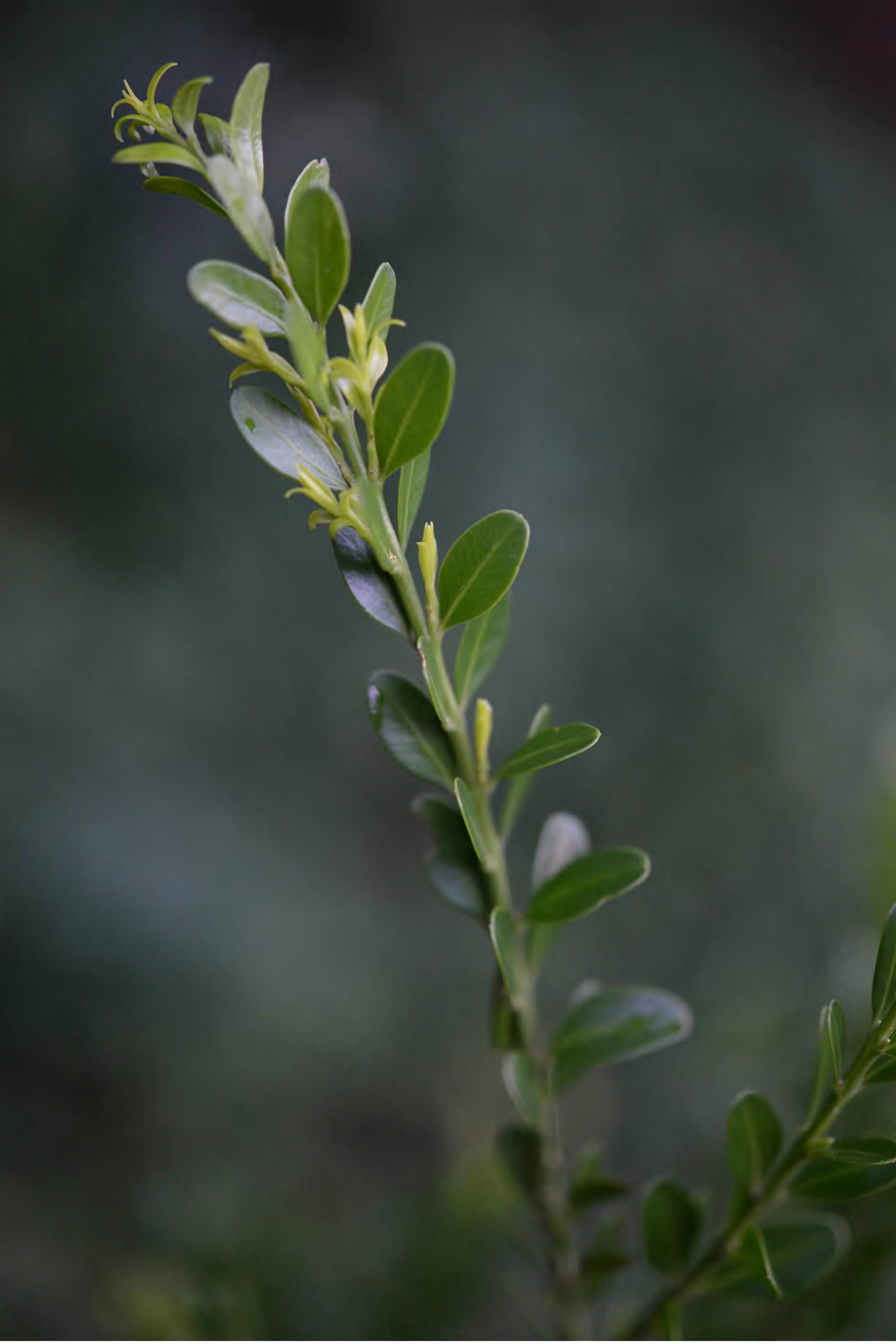

*b_207_buxus_macowanii_abs_5381.jpg*

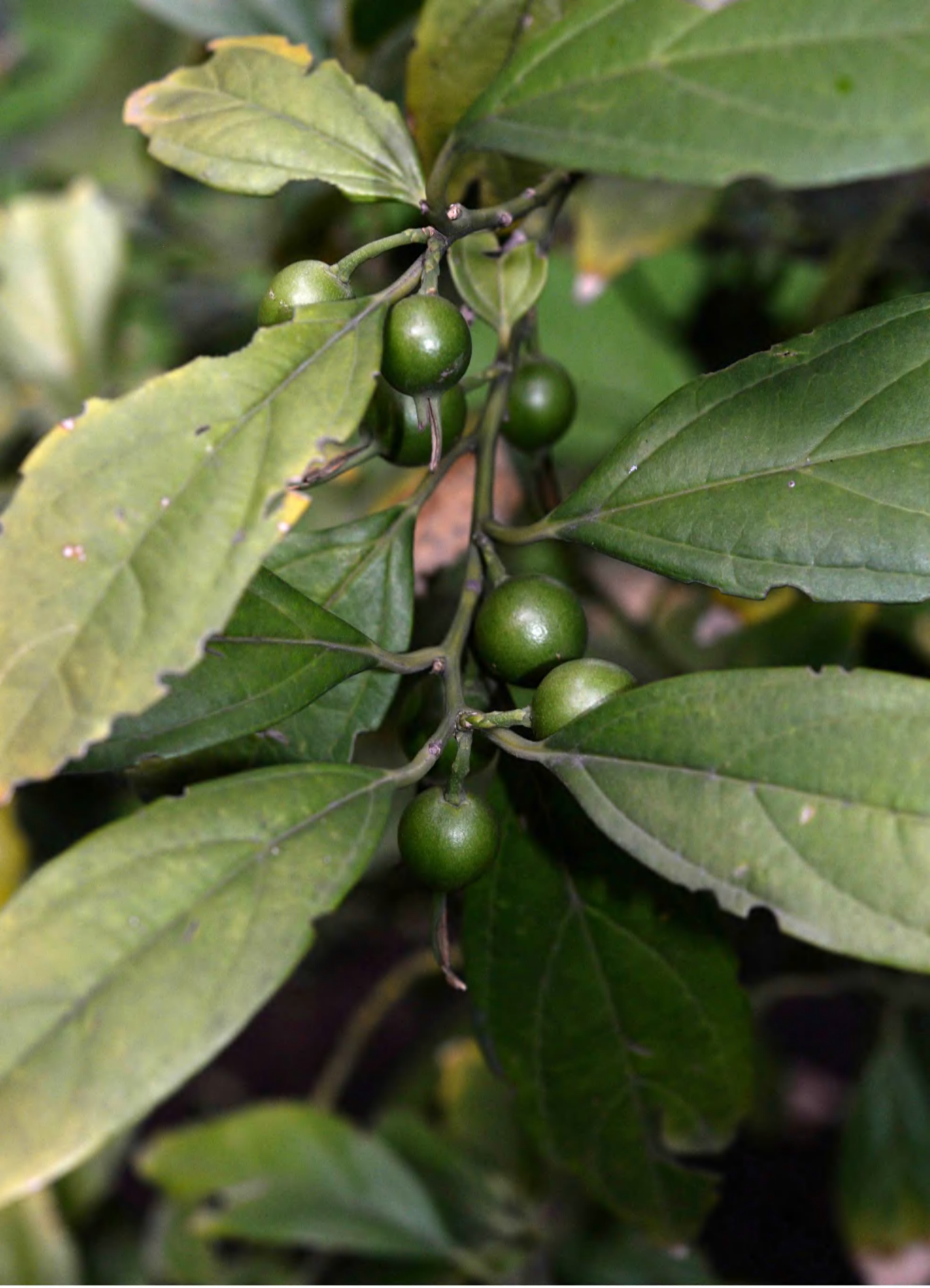

b_301_styloceras_laurifolium_abs_8795.jp
g

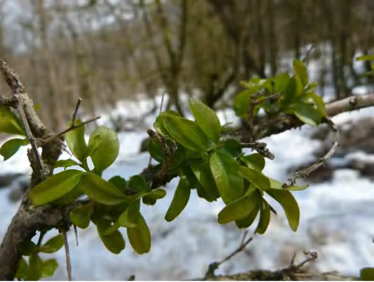

b_302_buxus_hyrcana_az_1.jpg

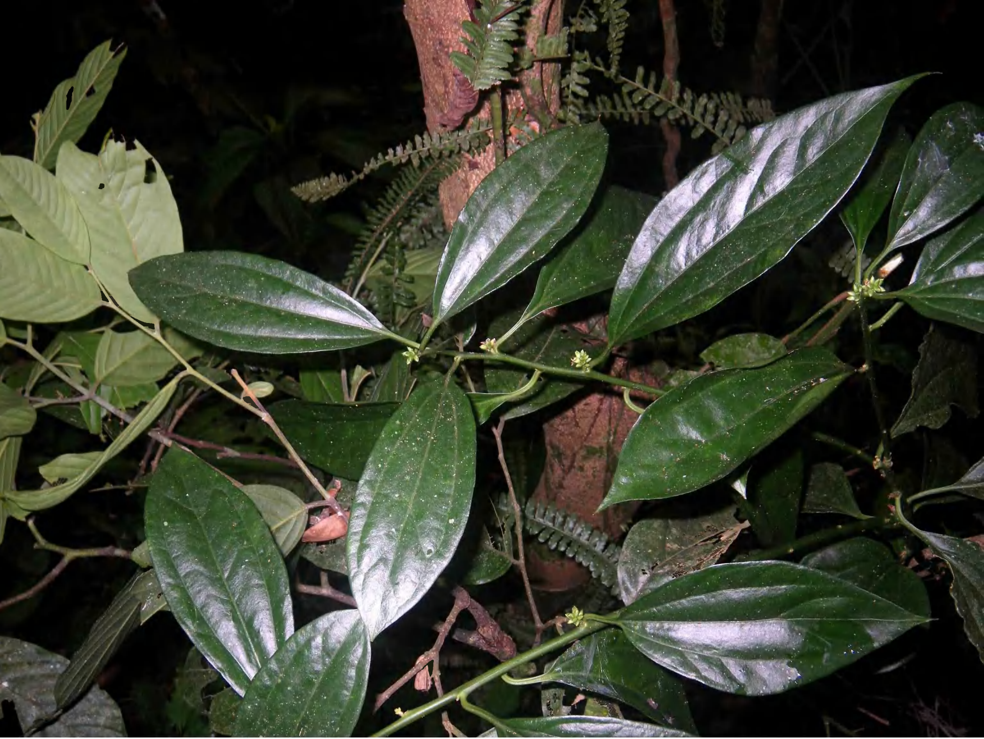

*b_303_sarcococca_balansae_imgp5650.jpg*
