## Supplementary figures and images for "Not out of the box: phylogeny of the broadly sampled Buxaceae"

### additional_trees.pdf

**cp rbcl + trnl + matk + petd MP**

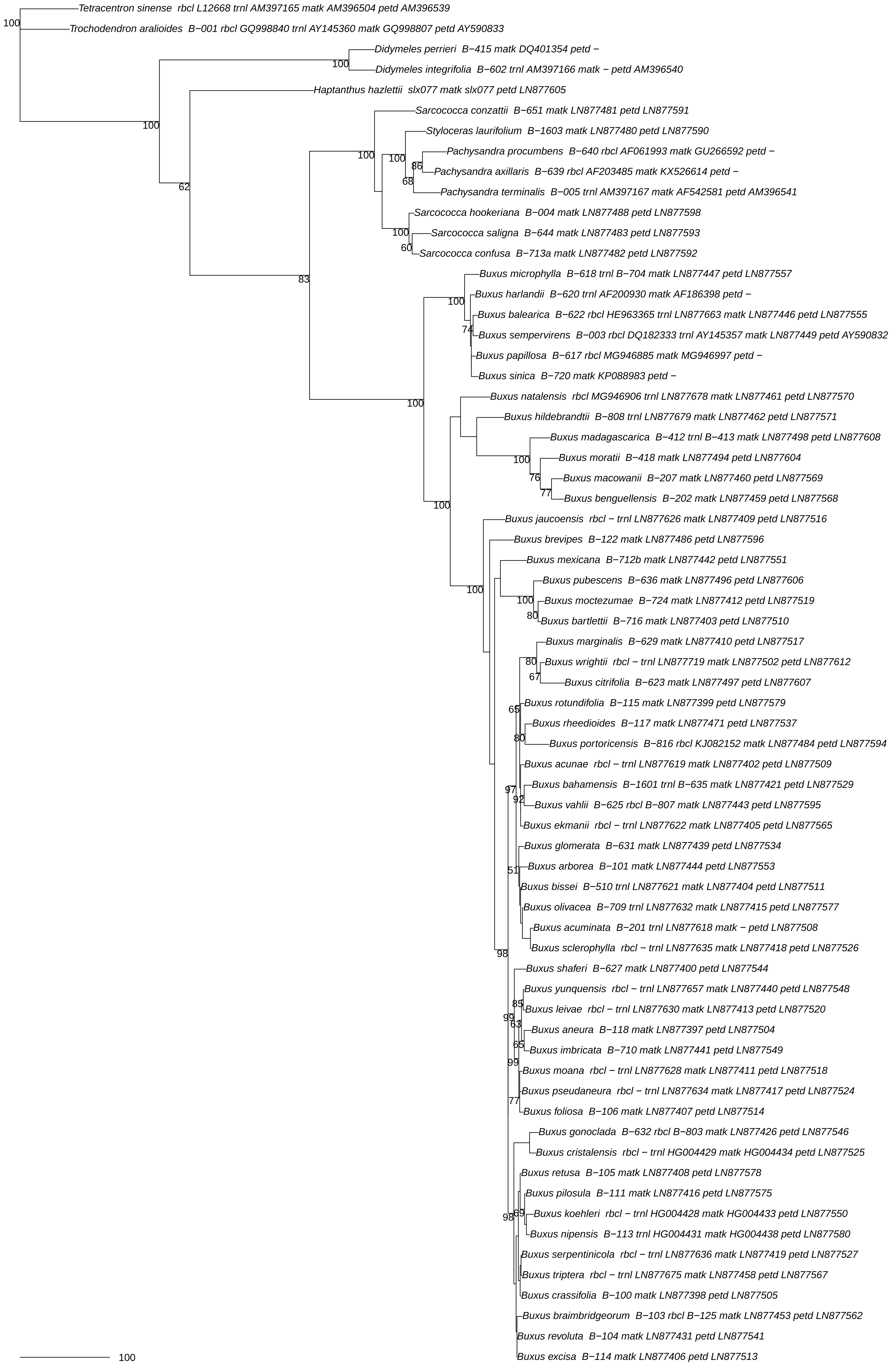

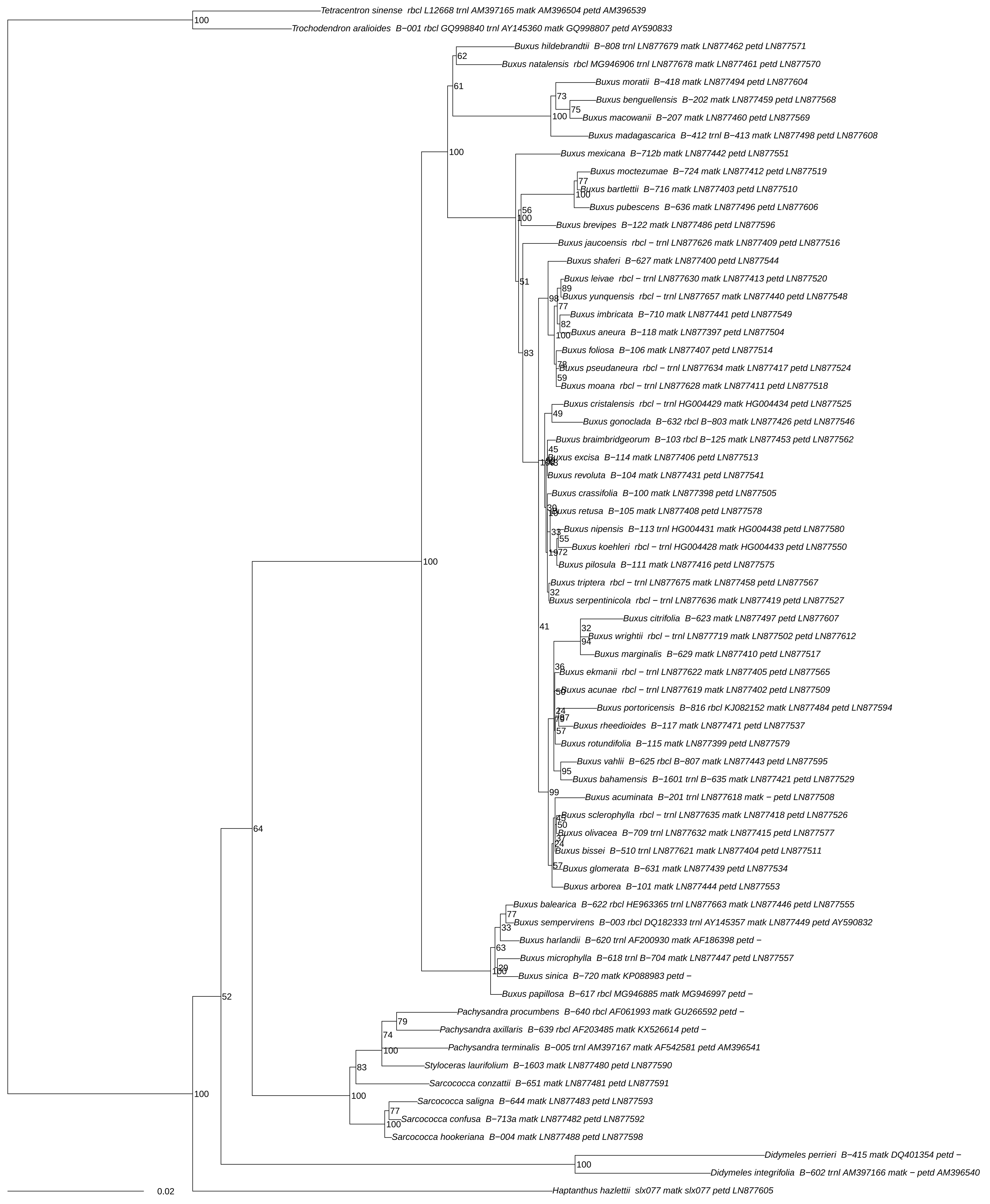

**oi its + rbcl + trnl MP**

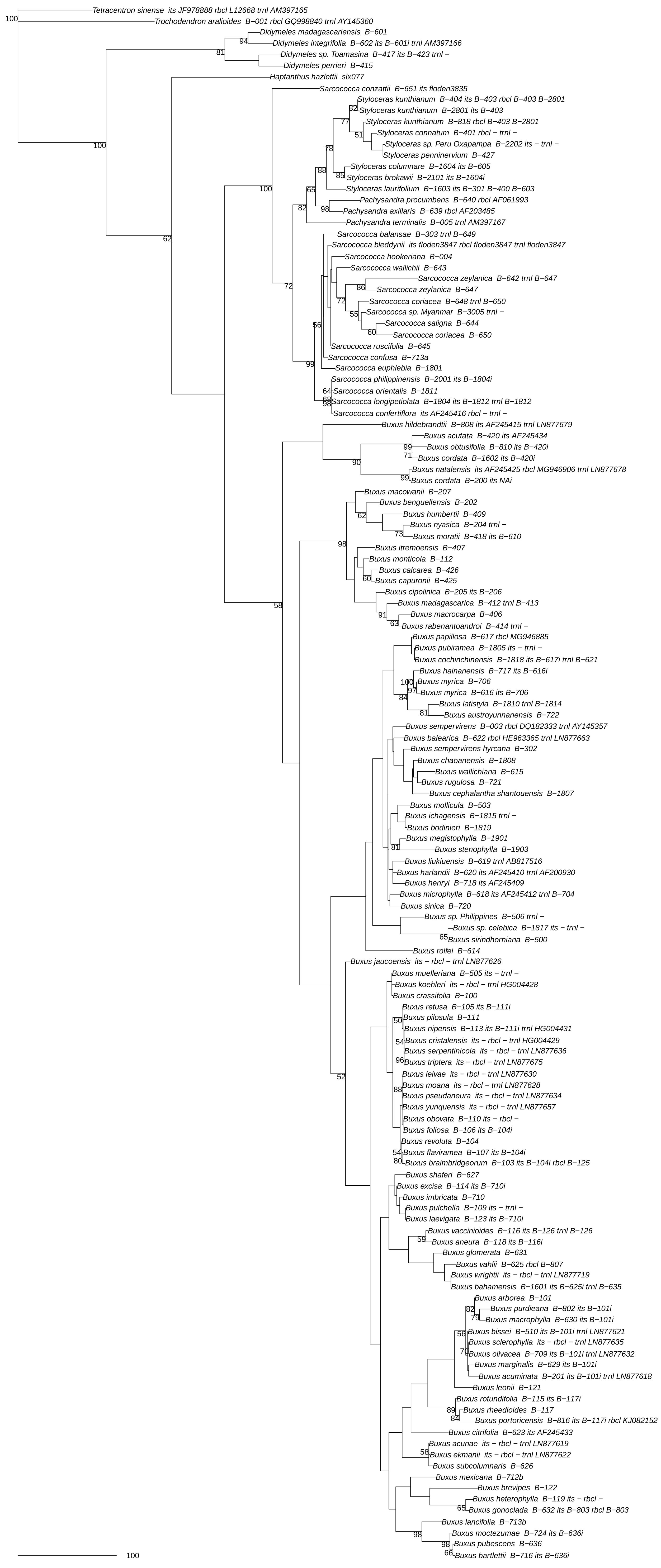

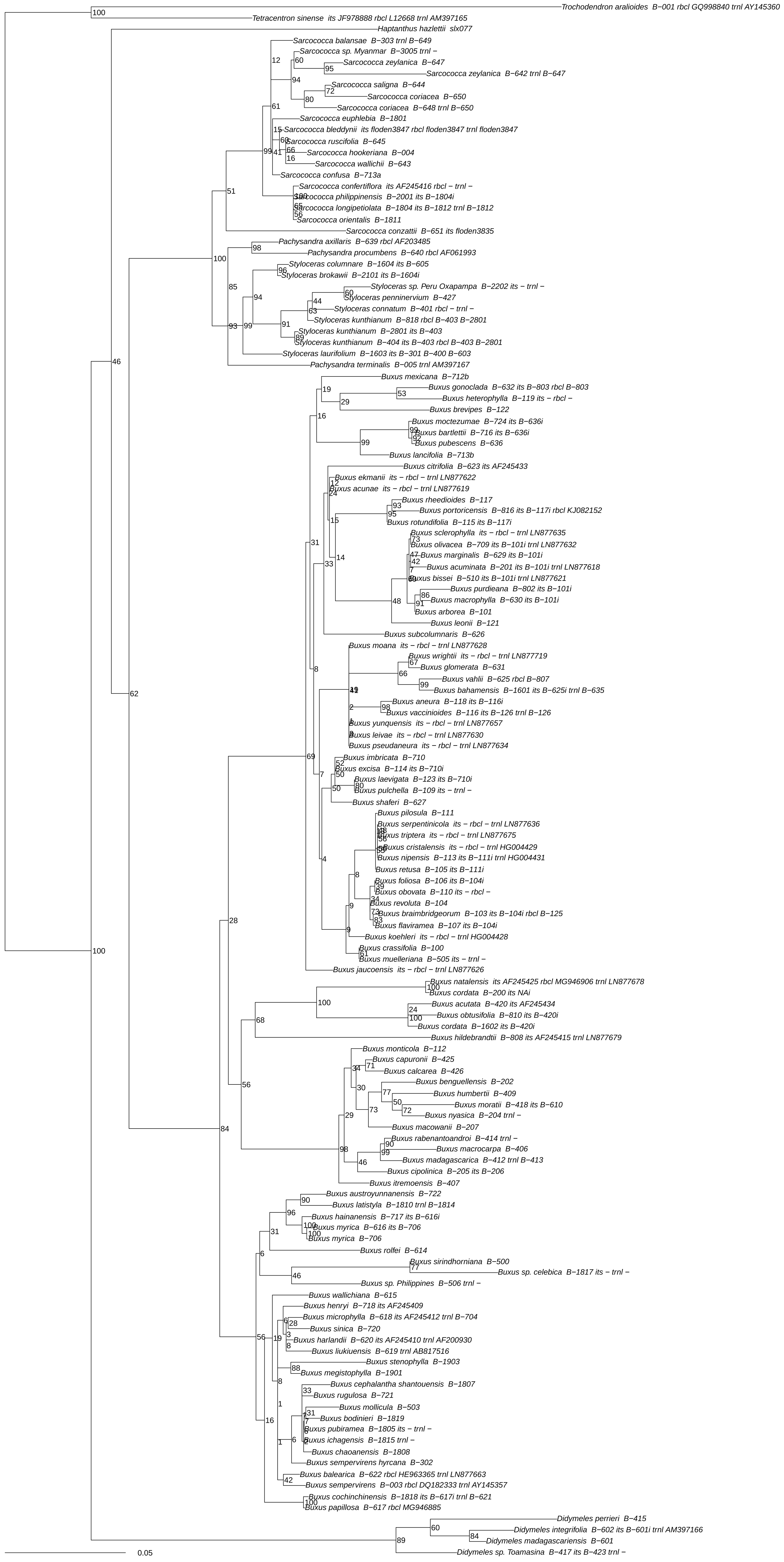
